# Beyond contacts: the important role of the support region to distinguish stable and transient protein interfaces

**DOI:** 10.1101/2025.01.29.635419

**Authors:** Tom Miclot, Stepan Timr

## Abstract

Protein-protein interactions (PPIs) are fundamental to virtually all cellular processes; however, elucidating the principles that govern protein association into complexes remains a significant challenge. In this study, we present a comparative analysis of stable and transient protein interfaces, offering a detailed perspective on the interactions that form them. Moving beyond the traditional focus on pairs of neighboring residues, we examine interacting pairs by identifying their distinct non-bonded interactions. Additionally, we account for the contextual dependence of these interactions by analyzing the regions of the interface where they occur. Our approach quantifies the diversity of pairs in each region, considering the type of interface. Furthermore, we introduce an innovative strategy to analyze pair co-occurrence, enabling a comparison of the inner local organization of stable and transient interfaces. Our findings reveal that stable and transient interfaces differ not in overall residue composition, but in the residue- and interaction-partitioning patterns across the variably hydrated regions of the interface. These results underscore the importance of considering the contextual environment in which pairs interact and identify the support region as a key determinant for distinguishing transient and stable protein complexes. The software package developed for this analysis is available as open source.

## 1 Introduction

Protein-protein interactions (PPIs) are physical associations between two or more protein molecules, resulting in the formation of higher-order structures that can involve distinct proteins or multiple copies of the same protein chain. The formation of these higher-order assemblies, held together by PPIs, is crucial for the function of many proteins, including structural proteins [1] and numerous enzymes [2, 3, 4, 5], as well as for various cellular processes [6, 7, 8, 9, 10].

The strength of PPIs is a key factor in determining the functional role of protein assemblies, and the nature of these interactions can be either transient or stable. For instance, PPIs involved in cell signaling pathways are typically transient, allowing for reversible protein association and repeated signal transduction [11]. In contrast, antibody-antigen complexes require high stability to efficiently neutralize pathogens [12]. Understanding the differences between transient and stable PPIs is a challenging task, particularly as the complexity of the system increases [13]. The stability of PPIs depends on various factors, including external conditions [14], the size of the interface [15], and the physico-chemical properties of the interface, such as residue composition [16, 17].

Within both stable and transient protein–protein interfaces, a complex network of non-bonded interactions between amino acid residues forms, playing a crucial role in protein assembly and complex stabilization [18, 19]. For example, recent work has highlighted the prevalence of electrostatic interactions in transient interfaces [20]. Given the importance of PPIs, a range of methods and software tools have been developed for their investigation, including approaches that focus on characterizing residue-residue interaction types using geometric descriptors [21], analyzing interaction networks [22], and employing machine learning to examine and validate the interfaces formed in protein complexes [23].

In this work, we characterize residue interactions within the various regions that compose an interface. We introduce the Molecular Interaction anaLysis toOl-kiT (MICLOT), a robust Python package capable of identifying a wide variety of non-bonded interaction types and their sub-types. By applying this tool to curated benchmark collections of high-quality structures, comprising both stable and transient complexes, we identify the primary differences in interaction patterns and their context-dependent properties within the interface. This comparative analysis reveals the unique characteristics of stable and transient interfaces, shedding light on the underlying mechanisms that govern their behavior.

## 2 Results and Discussion

Our study utilizes a comprehensive dataset comprising 413 protein hetero-complexes, compiled from multiple databases and filtered by UniprotKB accession number to eliminate redundancy. To ensure data quality, only structures with a resolution of 2.5 Å or better were retained. The dataset is categorically divided into three types of complexes based on their binding affinity: stable, transient, and intermediate. Further details regarding the dataset and methodology can be found in the Methods section and Supplementary Information. We initiate our analysis by examining the size and composition of the protein-protein interfaces, thereby validating the efficacy of our MICLOT software and the consistency of the dataset. Subsequently, we investigate the interactions between residues at the interface. To facilitate precise analysis, the interfaces are subdivided into distinct regions, adopting the nomenclature proposed by Levy [24] and Kastritis et al. [25], which delineates the Core as the solvent-inaccessible region, the Rim as the most hydrated area, and the Support as a semi-hydrated region.

### 2.1 Interface size and global composition are not critical descriptors for stability

Previous work reported a correlation between the size of the buried surface and the binding affinity [26], as well as the observation that transient PPI tend to be associated with smaller interfaces [27]. We reexamined this relationship using our dataset, featuring a broad range of interface areas, and found no strong correlation between binding affinity and area both when considering the entire interface and when evaluating the areas of the Core & Support and Rim regions separately, as identified by our software (see Figure 1). It is possible that crystal packing [28, 29, 30] effects and/or extreme value points influenced our results slightly. However, our data are in line with the work of Chen, Sawyer, and Regan [26] (Figure 1A of the study), as they also show an overlap between the interface sizes of transient and stable complexes. Therefore, interface size is not sufficient to clearly distinguish between stable and transient complexes from our dataset.

**Figure 1:**
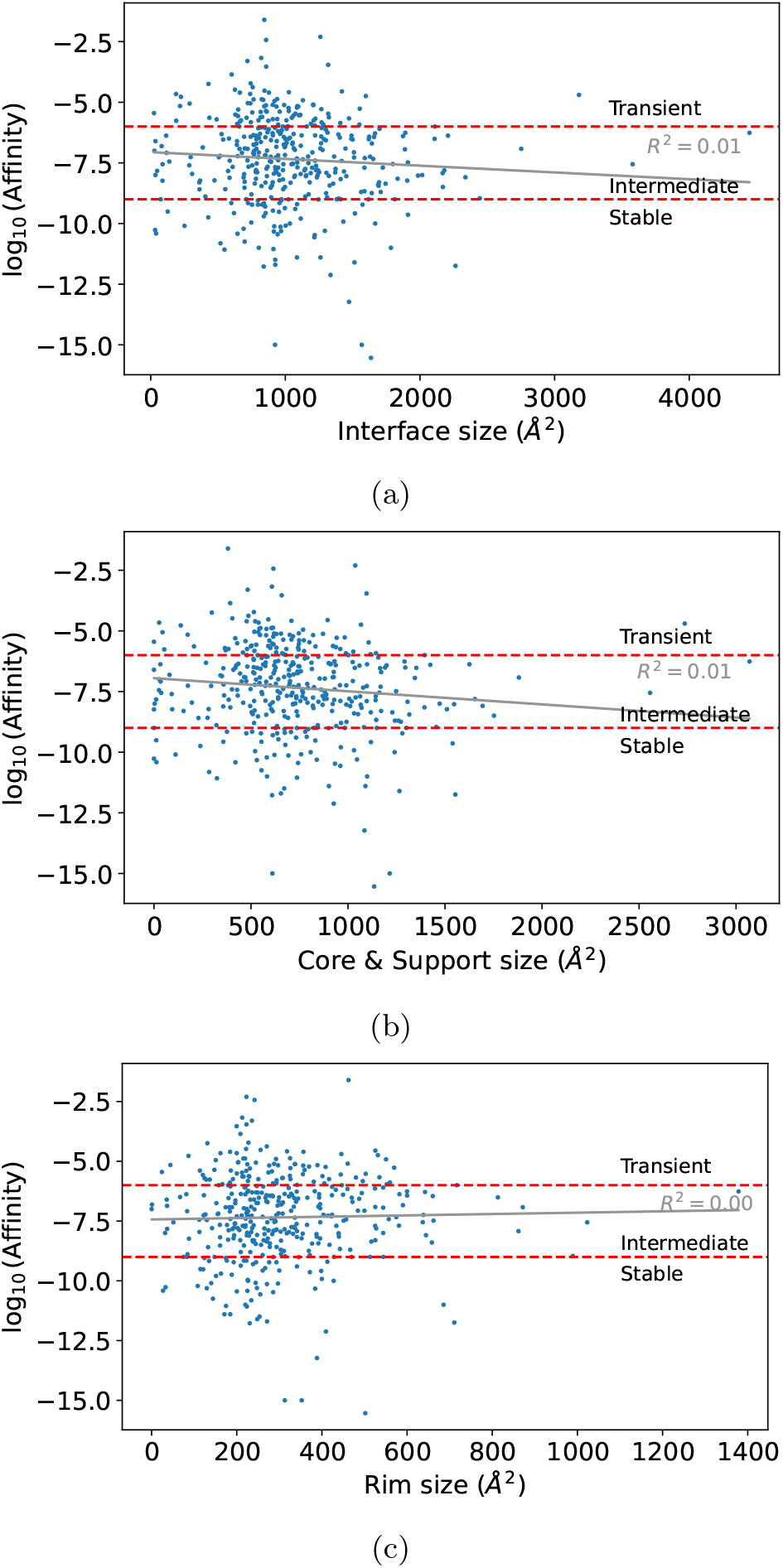
(a) There is no significant correlation between affinity and the size of the interface, nor with that of (b) Core & Support or (c) Rim. The regression lines were obtained using a linear regression tool provided by scikit-learn [31].

Previous work also found no obvious link between the global chemical composition of the interface and complex stability [26]. To characterize the global residue content, we evaluated the overall residue frequencies, as was done before [15]. Our data confirm that the global residue compositions of stable and transient interfaces are similar, with transient interfaces containing an increased population of leucine and, at the same time, decreased populations of glycine and aromatic residues such as tyrosine and tryptophan (Figure 2).

**Figure 2:**
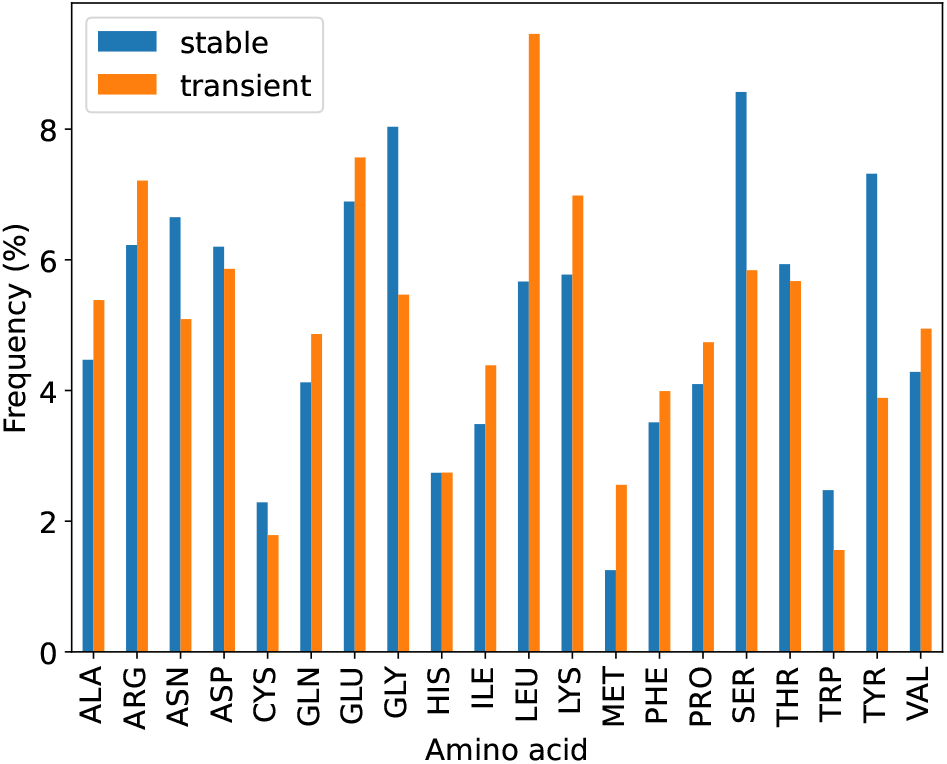
Frequencies of residues populating stable and transient interfaces.

### 2.2 Interactions differ across interface regions

Non-bonded interactions between amino acid residues are critical for stabilizing protein–protein interfaces and mediating their specificity. Consequently, a detailed characterization of these interfaces, particularly from the perspective of the interactions within them, is essential to understanding the factors that govern their stability. Our analysis extends beyond merely evaluating residue proximity or side-chain contacts. Instead, it enables the detection of various non-bonded interaction types and an assessment of their context dependence.

Our findings reveal that the propensity of a residue pair to engage in a specific type of interaction can vary depending on its location within the interface. For instance, an interacting ARG-ASP pair forms a salt bridge in 72% of its occurrences in the Core, 77% in the Support, and only 61% in the Rim (Table 1). More broadly, residue pairs are more likely to form favorable interactions in the Support region compared to Rim (see Figure S6). Conversely, potentially unfavorable interactions are more frequently observed in the Rim (see Figure 3).

**Table 1:** The probability of forming a salt bridge for amino-acid residue pairs in distinct regions.

| Pair | All | Protein | Core | Support | Rim |
| --- | --- | --- | --- | --- | --- |
| ARG-ASP | 0.73 | 0.75 | 0.72 | 0.77 | 0.61 |
| ARG-GLU | 0.70 | 0.73 | 0.67 | 0.71 | 0.62 |
| ASP-LYS | 0.52 | 0.45 | 0.65 | 0.67 | 0.61 |
| GLU-LYS | 0.56 | 0.60 | 0.53 | 0.60 | 0.47 |

**Figure 3:**
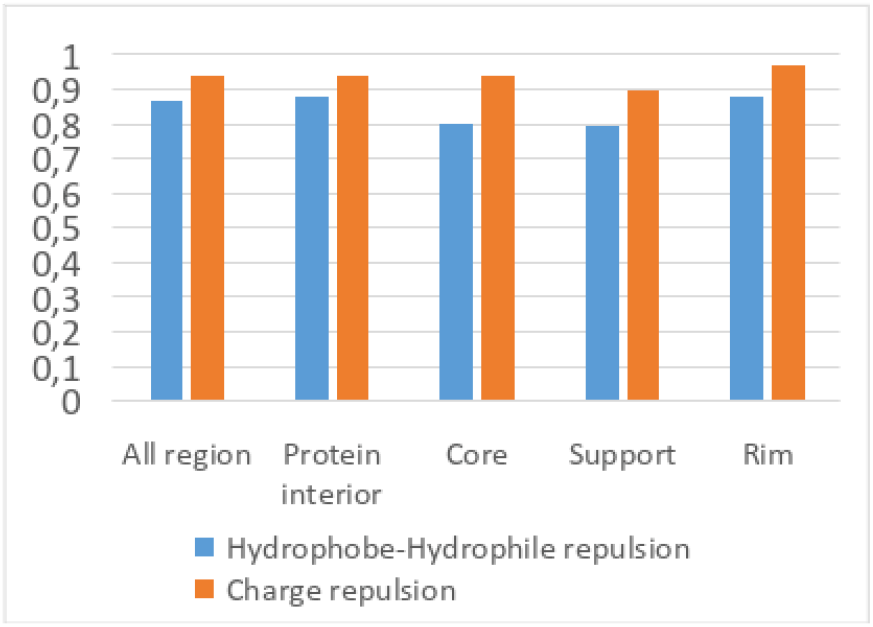
Overall trend of charge repulsion and hydrophilic-hydrophobic repulsion. The values represent the probability of the respective interaction for residue pairs that can engage in it.

Overall, we find that with the exception of salt bridges and charge repulsions, the total counts of interactions detected in Rim are smaller compared to Support (see Table S2). Therefore, despite the larger proportion of Rim in terms of its surface area (see Table 2), it hosts fewer interactions stabilizing the interface than Support.

**Table 2:** Mean areas of the whole interface and individual interface regions for stable, intermediate, and transient complexes. *The complete table is shown in SI subsection D*.*2*.

| Size |  |  |  |  |
| --- | --- | --- | --- | --- |
|  | Interface<br>(Å<sup>2</sup>) | Core<br>(Å<sup>2</sup>) | Support<br>(Å<sup>2</sup>) | Rim<br>(Å<sup>2</sup>) |
| Transient | 933.7 | 554.8 | 90.3 | 288.6 |
| Intermediate | 1092.1 | 664.4 | 106.5 | 321.2 |
| Stable | 989.9 | 607.2 | 123.5 | 259.2 |

| Ratio |  |  |  |
| --- | --- | --- | --- |
|  | Core<br>(%) | Support<br>(%) | Rim<br>(%) |
| Transient | 56.5 | 10.0 | 33.5 |
| Intermediate | 58.4 | 10.0 | 31.6 |
| Stable | 58.6 | 11.5 | 29.9 |

These observations suggest that the local context within an interface significantly influences residue pair interactions. Thus, partitioning the interface into Core, Support, and Rim regions may be a key factor governing its overall stability.

### 2.3 Core and Support are more prominent in stable interfaces

To compare the sizes of Core, Support, and Rim regions in our dataset, we calculated the mean areas of the whole interfaces and the individual regions. With the exception of Support, the mean areas show no clear trend across transient, intermediate, and stable complexes (Table 2). However, the relative proportions of each interface region do reveal noticeable trends in mean, although the standard deviation shows an overlap between interface sizes (Table S1). The Core region increases its relative size as the interfaces become more stable, expanding from 56.5% to 58.6% of the total interface area, an inverse relationship is found for the rim, shrinking from 33.5% down to 30% of the total interface area. The support region, having a comparable relative size in transient and intermediate complexes, becomes more pronounced in stable complexes. Overall, the results suggest changes in the partitioning of an interface into less hydrated (i.e., Core and Support) and more hydrated (i.e., Rim) regions when moving from stable to transient complexes. This observation is in line with the findings of a previous study (Luo et al. [27]), which showed that non-specific complexes contain more residues in Rim compared to specific complexes.

A detailed look at the physico-chemical composition of individual regions reveals pronounced differences between Core and Support on the one hand, and Rim on the other (see Figure S7 in SI). Using the residue classification introduced by Pommié et al. [32] we find that, as expected, Rim contains more charged residues than Core and Support, as well as more polar and hydrophilic residues. At the same time, it contains significantly fewer aromatic and aliphatic residues. These features are equally true for both stable and transient interfaces.

Compared with stable interfaces, the Rim region of transient interfaces is even more enriched in charged residues. However, Core and Support contain more apolar and hydrophobic residues (see Figure S7 in SI) in transient interfaces than in stable interfaces. This seemingly counterintuitive trend can be partially explained by an increased content of aliphatic residues in Core and Support of transient interfaces at the expense of aromatic residues such as tyrosine, which is classified as a polar residue with neutral hydropathy. In addition, Core and Support of transient interfaces contain fewer hydrogen bonding residues, which might be an additional factor that limits the stability of these complexes.

In many aspects, the composition of Support is more Rim-like in stable complexes than in transient complexes (see Figure S7 in SI). This further strengthens the idea that the distinction between stable and transient interfaces lies not in the overall residue composition, but in the context in which these residues exist. This idea is supported by examining the partitioning of the various residue classes across the different interface regions (Figure 4). This analysis is analogous to that performed by Levy [24]; however, it is based on residue classes rather than residue names. The analysis shows that stable and transient interfaces differ in the partitioning of a number of residue classes between Support and Rim. As an example, positively and negatively charged residues are more concentrated in the Support region of stable interfaces compared to transient interfaces, where they predominantly populate Rim. In contrast, the partitioning of residue classes into Core and Rim (NIS) does not exhibit any major differences between stable and transient interfaces.

**Figure 4:**
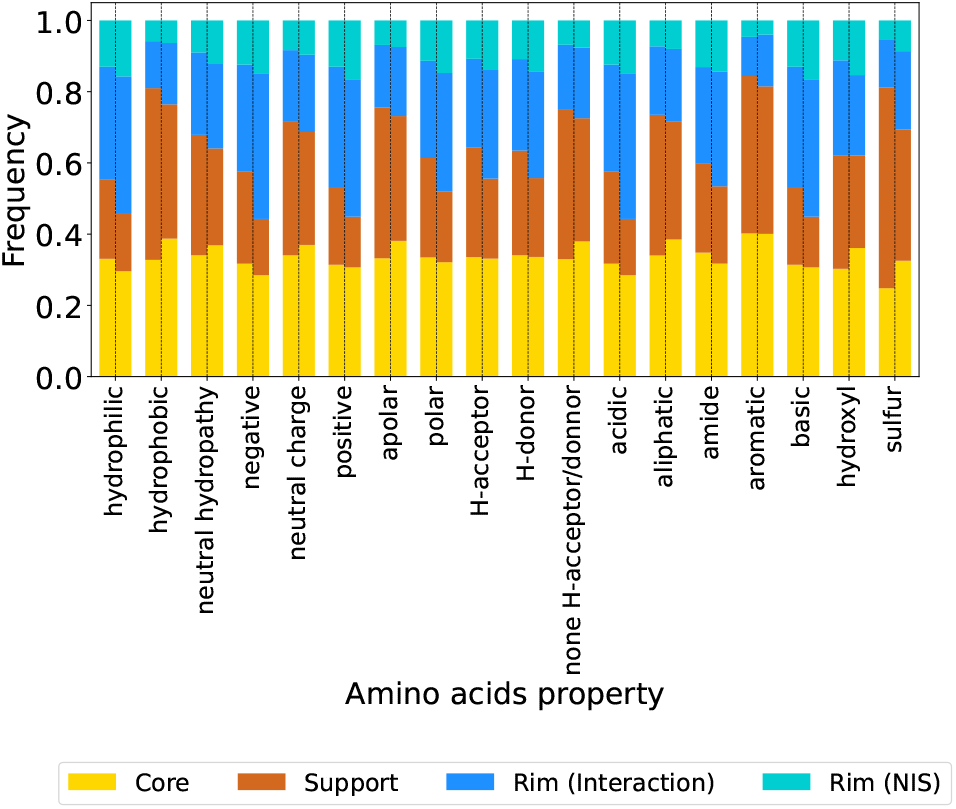
Stable and transient interfaces show differences in the partitioning of residue classes between Support and Rim (interaction). *Stable interfaces are shown on the left and transient interfaces on the right*.

The interplay between Support and Rim observed in stable and transient interfaces makes these regions promising candidates for interface type evaluation. Therefore, analyzing their physico-chemical properties and the interactions arising in these regions can be of great importance when trying to distinguish between stable and transient interfaces.

### 2.4 Pair class partitioning highlights the role of Support in stable interfaces

The interaction analysis allows us to evaluate the local structure of interfaces in terms of interacting pair types. Note that compared with previous approaches, our analysis does not rely on an arbitrary distance threshold for residue pair qualification but instead uses precise geometric criteria to determine whether two neighboring residues actually form an interacting pair. The interacting pair classes (see SI section I) are based on the residue classes defined by Pommié et al. [32] and used above.

The relative partitioning of interacting pair classes (see subsection I.1) reflects the Support–Rim interplay which we reported for the residue class composition. As a nearly universal trend across all the properties (i.e., charge, hydropathy, polarity, H-bonding ability, and chemical category), a larger proportion of interacting pairs form the Support region in stable interfaces than in transient interfaces. Pairs involving amide-containing side chains are an exception to this trend (see Figure S13).

In some cases, the relative partitioning simply varies between Support and Rim, without affecting the relative proportion of pairs belonging to Core, such as for the hydrophilic–neutral hydropathy pair (see Figure S10). In other cases, changes in the Core partitioning are also observed. These changes can go in both directions: On the one hand, the relative proportion of charged-charged pairs forming Core increases in stable interfaces, as do the proportions of hydrophilic–hydrophilic and polar–polar pairs. On the other hand, the hydrophobic–hydrophobic pairs are more concentrated to Support rather than Core in stable interfaces, whereas the opposite is true in transient interfaces. Similarly, the partitioning of pairs involving aromatic residues, sulfur, or amide groups exhibits a shift toward the Support in stable interfaces.

Overall, the results point to differences in the inner organization of stable and transient interfaces and highlight the important role of the Support region in stable interfaces. These findings underscore the importance of considering the protein regions when analyzing PPI.

### 2.5 Increased pair class diversity toward Rim

As expected, larger interfaces tend to harbor a greater number of pairs and interactions. The distributions of interaction counts and pair counts in protein–protein interfaces (see Figure S14 and Figure S4) have a shape similar to the interface size distribution. Indeed, we observe that these counts scale linearly with interface size (see SI section K). In contrast, the numbers of interaction types and pair types exhibit a weaker dependence on the interface size. In fact, their distributions are bell-shaped, with interfaces typically comprising between 6 and 13 interaction types and between 40 and 95 pair types. These findings imply that a moderate level of diversity in interaction and pair types is essential for effective protein complex formation.

The analysis of the Shannon entropy and evenness indices, as described in SI section L and presented in Table 3, reveals distinct patterns in the diversity of pair populations across different interface regions. In particular, Rim exhibits high evenness values, indicating a highly diverse pair population. The evenness values for Core and Support, evaluated together to maximize statistics, are smaller, but they still exceed those for the protein interior, which exhibits a relatively uniform pair composition. The quantification of heterogeneity using Shannon entropy and evenness indices also provides a more nuanced understanding of the differences between stable and transient interfaces. Namely, Core and Support of stable interfaces are found to be slightly less homogeneous than those of transient interfaces, which may be attributed to the need for more diverse pair types to ensure stable binding. In contrast, the Rim regions of both stable and transient interfaces exhibit a similar level of heterogeneity.

**Table 3:** Pair diversity evaluation for different protein regions of transient and stable interfaces. *H*_*m*_a_*x*_ is equal to 3.322 for all protein regions. C & S stands for the Core and Support regions, which are considered together here.

| Protein region | Entropy H (bits) | $\Delta H$ (bits) | Evenness index J |
| --- | --- | --- | --- |
| Interior | 2.254 | 1.068 | 0.679 |
| C & S *Stable* | 2.858 | 0.464 | 0.860 |
| C & S *Transient* | 2.759 | 0.563 | 0.830 |
| Rim *Stable* | 3.118 | 0.204 | 0.939 |
| Rim *Transient* | 3.168 | 0.154 | 0.954 |

### 2.6 Local organization of Core and Support resembles protein interior in stable interfaces

So far, our analysis has allowed us to evaluate the partitioning residues, interaction- and pair types, and their diversity in distinct interface regions. Here we examine whether certain residue types are likely to co-occur and form an interacting pair, and to what extent the pair type population is simply the product of individual frequencies of the residue types involved. To perform this analysis, which provides insight into the local interface architecture, we use mathematical tools inspired by natural language processing, namely positive point-wise mutual information (PPMI) [33, 34, 35] and term frequency-inverse document frequency (tf-idf) [36, 37]. The methodological approach is detailed in SI section M. First, we use the tf-idf values to filter out pair types that are only present in a limited subset of structures for the given interface region. The PPMI values of the remaining pair types display marked trends (see Table 4). Note that the PPMI score measures the co-occurrence of two residue types engaging in an interaction. Thus, a high PPMI values can be achieved although the frequency of the pair type is low in a given region, and vice versa. For example, the negative-positive pair type exhibits a high PPMI value in the protein interior. This shows that oppositely charged residues, which tend to be scarce in the protein interior, are likely to form a pair whenever they are present. In contrast, we find a zero PPMI value for the apolar-apolar pair type in the same environment, which is largely formed by apolar residues; however, lacking any strong mutual correlation.

**Table 4:** Co-occurency PPMI score, after data filtering based on tf-idf values lower than 1 (see SI section M). Removed values are replaced by a point (.). C & S stands for the Core and Support regions, which are considered together here.

|  | **Interior** | **C & S**<br>**transient** | **C & S**<br>**stable** |
| --- | --- | --- | --- |
| apolar-apolar | 0.000 | 0.743 | 0.292 |
| apolar-negative | 1.553 | 0.986 | 1.213 |
| apolar-polar | 0.924 | 1.270 | 1.191 |
| apolar-positive | 1.411 | 1.137 | 1.099 |
| negative-negative | . | . | . |
| negative-polar | 1.433 | 0.143 | 0.864 |
| negative-positive | 1.563 | . | 0.629 |
| polar-polar | 0.000 | 0.000 | 0.000 |
| polar-positive | 1.029 | 0.000 | 0.421 |
| positive-positive | 1.338 | 0.064 | 0.246 |

Table 4: Co-occurrence PPMI score, after data filtering based on tf-idf values lower than 1 (see SI section M). Removed values are replaced by a point (.). C & S stands for the Core and Support regions, which are considered together here.
|  | **Rim**<br>**transient** | **Rim**<br>**stable** |
| --- | --- | --- |
| apolar-apolar | . | . |
| apolar-negative | . | . |
| apolar-polar | 0.583 | 0.626 |
| apolar-positive | 1.007 | 1.303 |
| negative-negative | . | . |
| negative-polar | 1.285 | . |
| negative-positive | . | . |
| polar-polar | 0.000 | 0.000 |
| polar-positive | 0.706 | 0.761 |
| positive-positive | 2.163 | 2.030 |

Compared with the protein interior, the core & support region tends to exhibit lower PPMI values for pair types involving charged residues and higher PPMI values for residues involving apolar residues. Importantly, with the exception of the apolar-positive pair type, the core & support of stable interfaces features more interior-like PPMI values than that of transient interfaces. This indicates that stable interfaces can be distinguished from transient ones by the greater similarity of the core & support region to the protein interior in terms of their inner organization.

In contrast, the rim region does not show any major differences in PPMI values between stable and transient interfaces. In both subsets, the positive-positive pair type exhibits a high degree of co-occurrence, forming interacting pairs. This result is consistent with our previous findings. Finally, the polar-polar pair exhibits a PPMI equal to zero in all the regions, indicating that the contacts of these abundant residue types tend to be random and less specific.

## 3 Conclusion

In this work, we presented a detailed analysis of the interactions sustaining protein-protein interfaces, focusing on the distinct regions that form an interface. Our results confirm that the stability of an interface is not primarily determined by its overall size or specific residue composition, nor do stable interfaces exhibit a dramatically different residue composition compared to transient ones. Instead, stability is influenced by the partitioning of residues across the main regions of an interface—the core, support, and rim—-which differ in their hydration levels. Specifically, we found that the support region of stable interfaces, enriched in aromatic, polar, and charged residues, hosts a significant number of interactions, making it a critical component of stable interfaces. Overall, while the core and support regions of stable interfaces exhibit a higher heterogeneity in interacting pairs, they display a degree of local organization similar to that of the protein interior, as demonstrated by our residue co-occurrence analysis.

This analysis was enabled by our newly developed MICLOT software, which has proven to be a powerful tool for identifying diverse interaction types, classifying interacting residue pairs, and characterizing their local interaction contexts in stable and transient interfaces.

We anticipate that MICLOT will be valuable for analyzing the time evolution of protein-protein interfaces during molecular dynamics simulations of macro-molecular events in the cytoplasm, such as the formation of dynamic enzyme assemblies. Ultimately, the detailed insights provided by this software could aid in predicting stable interfaces and distinguishing them from transient ones [38, 39]. In the future, MICLOT could be extended to analyze indirect non-bonded interactions, such as water-mediated hydrogen bonds, which also contribute to interface stabilization [40]. Furthermore, incorporating additional physico-chemical properties of amino acids [41] or molecular descriptors [42] could enhance the depth of the analysis.

Our analysis software used in this study is freely available as open source, with a complete documentation, on GitHub: https://github.com/TMiclot/MICLOT.

## 4 Methods

### 4.1 Interaction types identified by the software

This part gives a general overview, but complete documentation of the recognition parameters is provided in the GitHub repository https://github.com/TMiclot/MICLOT.1) Identified non-bonded interactions that promote PPI are: *n* → *π*^*∗*^: Interaction between the carbon (C) and oxygen (O) atoms of two carbonyl groups (C=O) of the backbone. *C-bond*: *n* → *σ*^*∗*^ electron delocalization between a donor *C* ^*sp*3^ and an acceptor carbonyl O. *Chalcogen bond*: Attractive interaction between sulfur (S) or selenium (Se) and an electrophilic region [43]. *S/Se mediated H-bond*: Hydrogen bond involving a S or Se as H-donor or H-acceptor. *Hydrogen bond*: Attractive interaction between a hydrogen (H) with a more electronegative atom [44]. *π* − *H bond*: Attractive interaction between a hydrogen (H) with the *π* part of an aromatic cycle. *π* − *π*: Interaction between the *π* parts of two aromatic residues. *π* − Quadrupole: Interaction between the *π* part of one aromatic residue and the quadrupole part of another. *Quadrupole - Quadrupole*: Interaction between the quadrupole parts of two aromatic residues. Amino − *π*: Interaction between the *π* part of an aromatic cycle and the amino group of a glutamine or asparagine. *Aromatic - Charge*: Positively and negatively charged amino acids can interact with the *π* area or the quadrupole area of an aromatic cycle. *Salt bridge*: Electrostatic interaction involving both H-bond and ionic bond between amino acids with opposite charges. *van der Waals*: Interaction between atoms due to dipole-induced dipole and dispersion forces [45]. *Hydrophobic*: Interaction between two close hydrophobic residues. 2) Repulsion and clash involve a pair of close residues with the same charge or different hydropathy. As a consequence, all repulsion/clash types are: Anion–Anion, Cation–Cation, Hydrophobe–Hydrophophile. Two residues are in *repulsion* if the distance between the backbones is close but the side chains are far apart, then repulsion occurs. In contrast, two residues are in *clash* if the distance between the backbones is close and the side chains are close. The presence of a repulsion or a clash is not optimal; however, their overall effect depends on the presence of other surrounding residues interacting with the two side chains.

From all possible pairs in a system, only residues for which the distance between teir *C*_*α*_ atoms is not greater than 14 Å are kept in the analysis process. These values allow identifying short-range and long-range interactions. Because our software is able to identify multiple interaction types and their subtypes, some interaction (sub-)types have been concatenated for ease of analysis. In addition, alternative residues positions, if any, recorded in the structure file are not taken into account.

### 4.2 Dataset preparation

All complex structures come from five validated databases: Protein-Protein Docking Benchmark 5.5 [46, 47] (BM5.5), SabDab [48], PPI4DOCK [49], 3DComplex [50] v6.0 and PDBind [51, 52, 53] v2020. For this last one, only structures with two chains (dimers) were retained. In addition, all information concerning protein family, reference article (PubMed format), classification, organism, presence of mutation, method used to solve the structure, resolution (if available), and the UniProtKB [54] number was taken from RCSB PDB [55].

The first step of the selection process consisted in conserving only structures with known binding affinities. These values came from the database from which the structure originated, if not, from the PDBind database. Next, the selection process only retained structures with a resolution below 2.5 Å or NMR structures. Subsequently, the UniprotKB entry was used to remove redundancy: all duplicates were removed to ensure that each entry was completely unique in the dataset. Structures with many residues, where too many atoms were missing to be rebuilt, were not taken into account. The final dataset contained 413 protein hetero-complexes. Please note that preselection was applied to structures from PDBind and 3DComplex. For PDBind, only dimeric complexes were kept. For 3DComplex, the selected structures were not identical, not homologous, and domain architectures of chains 1 and 2 were unique (for SCOPE and PFAM definition). For more information see SI section A and SI section B.

### 4.3 Separation of complexes into stable and transient interfaces

Adopting a similar approach to that employed by Grassmann et al. [20], we segregated our dataset into distinct categories. Complexes with log_10_(Affinity), where “Affinity” stands for *K*_*D*_, *IC*50, or *K*_*i*_, greater than -6 were classified as transient, whereas those with log_10_(Affinity) lower than -9 were designated as stable. Complexes falling between these two values were identified as intermediate. Figure 5 shows that the majority of structures in the dataset were classified as intermediate.

**Figure 5:**
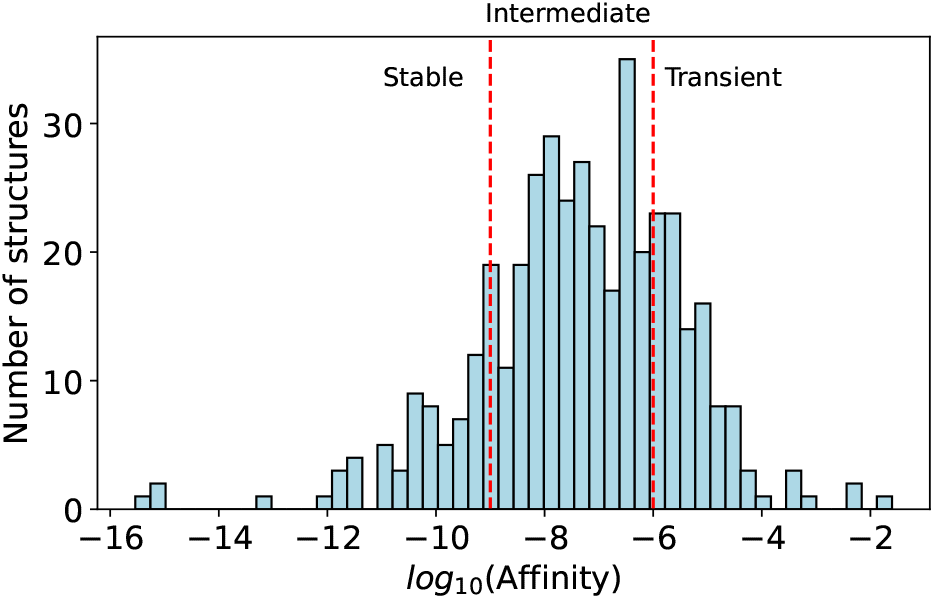
Distribution of complex affinities in the dataset. 94 complexes of the dataset are identified as transient interface and 66 are classified as stable, while 253 complexes are intermediate.

### 4.4 Identification of protein regions

We use the same relative ASA (rASA) based method as described by Levy [24] to distinguish five regions. A residue can be located at the surface, in the interior, or at the interface, which is divided into Core, Support and Rim regions. We go deeper in the nomenclature and consider a residue on the surface as hydrated if its rASA in complex is *> 40 %* [56]. Moreover, we distinguish residues on the rim as part of the interacting surface if their ΔrASA *>* 5%, or as part of the non-interacting surface (NIS) if their ΔrASA ≤ 5% [25]. For each residue, a set of structural regions is identified using theoretical and empirical maximum ASA values (*MaxASA*) from Samanta, Bahadur, and Chakrabarti [57], Lins, Thomas, and Brasseur [58], Tien et al. [59], Miller et al. [60], Rose et al. [61], and Hubbard [62]. Then the more frequent is selected as the region where the residue is located.

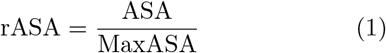

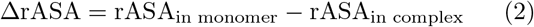

## Acknowledgements

We acknowledge funding support by the Czech Academy of Sciences (Lumina Quaeruntur Fellowship LQ200402301). All calculations were carried out on the local HPC of the J. Heyrovsky Institute of Physical Chemistry, and we thank Michal Tarana for the IT technical support.

## Declaration of use of generative artificial intelligence technologies

The manuscript was reviewed for grammatical accuracy using the HuggingChat tool with the Meta-Llama-3.1-70B-Instruct model, followed by a thorough manual review. The authors assume full responsibility for the content of the publication.

## Contributions

T.M. and S.T. conceptualized the work. T.M. wrote the analysis script and performed data analysis. T.M. and S.T prepared and edited the manuscript. S.T. supervised the work.

