## Supplementary Information for "Beyond contacts: the important role of the support region to distinguish stable and transient protein interfaces"

January 29, 2025

**E Non-bonded interaction trends in protein regions** **9**

**F Description and selection of residue pairs** **10**

**G Composition of interface region by residue classes** **11**

**H Interaction type properties** **11**

**J Composition of residue pairs and interactions in PPI interfaces** **17**

**K Relation between interface size and interaction and pair counts** **18**

**L Evaluating the diversity of pairs** **19**
L.1 Method . . . . . . . . . . 19
L.2 Results . . . . . . . . . . 20

**M Identification of key amino acid pairs** **21**

**N List of used structures** **23**

**References** **48**

### A Dataset structure quality

**Note** Information concerning each protein complex comes from keys in the website of the RCSB PDB [1].

More than half of the structures selected to make the dataset are dimeric complexes from the PDBind database and more than a quarter come from PPI4DOC (see Figure S1). Moreover, we used BBM5.5, while other used databases play a much smaller role in the creation of the dataset. The most important thing is not so much the database from which the structure comes as the species to which it belongs. *Homo sapiens* is the leading species, species, accounting for 57.3 % of the dataset, followed by model organisms: *Mus musculus*, *Escherichia coli*, *Saccharomyces cerevisiae* and multiple *Rattus* species. 9 other organisms including a virus (HIV) compose more than 1 % of the dataset and 27.8 % of other species. It is interesting to note that 2.4 % of the dataset correspond to synthetic constructs. However, considering these properties, the dataset essentially represents human protein complexes.

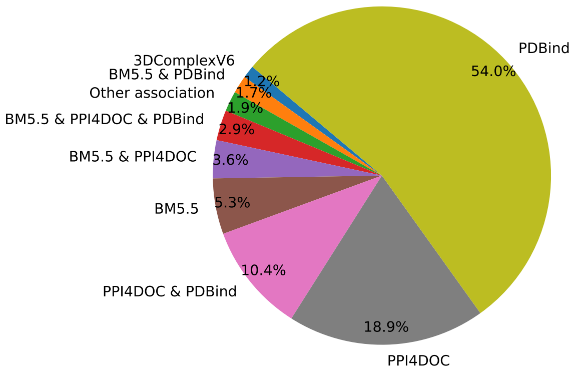

(a)

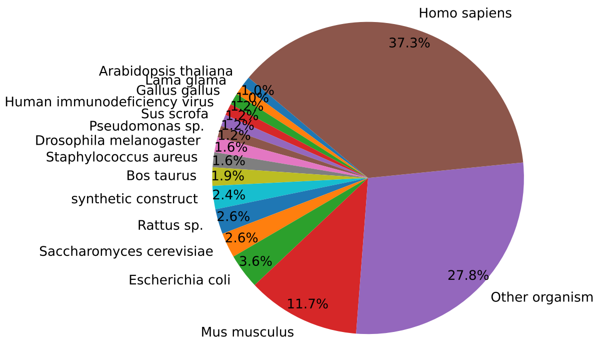

(b)

Figure S1: (a) Partitioning of selected structures among databases. Note that we only retain dimeric structures from PDBind, so the quality analysis returns a result for this selection and not the whole database. (b) Biological origin of the selected structures. Organisms classified as 'other' are those representing less than 1% of the dataset's diversity.

In the RCSB PDB database, the mutation information key indicates sequence mutations, but without giving more details. In general, mutations come from a mutagenesis strategy used by authors to ensure the crystallization of their studied protein [2, 3]. Alternatively, it can be due to the need of enzyme inhibition; as shown in Figure S2, a significant part of the dataset corresponds to enzymes: hydrolase in particular. In this regard, protein classification shows a large range of categories in the dataset. The biggest is *immune system*, then it is hydrolase-involving complexes: *hydrolase* and *hydrolase/jydrolase inhibitor* and the third is *protein binding*. 11 additional categories compose more than 1% of the dataset, while 45.5 % correspond to other categories. As a consequence, the dataset is not representative of a particular type of protein complex, but has enough diversity to be a general representation.

Figure S3 shows that protein complex structures are mainly solved by X-ray diffraction with resolution between 1.80 to 2.50 Å. Structures with lower resolution are less represented. This means that the structures have a good degree of quality, but that the position of the hydrogen has to be remodeled.

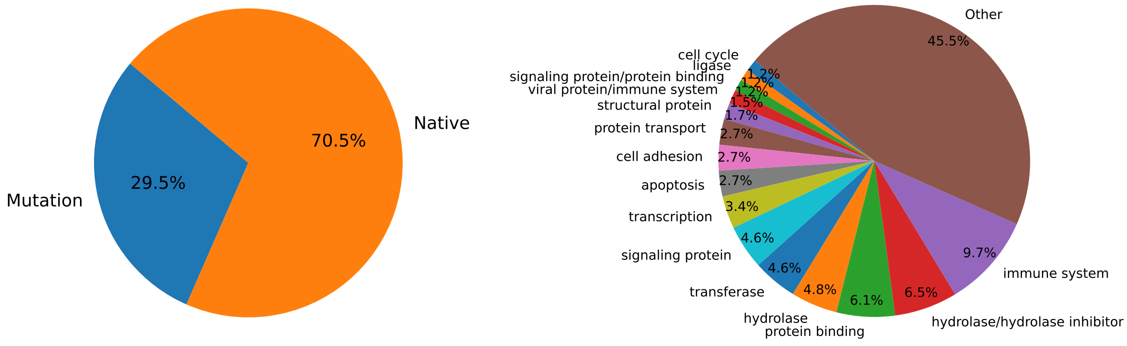

(a) (b)

Figure S2: (a) Proportion of mutated structures with (b) the classification of proteins constituting the dataset. Classification categories identified as 'other' are those representing less than 1% of the dataset's diversity.

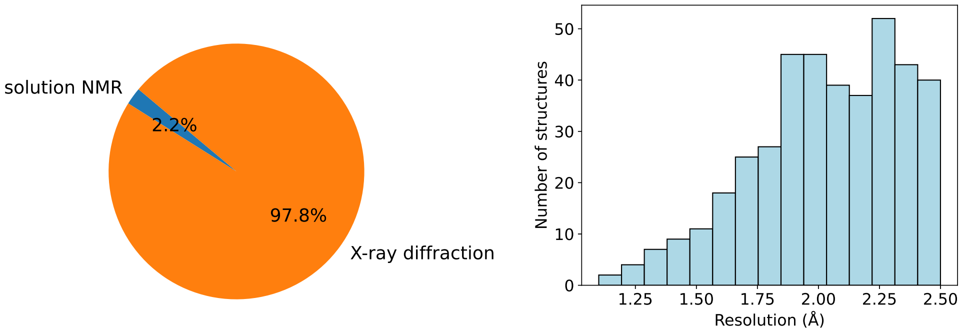

(a) (b)

Figure S3: (a) Proportion of methods used to solve structures with (b) the resolution distribution of X-ray solved structures.

### B Dataset preparation

#### B.1 Add missing atoms and optimize hydrogen bonds

All heteroatoms are removed from the original file, as well as unknown residues (UNK). Then PDB2PQR [4] is used with its PROPKA [5, 6] parsing option to assign the protonation states of residues in the structure and add missing hydrogen atoms. pH is set at 7.2 when unknown. Another option is used to ensure that rebuilt atoms do not overlap or are not too close from each other. In addition, the procedure is set to ensure optimisation of hydrogen bonds by the software. The output of PDB2PQR is exported in the pdb file format and submitted to PDBFixer [7] to rebuild missing heavy atoms and ensure that atom names fit those of AMBER ff14SB force field [8]. The same pH value is used as the one provided to PDB2PQR.

#### B.2 Optimising hydrogen atom positions

The previous steps rebuild hydrogen atoms in non-optimised positions, except for those making hydrogen bonds. To ensure that there are no clashes or bad positions of hydrogen atoms, a vacuum minimization is performed using openMM [7]. A constraint is imposed on hydrogen bonds to avoid loss of previously optimized hydrogen bonds, except those found in NMR structures. Another restraint is set at 1.0e5 kJ/nm to ensure heavy atoms do not move from their initial positions. This ensures that the structure of the complex does not change compared to the experimental structure and avoids force field-dependent modifications. No maximum number of minimisation steps is used to ensure that the minimisation continues until convergence.

#### B.3 Fix missing bonds

In some cases, connected atoms may not be properly identified by software such as openMM or MDTraj [9]. To avoid this problem, bonded atoms are recognized using same method as VMD [10]: two atoms are bonded if their distance (d) is $d_{min} \leq d \leq d_{max}$, where:

$$d_{max} = 0.6 \times (d_{vdw\ atom\ 1} + d_{vdw\ atom\ 2}) \tag{S1}$$

$$d_{min} = d_{covalent\ atom\ 1} + d_{covalent\ atom\ 2} \tag{S2}$$

Then the CONNECT part of the pdb file is written for all bonded atoms identified.

Cysteine bridges are identified based on their chalcogen atoms (sulfur or selenium) distance. Cysteines involved in a bridge are renamed as CYX, while selenocysteines are renamed as XSE. Finally, the CONNECT line is written for the bonded S/Se atoms.

### C Dataset: Interface areas of protein assembly

**Method** The accessible surface area (ASA) of associated and free monomers forming interface 1 was calculated with the Shrake and Rupley algorithm [11] with the Golden Section Spiral algorithm as implemented in MDTraj [9]. All atoms, including hydrogen, of residues are taken into account. Then the interface is calculated as the buried surface area (BSA) divided by 2, and the relative interface is the interface area divided by the surface of each complex partner. Equations are those below:

$$\text{BSA} = (\text{ASA}_{\text{partner i}} + \text{ASA}_{\text{partner j}}) - \text{ASA}_{\text{complex}} \tag{S3}$$

$$\text{interface} = \frac{\text{BSA}}{2} \tag{S4}$$

$$\text{relative interface} = \frac{\text{interface}}{\text{ASA}_{\text{partner i or j}}} \times 100 \tag{S5}$$

**Results** Interface distribution of the dataset peaks at around 1000 Å$^2$, which is in line with the average of 1227 Å$^2$ reviewed [12]. However, the interface profile below 1000 Å$^2$ is different and shows relatively reduced information in this region. It is interesting to note that, despite the wide range of interface area values, the interface area represents only around 5-15% of the total surface area of each protein partner forming a complex.

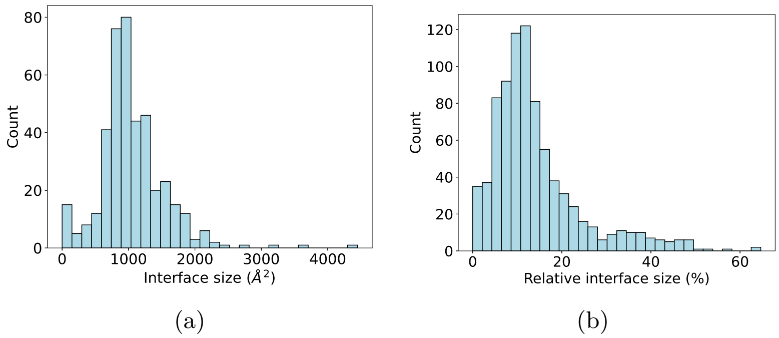

Figure S4: (a) Distributions of interface size and (b) of relative interface areas in the dataset.

### D Comparison between stable and transient interfaces

#### D.1 Overview of interfaces areas

Distribution of interface sizes between stable and transient complexes exhibits a high degree of similarity. Similarly, the distribution of relative interface areas also shows a marked resemblance. The difference lies in the count of structures, which is directly correlated with the size difference between the two data subsets. This suggests that considering the whole interface may not be a highly discriminative approach, as the overall characteristics of stable and transient complexes appear to be comparable. To gain a deeper understanding of the underlying differences, we adopted a more nuanced approach by dividing the interface into distinct regions: Core, Support, and Rim, as described by [13]. By examining these regions separately, we aimed to uncover more subtle distinctions that may not be apparent when considering the interface as a whole.

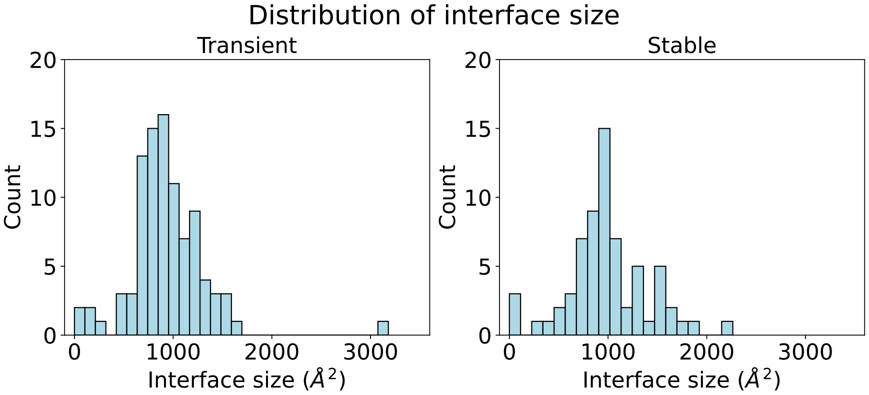

(a)

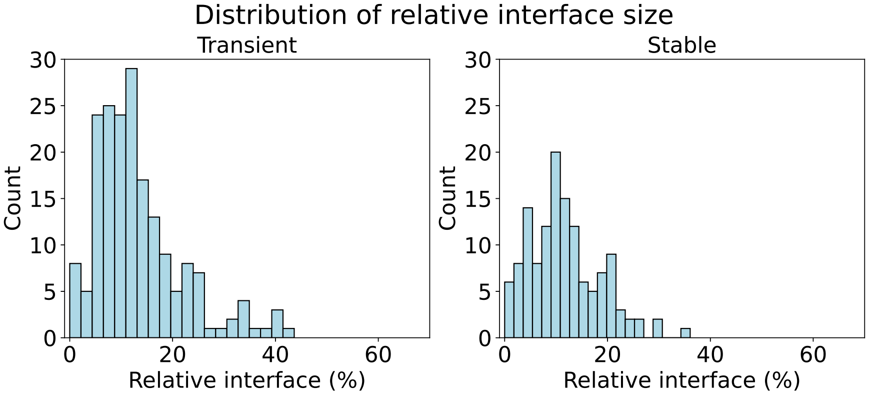

(b)

Figure S5: (a) Distributions of interface size and (b) of relative interface areas show few differences for stable and intermediate complexes in the dataset.

#### D.2 Core, Support and Rim ratio

| | Total interface ($Å^2$) | Interface Core ($Å^2$) | Interface Support ($Å^2$) | Interface Rim ($Å^2$) | Ratio Core (%) | Ratio Support (%) | Ratio Rim (%) |
|---|---|---|---|---|---|---|---|
| **Stable** | | | | | | | |
| mean | 989.874677 | 607.208394 | 123.476492 | 259.189791 | 58.580564 | 11.501436 | 29.918000 |
| std | 425.296214 | 275.951603 | 72.097827 | 125.723235 | 13.236188 | 4.769504 | 15.900780 |
| 25% | 796.521692 | 461.666735 | 81.986300 | 178.704510 | 53.615981 | 9.168470 | 19.743660 |
| 50% | 934.196376 | 579.348745 | 102.934854 | 241.232073 | 60.788616 | 11.522166 | 27.612149 |
| 75% | 1217.829464 | 774.178134 | 170.973008 | 299.066617 | 67.002718 | 15.455441 | 33.495636 |
| **Intermediate** | | | | | | | |
| mean | 1092.089007 | 664.355783 | 106.509581 | 321.223643 | 58.438802 | 10.008344 | 31.552854 |
| std | 545.701572 | 354.982146 | 62.237815 | 178.345136 | 13.036700 | 6.265982 | 14.088080 |
| 25% | 784.718914 | 451.388460 | 64.511889 | 219.188447 | 54.443073 | 7.212961 | 24.083959 |
| 50% | 995.119005 | 638.520167 | 99.387733 | 283.506684 | 61.538548 | 9.349579 | 28.718762 |
| 75% | 1326.425056 | 838.759044 | 145.916346 | 395.864791 | 66.131390 | 12.158629 | 35.452407 |
| **Transient** | | | | | | | |
| mean | 933.722276 | 554.794663 | 90.295209 | 288.632405 | 56.450750 | 10.029828 | 33.519422 |
| std | 400.471166 | 293.255516 | 60.658658 | 125.472156 | 14.281923 | 7.388917 | 14.577434 |
| 25% | 740.403233 | 419.622314 | 56.147998 | 208.572851 | 53.272626 | 6.336329 | 26.268331 |
| 50% | 878.268117 | 522.612732 | 82.317239 | 270.496404 | 58.892119 | 8.388054 | 31.737366 |
| 75% | 1111.908862 | 651.509193 | 115.613281 | 356.622329 | 65.125049 | 12.776661 | 36.997802 |

Table S1: Descriptive statistics of interface region size and their relative size in an interface.

### E Non-bonded interaction trends in protein regions

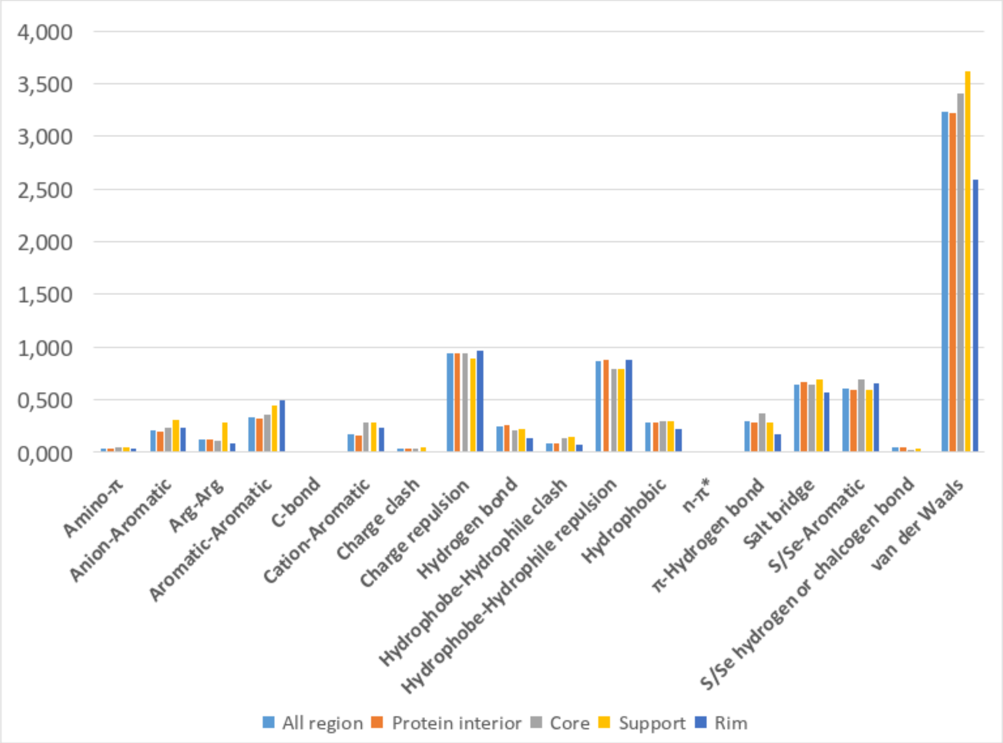

Figure S6: Non-bonded interaction trends in protein regions. The values represent the average count of the given interaction type for residue pairs that can engage in this type of interaction.

### F Description and selection of residue pairs

We only consider residues that form a pair if they have an identified interaction type Describing a pair using residues names and secondary structures leads to a complex description for the analysis of their interaction and is not very detailed concerning the interaction between the side chains and backbone. To ensure a simplified but more complete description of pair, residues are divided into two beads: one for the side chain and one for the backbone. The side chain bead has the same name as the residue; for example: the side chain of the arginine residue is named ARG. For the backbone bead, the name associated with it does not correspond to the name of the residue, but to the type of secondary structure. This choice was made because it is the backbone that indicates in which secondary structure a residue is located, and also to differentiate these backbone conformations. Thus, there are three different bead names: backbone in coil (BB_C), backbone in strand (BB_E), and backbone in helix (BB_H). As a consequence, there are 23 beads in this nomenclature: 20 for side chains and 3 for backbone types, and we can expect approximately $\frac{23(23+1)}{2} = 276$ possibilities of pairs, but 278 pairs have been identified as interacting.

All consecutive residues ($i$,$i$+1) are excluded. Finally, only pairs involving residues in the interior (I) are considered as part of protein interior. Analogously, we handle pairs in the core (C) and support (P), respectively, for PPI. Finally, residues in interacting rim (R) and NIS rim (N) are considered to study the extended area around the interface.

### G Composition of interface region by residue classes

We now examine the distribution of residues in the interface, using their physico-chemical classes as defined by Pommié et al. [14], rather than their individual names as previously considered.

Charge and polarity are key parameters used to understand protein complex formation, describe relevant interfaces, and predict binding affinity [15, 16, 17]. Our investigation analyses the distribution of residues classes in the regions forming the interface, as shown in Figure S7. It reveals that there is no significant difference in charge distribution between stable and transient interfaces in the core and support regions. However, a slight variation is observed in the rim, where transient complexes exhibit a rim (interaction) with a higher proportion of both positive and negative charges, whereas their rim (NIS) is more positively charged. This finding is consistent with previous work by Grassmann et al. [18]. In contrast, polarity remains unchanged in the rim but exhibits high variation in the core and support regions, with stable interfaces displaying higher polarity in these regions. This can be partially explained since Pommié et al. [14] defined Tyr as polar due to its hydroxyl group (-OH).

Furthermore, examination of other amino acid properties, such as hydropathy, reveals no significant differences in the rim (NIS), but notable variations in other regions. Analysis of potential hydrogen bonding partners reveals no significant variation in the rim but a higher proportion of none H-acceptor/donor residues in the core and support regions of transient interfaces. This observation imply that transient interfaces are less able of forming hydrogen bonds than stable ones. Chemical classes exhibit more variation in their distribution across interface regions. Notably, aromatic residues are less represented in transient interfaces. In contrast, the rim (interaction) of transient complexes contains fewer hydroxyl residues but more acidic and basic residues compared to stable interfaces. In the support and core regions, the primary variations are observed for aliphatic and aromatic residues, with stable interfaces showing higher aromaticity and lower aliphatic content compared to transient interfaces. Additionally, amide residues are slightly underrepresented in transient interfaces in the core region.

### H Interaction type properties

#### H.1 Distribution of *van der Waals*

Number of van der Waals contacts was counted between residues in all non-consecutive pairs, without any other selection. The figure below shows that a typical vdw interaction between residues forming a pair involves less than 5 contacts.

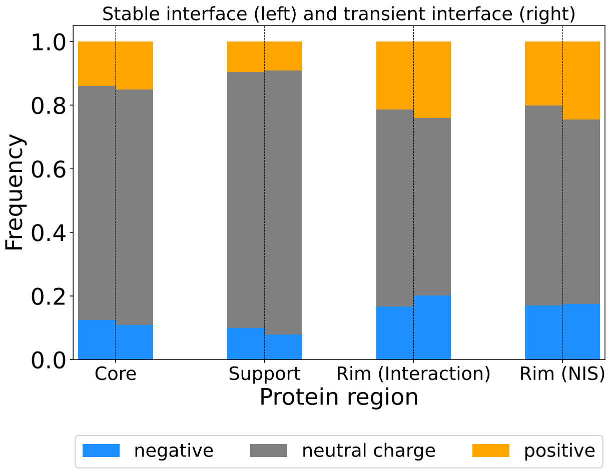

(a) Charge

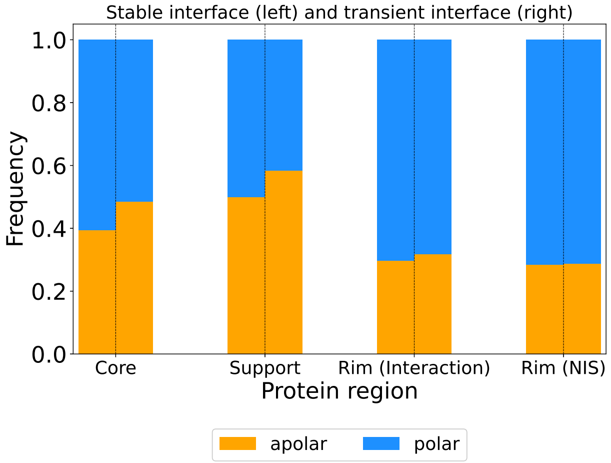

(b) Polarity

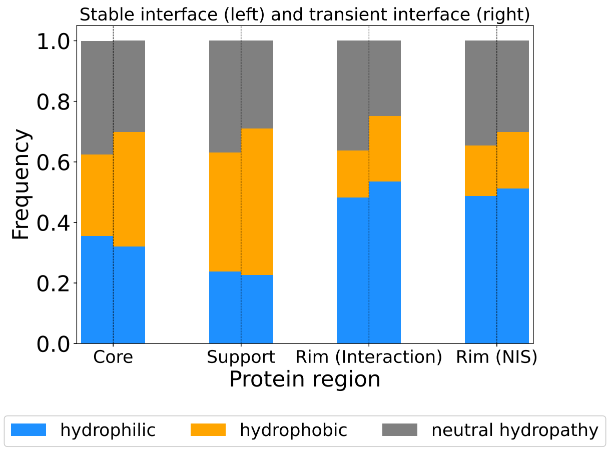

(c) Hydropathy

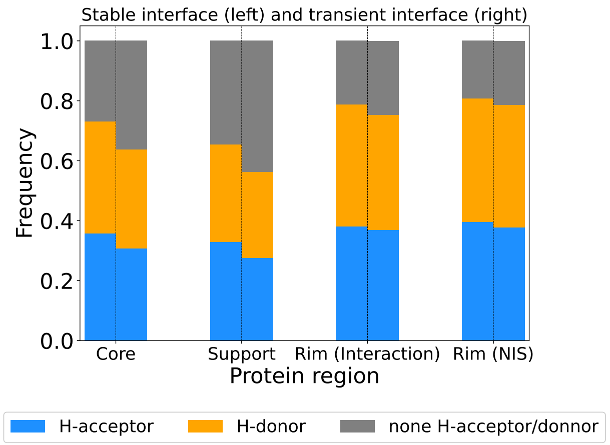

(d) Hydrogen bond

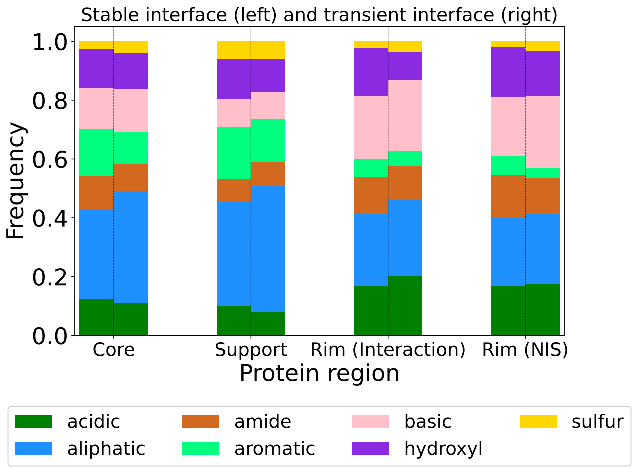

(e) Chemical class

Figure S7: Distribution of classes in stable and transient interface regions.

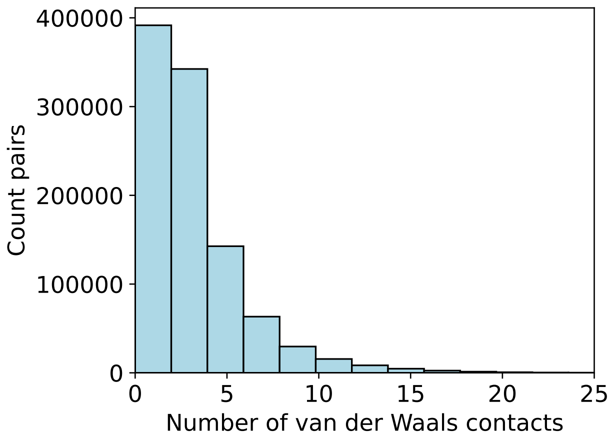

Figure S8: Distribution of number of van der Waals contact between non-consecutive residue pairs.

#### H.2 Comparing interaction type count in context of protein interior and PPI

##### Results

| Interaction type | All | Interior | Core | Support | Rim (interaction) | Rim (NIS) |
|---|---|---|---|---|---|---|
| Amino−Π | 241 | 197 | 24 | 16 | 3 | 1 |
| Aromatic−Anion | 1380 | 1119 | 127 | 102 | 29 | 3 |
| Aromatic−Aromatic | 1990 | 1734 | 106 | 141 | 9 | 0 |
| Aromatic−Cation | 943 | 690 | 156 | 66 | 30 | 1 |
| Aromatic−S/Se | 2270 | 2074 | 104 | 86 | 6 | 0 |
| C-bond | 105 | 96 | 3 | 3 | 3 | 0 |
| Charge clash | 217 | 160 | 29 | 11 | 16 | 1 |
| Charge clash: ARG−ARG | 112 | 70 | 19 | 15 | 7 | 1 |
| Charge repulsion | 5725 | 3872 | 774 | 227 | 768 | 84 |
| Hydrogen bond | 43435 | 39235 | 2379 | 1309 | 469 | 43 |
| Hydrogen bond: H−Π | 22565 | 19988 | 1644 | 832 | 101 | 0 |
| Hydrophobic | 21940 | 20684 | 688 | 544 | 24 | 0 |
| Hydrophobic−Hydrophilic clash | 6512 | 5466 | 644 | 304 | 90 | 8 |
| Hydrophobic−Hydrophilic repulsion | 59262 | 52821 | 3690 | 1618 | 1043 | 90 |
| S/Se Chalcogen or Hydrogen bond | 831 | 774 | 28 | 25 | 4 | 0 |
| Salt bridge | 1513 | 1015 | 244 | 68 | 182 | 4 |
| $n \rightarrow \Pi^*$ | 38 | 34 | 4 | 0 | 0 | 0 |
| van der Waals | 1233848 | 1106633 | 71726 | 40144 | 14206 | 1139 |

Table S2: Total interaction type counts in the various interfaces regions for the entire dataset.

### I Pair class partitioning in protein regions

In this section, we analyze the protein region location of residue pairs based on their classes, as described in the work by Pommié et al. [14]. (2004). Glycine has a unique hydrogen atom in its side chain, rendering its properties similar to those of the backbone. Consequently, we assigned Gly properties to all backbone beads (BB_C, BB_H, BB_E). Finally, only pairs in which each partner is located in the same protein region are considered. This means that interfacial pairs: core–support, support–rim (interaction), and rim (interaction)–rim (NIS) are not taken into account.

#### I.1 Partitioning in the different protein regions

**Results**

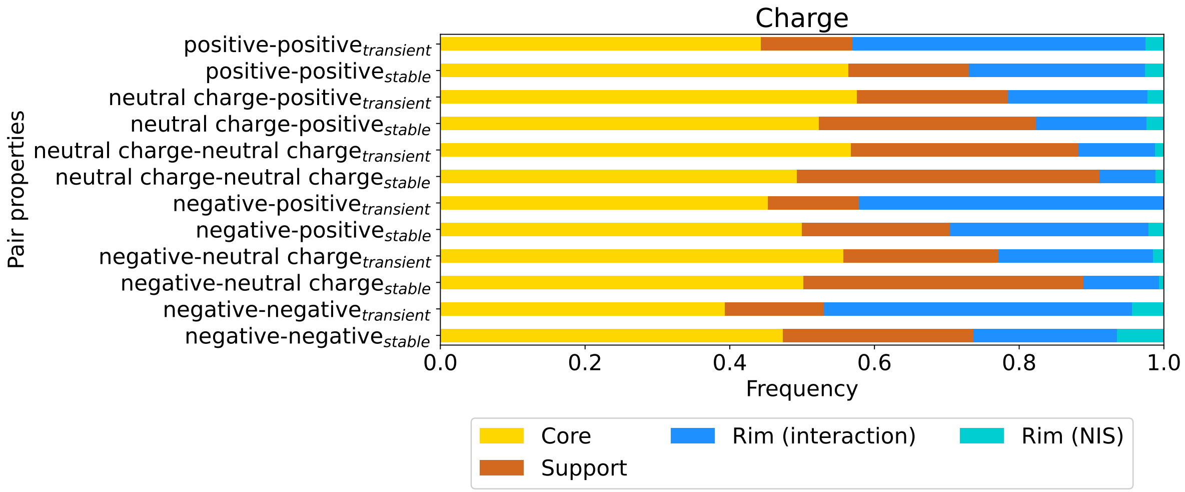

Figure S9: Distribution of pairs in the different protein regions, based on their charge property.

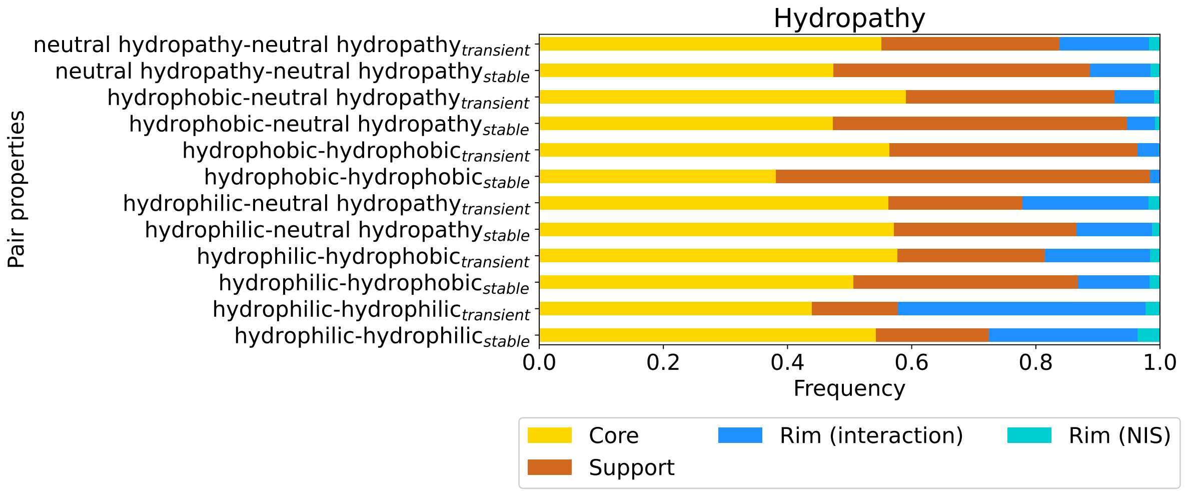

Figure S10: Distribution of pairs in the different protein regions, based on their hydropathy property.

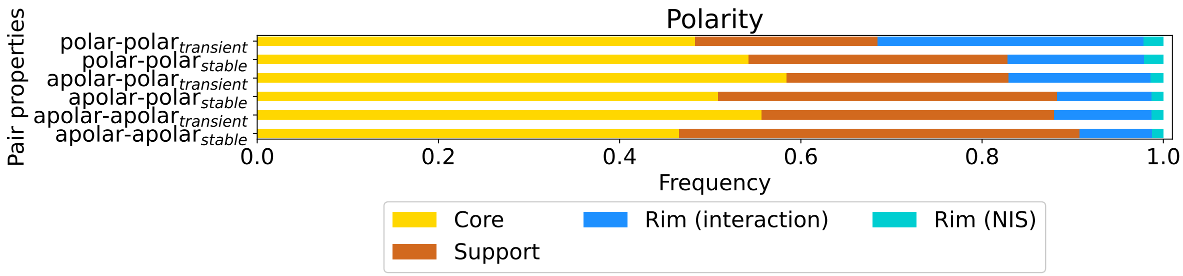

Figure S11: Distribution of pairs in the different protein regions, based on their polarity property.

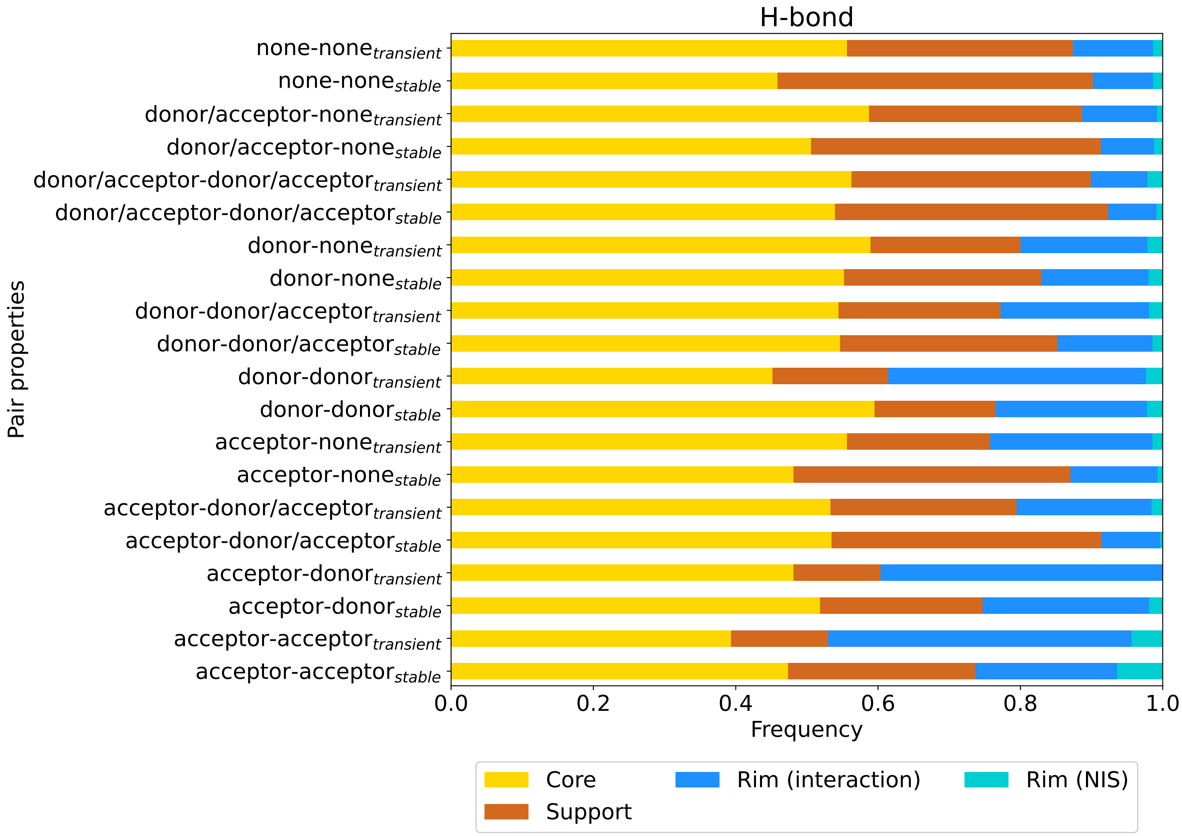

Figure S12: Distribution of pairs in the different protein regions, based on their hydrogen donnor/acceptor property.

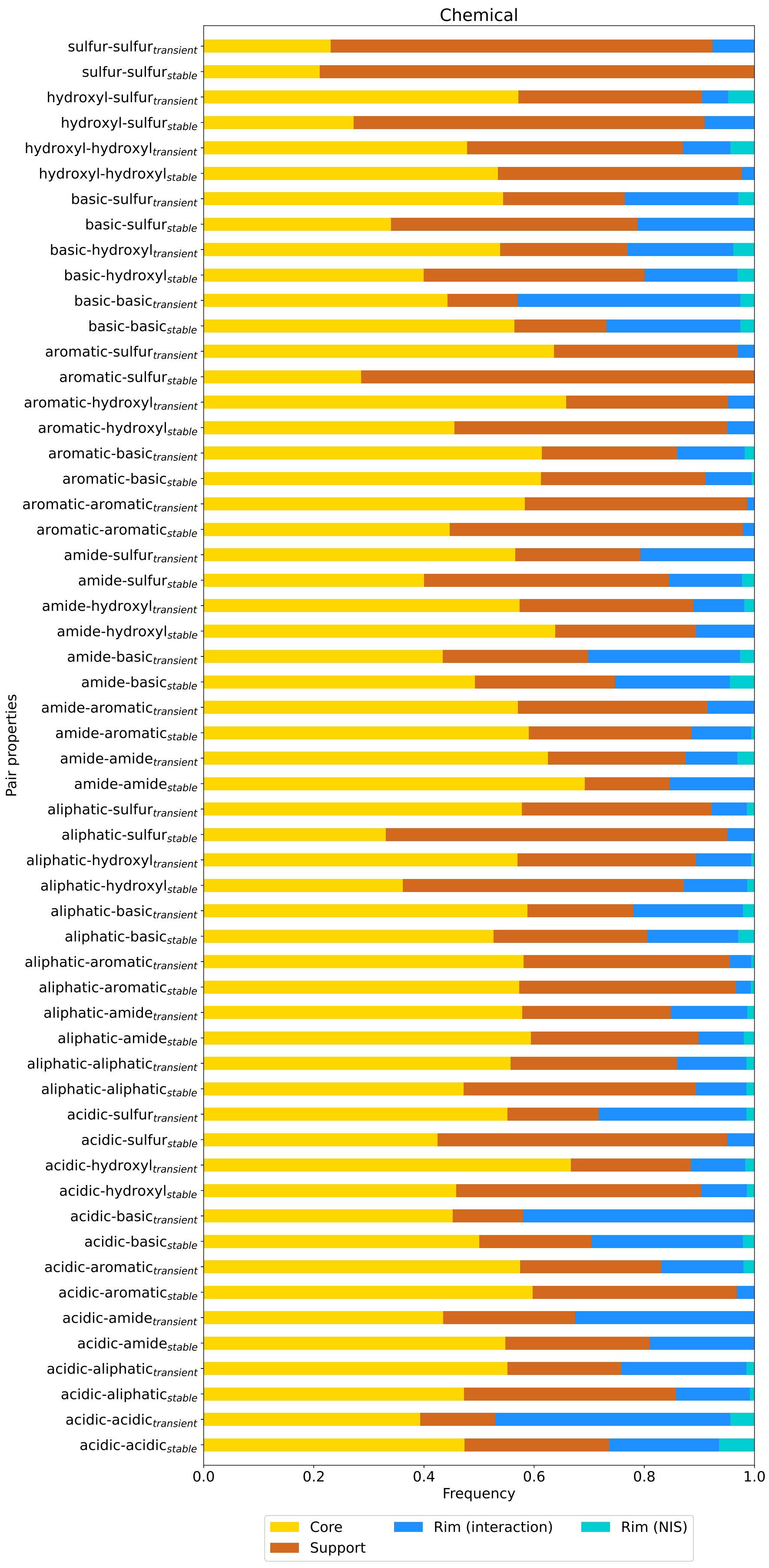

Figure S13: Distribution of pairs in the different protein regions, based on their chemical class property.

### J Composition of residue pairs and interactions in PPI interfaces

**Method** The primary objective of this analysis is to provide a comprehensive overview of the composition of protein–protein interfaces in terms of their residue pair and interaction content. To achieve this, we choose not to differentiate between stable, transient, and intermediate complexes, nor to distinguish between different protein regions that contribute to the interface. Instead, we focus on analyzing the distributions of interaction types (e.g., van der Waals, hydrogen bonds, etc.) and the count of interactions. To illustrate this approach, we can use van der Waals (vdW) interactions as an example: For each pair of residues, we assign a binary value (1 or 0) indicating whether a vdW interaction is present or absent, regardless of the number of vdW interactions involved. In contrast, the interaction count focuses on the actual number of vdW interactions between the pair, such that a value of 5 represents five vdW interactions. These approaches are applied to all interaction types and each PDB structure. The same protocol is employed for residue pairs, utilizing a simplified two-bead representation (i.e., backbone and side chain). Finally, we plot the distribution of these values against the count of PDB structures, providing insight into the major characteristics of the dataset.

**Results**

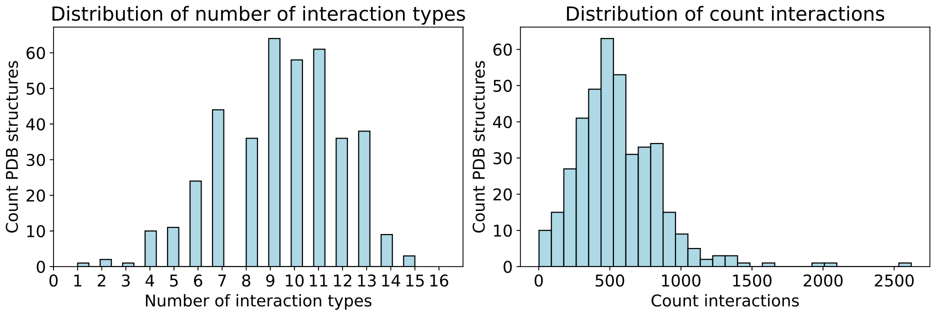

(a)

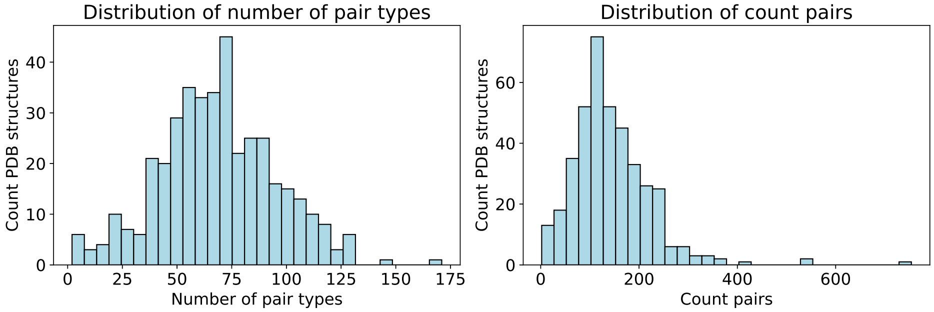

(b)

Figure S14: (a) Distribution of the number of interaction type and the count of interaction over the structure dataset. (b) Distribution of the number of pair type and the count of pair over the structure dataset.

### K Relation between interface size and interaction and pair counts

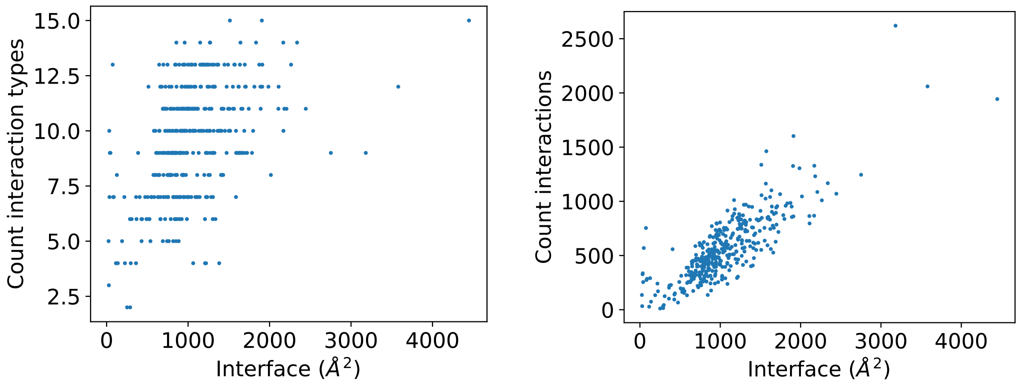

(a) (b)

Figure S15: Correlation between the interface size and (a) the count of interaction types or (b) the count of interactions forming the interface.

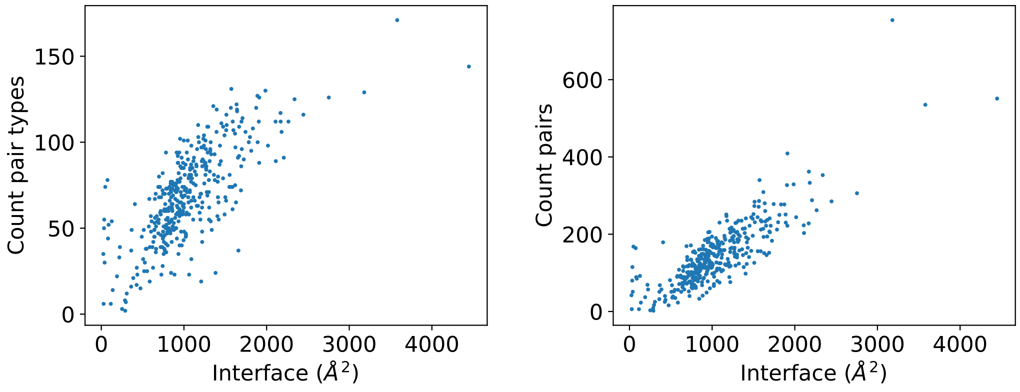

(a) (b)

Figure S16: Correlation between the interface size and (a) the count of pair types or (b) the count of interacting pairs.

### L Evaluating the diversity of pairs

#### L.1 Method

This analysis aims to provide a comprehensive overview of the diversity of residue pairs in protein interfaces. However, a general analysis using a simplified two-bead description of pairs (see SI section J) may not be sufficient, because different side chains can exhibit similar properties [14]. To address this limitation, we characterize residue pairs based on their mixed polarity and charge properties, which are commonly used in binding energy prediction methods [17], and specify the charge type as either positive or negative. The table below summarizes the residue properties used. Furthermore, we adopt a more detailed approach by considering the protein regions, including the interior, core & support, and rim, which is further subdivided into rim (interaction) and rim (NIS). Note that this case, and compared to previous analysis, interactions involving the backbone are excluded from this analysis. This is done to avoid artificial inflation of apolarity values, as the backbone properties are similar to those of glycine. The rationale for this decision is that the backbone is a common feature of all residues, and its inclusion would introduce a bias towards over-representing apolar interactions in the statistics.

| Side chain beads | Property | Side chain beads | Property | Removed beads | Property |
|---|---|---|---|---|---|
| ALA | apolar | LEU | apolar | BB_C | apolar |
| ARG | positive | LYS | positive | BB_H | apolar |
| ASN | polar | MET | apolar | BB_E | apolar |
| ASP | polar | PHE | apolar | | |
| CYS | apolar | PRO | apolar | | |
| GLN | polar | SER | polar | | |
| GLU | negative | THR | polar | | |
| GLY | apolar | TRP | apolar | | |
| HIS | positive | TYR | polar | | |
| ILE | apolar | VAL | apolar | | |

Table S3: The properties of individual residues are utilized to derive the properties of residue pairs. Notably, the backbone (BB) is similar to Gly in terms of its properties, and as such, the properties of BB are assigned based on those of Gly.

To assess the diversity of residue pairs, we calculate the Shannon entropy (H) [19] for each protein region. This metric provides a quantitative measure of the information content, which reflects the degree of uncertainty or randomness in the data. This approach compares the frequency of each pair within a protein region, allowing us to evaluate the complexity of the region's composition. A low entropy value indicates that the region is comprised of a limited diversity of pairs, making it more predictable. Conversely, a high entropy value suggests that the region exhibits a high diversity of pairs, rendering it more challenging to predict due to its randomness.

$$\mathrm{H}_{\text{pair in region}} = -\mathrm{Frequency}_{\text{pair in region}} \times \log_2(\mathrm{Frequency}_{\text{pair in region}}) \tag{S6}$$

$$H_{region} = \sum H_{pair\ in\ region} \tag{S7}$$

The entropy value provides insight into the characteristics of the studied system. However, to determine if the system exhibits a random distribution, it is essential to compare the entropy value with the maximum entropy ($H_{max}$). This comparison enables us to assess whether the frequency of each pair is uniform, indicating a completely random distribution. A simple approach to quantify this comparison is to calculate the difference ($\Delta H$) between the system's entropy (H) and $H_{max}$. Although this value provides useful information, it only indicates the extent to which the pair population deviates from a completely random distribution. To gain a more nuanced understanding, we employ the evenness index (J) [20], which ranges from 0 to 1 and reflects the similarity in the frequency of each pair within a protein region. A low value of J indicates that a few pairs dominate the population, whereas a high value suggests that the pair population is more homogeneous, exhibiting a high degree of evenness.

$$H_{max} = -\frac{1}{N_{pair\ types}} \times \sum_{n=1}^{N_{pair\ types}} \log_2 \left( \frac{1}{N_{pair\ types}} \right) \tag{S8}$$

$$= -\log_2 \left( \frac{1}{N_{pair\ types}} \right) \tag{S9}$$

*With the properties used to descripe pair, as show in Table S3, value of $N_{pair\ types}$ is equal to 10.*

$$\Delta H = H_{max} - H_{region} \tag{S10}$$

$$J = \frac{H_{region}}{H_{max}} \tag{S11}$$

#### L.2 Results

| **Pair** | **Interior** | **Core & Support** (Stable) | **Core & Support** (Transient) | **Rim** (Stable) | **Rim** (Transient) |
|---|---|---|---|---|---|
| apolar-apolar | 0.401 | 0.141 | 0.225 | 0.023 | 0.043 |
| apolar-negative | 0.053 | 0.062 | 0.071 | 0.068 | 0.092 |
| apolar-polar | 0.311 | 0.320 | 0.305 | 0.216 | 0.189 |
| apolar-positive | 0.111 | 0.123 | 0.154 | 0.141 | 0.140 |
| negative-negative | 0.003 | 0.013 | 0.010 | 0.054 | 0.056 |
| negative-polar | 0.018 | 0.060 | 0.037 | 0.089 | 0.082 |
| negative-positive | 0.005 | 0.021 | 0.017 | 0.050 | 0.054 |
| polar-polar | 0.057 | 0.134 | 0.079 | 0.125 | 0.072 |
| polar-positive | 0.032 | 0.093 | 0.063 | 0.118 | 0.107 |
| positive-positive | 0.010 | 0.034 | 0.038 | 0.116 | 0.163 |

Table S4: Pair property frequencies in interior, core & support and rim of stable and transient complexes.

### M Identification of key amino acid pairs

**Method** In this section, we use the same pair property descriptors as in SI section L. To identify key pairs, we draw inspiration from Natural Language Processing (NLP) methods, where residue pairs are treated as *words* or *word combinations*. Specifically, we derive two weighting functions: a modified term frequency-inverse document frequency (tf-idf) [21, 22] and positive PMI (PPMI) [23, 24, 25], derived from the pointwise mutual information (PMI) [26, 23, 27]. The tf-idf metric characterizes the structural frequency of a pair, reflecting the number of times an interacting residue pair appears in the sub-dataset: interior, transient interface, stable interface. Values below 1 indicate pairs widely represented across our sub-datasets.

$$\mathrm{tf}_{\mathrm{pair}} = \begin{cases} 1 + \log_{10} count(pair) & \text{if } count(pair) > 0 \\ 0 & \text{oherwise} \end{cases} \tag{S12}$$

$$\mathrm{df}_{\mathrm{pair}} = \text{Count of PBD where a given pair is found} \tag{S13}$$

$$\mathrm{idf}_{\mathrm{pair}} = \log_{10}\left(\frac{\text{Number of PDB in the sub} - \text{dataset}}{\mathrm{df}_{\mathrm{pair}}}\right) \tag{S14}$$

$$Where:\ N_{PDB\ interior} = 413,\ N_{PDB\ stable\ interface} = 66,\ N_{PDB\ transient\ interface} = 94$$

$$\mathrm{tf} - \mathrm{idf}_{\mathrm{pair}} = \mathrm{tf}_{\mathrm{pair}} \times \mathrm{idf}_{\mathrm{pair}} \tag{S15}$$

The PPMI score evaluates the pairing of residues by considering each residue as a *word* and their pairing as *co-occurrence*. This score compares the frequency of interacting pairs to the expected frequency of random associations. Consequently, a high positive value indicates a strong association between an interacting pair. In contrast, zero values correspond to associations that only occur as often as expected by random chance or less.

$$\mathrm{PMI}_{\mathrm{pair}} = \log_2\left(\frac{\mathrm{Frequency}_{\mathrm{pair}}}{\mathrm{Frequency}_{\mathrm{residue\ 1}} \times \mathrm{Frequency}_{\mathrm{residue\ 2}}}\right) \tag{S16}$$

$$\mathrm{PPMI}_{\mathrm{pair}} = \max(\mathrm{PMI}_{\mathrm{pair}}, 0) \tag{S17}$$

We computed score values for each pair in each protein region for both transient and stable interfaces. Subsequently, all pairs with tf-idf values greater than 1 were removed from the PPMI table, as their representation is limited across the dataset, making their interpretation unreliable due to insufficient structural data.

**Results** Table S5 show that there are 13 tf-idf values exceeding 1, indicating that the corresponding pairs only occur in a limited set of protein complex structures. Notably, these pairs are predominantly located in the rim, which can be attributed to a high variability in pair composition within this region (see also SI section L). In contrast, with the exception of the negative–negative pair, all pair types are widely represented in the core & support regions of the stable interface sub-dataset, whereas the negative-positive pair is not fully represented in the core & support region of the transient interface sub-dataset.

| | Interior | Core & Support transient | Core & Support stable | Rim transient | Rim stable |
|---|---|---|---|---|---|
| apolar-apolar | 0.031 | 0.078 | 0.105 | 1.215 | 1.732 |
| apolar-negative | 0.196 | 0.629 | 0.519 | 1.019 | 1.231 |
| apolar-polar | 0.036 | 0.080 | 0.115 | 0.306 | 0.574 |
| apolar-positive | 0.108 | 0.234 | 0.130 | 0.575 | 0.539 |
| negative-negative | 1.327 | 1.461 | 1.149 | 1.423 | 1.284 |
| negative-polar | 0.295 | 0.803 | 0.651 | 0.834 | 1.092 |
| negative-positive | 0.779 | 1.017 | 0.838 | 1.119 | 1.269 |
| polar-polar | 0.108 | 0.359 | 0.132 | 0.982 | 0.856 |
| polar-positive | 0.177 | 0.350 | 0.180 | 0.620 | 0.742 |
| positive-positive | 0.676 | 0.860 | 0.605 | 0.656 | 0.884 |

Table S5: td-idf values for each pair properties in the different protein regions.

| | Interior | Core & Support transient | Core & Support stable | Rim transient | Rim stable |
|---|---|---|---|---|---|
| apolar-apolar | 0.000 | 0.743 | 0.292 | 0.000 | 0.000 |
| apolar-negative | 1.553 | 0.986 | 1.213 | 1.364 | 1.332 |
| apolar-polar | 0.924 | 1.270 | 1.191 | 0.583 | 0.626 |
| apolar-positive | 1.411 | 1.137 | 1.099 | 1.007 | 1.303 |
| negative-negative | 1.912 | 0.106 | 1.042 | 2.557 | 3.086 |
| negative-polar | 1.433 | 0.143 | 0.864 | 1.285 | 1.446 |
| negative-positive | 1.563 | 0.000 | 0.629 | 1.523 | 1.901 |
| polar-polar | 0.000 | 0.000 | 0.000 | 0.000 | 0.000 |
| polar-positive | 1.029 | 0.000 | 0.421 | 0.706 | 0.761 |
| positive-positive | 1.338 | 0.064 | 0.246 | 2.163 | 2.030 |

Table S6: Raw PPMI values computed for pair properties in the different protein regions.

### N List of used structures

Temperature is set to 25.0 C and pH is set to 7.2 when unknown.

| PDB | Database origin | Protein family | PubMed | Classification | Organism | Mutation | Method | Res. (Å) | UniprotKB | Affinity type | Affinity (M) | Chain 1 | Chain 2 | Temp. (C) | pH |
|---|---|---|---|---|---|---|---|---|---|---|---|---|---|---|---|
| 1acb | BM5.5 PPI4DOC PDBind | PF00089 PF00280 | 1583684 | hydrolase / hydrolase inhibitor | Bos taurus Hirudo medicinalis | No | x-ray diffraction | 2.0 | P00766 P01051 | Kd | 2e-10 | E | I | 25.0 | 7.2 |
| 1axi | PPI4DOC PDBind | PF00103 PF09067 PF00041 | 9353194 | complex (hormone / receptor) | Homo sapiens | Yes | x-ray diffraction | 2.1 | P01241 P10912 | Kd | 1.4e-08 | A | B | 25.0 | 7.2 |
| 1ay7 | BM5.5 PPI4DOC PDBind | PF00545 PF01337 | 9757110 | complex (enzyme / inhibitor) | Kitasatospora aureofaciens Bacillus amyloliquefaciens | No | x-ray diffraction | 1.7 | P05798 P11540 | Kd | 1e-06 | A | B | 25.0 | 7.2 |
| 1dfj | BM5.5 PDBind | PF00074 PF13516 PF18779 | 7877692 | complex (endonuclease / inhibitor) | Bos taurus Sus scrofa | No | x-ray diffraction | 2.5 | P61823 P10775 | Ki | 5.9e-14 | E | I | 25.0 | 6.0 |
| 1e6e | BM5.5 | PF07992 PF00111 | 11053423 | oxidoreductase | Bos taurus | Yes | x-ray diffraction | 2.3 | P08165 P00257 | Kd | 8.6e-07 | A | B | 25.0 | 7.4 |
| 1e96 | BM5.5 PPI4DOC PDBind | PF00071 PF13181 | 11090627 | signaling protein | Homo sapiens | Yes | x-ray diffraction | 2.4 | P63000 P19878 | Kd | 2.7e-06 | A | B | 18.0 | 7.0 |
| 1eer | BM5.5 | PF00758 PF09067 PF00041 | 9774108 | complex (cytokine / receptor) | Homo sapiens | Yes | x-ray diffraction | 1.9 | P01588 P19235 | Kd | 1e-09 | A | BC | 25.0 | 7.2 |
| 1efn | BM5.5 PPI4DOC | PF00018 PF00469 | 8681387 | complex (sh3 domain / viral enhancer) | Homo sapiens Human immunodeficiency virus 1 | Yes | x-ray diffraction | 2.5 | P06241 P03406 | Kd | 3.8e-07 | B | A | 25.0 | 7.2 |
| 1ewy | BM5.5 PPI4DOC | PF00175 PF00111 | 11053838 | oxidoreductase | Nostoc sp. PCC 7119 | No | x-ray diffraction | 2.38 | P21890 P0A3C8 | Kd | 3.57e-06 | A | C | 25.0 | 7.2 |
| 1f34 | BM5.5 PPI4DOC PDBind | PF00026 | 10932249 | hydrolase / hydrolase inhibitor | Sus scrofa Ascaris suum | Yes | x-ray diffraction | 2.45 | P00791 P19400 | Kd | 1e-10 | A | B | 25.0 | 7.2 |
| 1f3v | 3DComplexV6 PDBind | PF09034 PF21355 | 10892748 | apoptosis | Homo sapiens | No | x-ray diffraction | 2.0 | Q15628 Q12933 | Kd | 7.8e-06 | 0 | 1 | 25.0 | 7.2 |
| 1fle | BM5.5 PPI4DOC PDBind | PF00089 PF00095 PF10511 | 8794736 | complex (serine protease / inhibitor) | Sus scrofa Homo sapiens | No | x-ray diffraction | 1.9 | P00772 P19957 | Ki | 6e-09 | E | I | 25.0 | 7.2 |
| 1fqj | BM5.5 PPI4DOC | PF00503 PF00615 PF00631 PF04868 | 11234020 | signaling protein | Bos taurus Rattus norvegicus | No | x-ray diffraction | 2.02 | P10824 P04695 O46469 P04972 | Kd | 6.7e-08 | A | B | 25.0 | 7.2 |
| 1fy8 | PDBind | PF00014 | 11420435 | hydrolase / hydrolase inhibitor | Rattus rattus Bos taurus | No | x-ray diffraction | 1.7 | P00763 P00974 | Ki | 9e-06 | 0 | 1 | 25.0 | 7.2 |
| 1g9i | PDBind | PF00089 | 11257512 | hydrolase / hydrolase inhibitor | Bos taurus Vigna radiata var. radiata | Yes | x-ray diffraction | 2.2 | P00760 P01062 | Kd | 1.2e-07 | 0 | 1 | 25.0 | 7.2 |
| 1gcq | BM5.5 | PF00018 PF00018 | 11406576 | signaling protein / signaling protein | Homo sapiens Mus musculus | No | x-ray diffraction | 1.68 | P62993 P27870 | Kd | 1.68e-05 | B | C | 25.0 | 7.4 |

| PDB | Database origin | Protein family | PubMed | Classification | Organism | Mutation | Method | Res. (Å) | UniprotKB | Affinity type | Affinity (M) | Chain 1 | Chain 2 | Temp. (C) | pH |
|---|---|---|---|---|---|---|---|---|---|---|---|---|---|---|---|
| 1grn | BM5.5<br>PDBind | PF00071<br>PF00620 | 9846874 | gene regulation | Homo sapiens | No | x-ray diffraction | 2.1 | P60953<br>Q07960 | Kd | 3.88e-07 | A | B | 25.0 | 8.0 |
| 1h59 | PDBind | PF00049<br>PF00219 | 11447105 | insulin | Homo sapiens | No | x-ray diffraction | 2.1 | P05019<br>P24593 | Kd | 3.7e-08 | 0 | 1 | 25.0 | 7.2 |
| 1hia | BM5.5 | PF00089<br>PF00089<br>PF02822 | 9032072 | complex (protease / inhibitor) | Sus scrofa Hirudo medicinalis | No | x-ray diffraction | 2.4 | P00752<br>P00752<br>P80302 | Ki | 1.3e-08 | AB | I | 25.0 | 7.2 |
| 1i2m | BM5.5<br>PPI4DOC | PF00071<br>PF00415 | 11336674 | cell cycle | Homo sapiens | No | x-ray diffraction | 1.76 | P62826<br>P18754 | Kd | 2.5e-12 | A | B | 25.0 | 7.2 |
| 1i8k | PPI4DOC | PF07686<br>PF07686 | 11352579 | immune system | Mus musculus | Yes | x-ray diffraction | 1.8 | P01660<br>P18529 | Kd | 2.2e-08 | A | B | 25.0 | 7.2 |
| 1jiw | BM5.5<br>PDBind | PF13583<br>PF00353<br>PF08548<br>PF02974 | 11445573 | hydrolase / hydrolase inhibitor | Pseudomonas aeruginosa | No | x-ray diffraction | 1.74 | Q03023<br>Q03026 | Kd | 4e-12 | P | I | 25.0 | 7.2 |
| 1jmo | BM5.5 | PF09396<br>PF00089<br>PF00079 | 12169660 | blood clotting | Homo sapiens | Yes | x-ray diffraction | 2.2 | P00734<br>P00734<br>P05546 | Kd | 1.15e-07 | A | HL | 25.0 | 7.2 |
| 1jps | BM5.5<br>PPI4DOC | PF09294 | 11601848 | immune system | Homo sapiens | No | x-ray diffraction | 1.85 | P13726 | Kd | 1e-10 | HL | T | 25.0 | 7.4 |
| 1jtd | BM5.5<br>PPI4DOC<br>PDBind | PF13354<br>PF00144<br>PF13540 | 11573088 | hydrolase / inhibitor | Escherichia coli Streptomyces exfoliatus | No | x-ray diffraction | 2.3 | P62593<br>O87916 | Ki | 2.72e-11 | B | A | 25.0 | 7.0 |
| 1ktz | BM5.5<br>PPI4DOC<br>PDBind | PF00019<br>PF08917 | 11850637 | cytokine / cytokine receptor | Homo sapiens | No | x-ray diffraction | 2.15 | P10600<br>P37173 | Kd | 2.9e-07 | A | B | 25.0 | 7.2 |
| 1l4d | PPI4DOC<br>PDBind | PF00089<br>PF02821 | 12456874 | hydrolase / hydrolase activator | Homo sapiens Streptococcus dysgalactiae subsp. equisimilis | Yes | x-ray diffraction | 2.3 | P00747<br>P00779 | Kd | 1.966e-07 | A | B | 25.0 | 7.2 |
| 1lp1 | PDBind | PF02216 | 12604795 | immune system | Staphylococcus aureus | Yes | x-ray diffraction | 2.3 | P38507 | Kd | 2e-06 | 0 | 1 | 25.0 | 7.2 |
| 1lw6 | PDBind | PF00082<br>PF00280 | 12142461 | hydrolase | Bacillus amyloliquefaciens Hordeum vulgare | Yes | x-ray diffraction | 1.5 | P00782<br>P01053 | Kd | 2e-12 | 0 | 1 | 25.0 | 7.2 |
| 1m1e | PDBind | PF00514<br>PF06384 | 12408825 | structural protein | Mus musculus Homo sapiens | No | x-ray diffraction | 2.1 | Q02248<br>Q9NSA3 | Kd | 1e-09 | 0 | 1 | 25.0 | 7.2 |
| 1mzw | PDBind | PF00160<br>PF08799 | 12875835 | isomerase | Homo sapiens | No | x-ray diffraction | 2.0 | O43447<br>O43172 | Kd | 1.97e-06 | 0 | 1 | 25.0 | 7.2 |
| 1nb5 | PPI4DOC | PF00112<br>PF00031 | 12581647 | hydrolase / hydrolase inhibitor | Sus scrofa Homo sapiens | No | x-ray diffraction | 2.4 | O46427<br>O46427<br>P01040 | Ki | 6.9e-11 | A | C | 25.0 | 7.2 |

| PDB | Database origin | Protein family | PubMed | Classification | Organism | Mutation | Method | Res. (Å) | UniprotKB | Affinity type | Affinity (M) | Chain 1 | Chain 2 | Temp. (C) | pH |
|---|---|---|---|---|---|---|---|---|---|---|---|---|---|---|---|
| 1nca | PPI4DOC | PF00064 PF07654 | 1381757 | hydrolase(o-glycosyl) | Influenza A virus (A / tern / Australia / G70C / 1975(H11N9)) Mus musculus | No | x-ray diffraction | 2.5 | P03472 P01865 | Kd | 8.3e-09 | HL | N | 25.0 | 7.2 |
| 1nw9 | BM5.5 PDBind | PF00653 PF00656 | 12620238 | apoptosis | Homo sapiens | No | x-ray diffraction | 2.4 | P98170 P55211 | Ki | 1.3e-08 | B | A | 25.0 | 7.2 |
| 1oc0 | BM5.5 PPI4DOC PDBind | PF00079 PF01033 | 12808446 | hydrolase / inhibitor | Homo sapiens | Yes | x-ray diffraction | 2.28 | P05121 P04004 | Kd | 1e-09 | A | B | 25.0 | 7.2 |
| 1op9 | PDBind | PF00062 | 12917687 | hydrolase | Camelus dromedarius Homo sapiens | No | x-ray diffraction | 1.86 | P61626 | Kd | 7e-10 | 0 | 1 | 25.0 | 7.2 |
| 1oyv | BM5.5 PPI4DOC | PF00082 PF02428 | 12684499 | hydrolase | Bacillus licheniformis Solanum lycopersicum | No | x-ray diffraction | 2.5 | P00780 P05119 | Ki | 9e-09 | B | I | 25.0 | 7.2 |
| 1pk1 | PPI4DOC | PF07647 PF00536 | 15905166 | transcription repression | Drosophila melanogaster | Yes | x-ray diffraction | 1.8 | P39769 Q9VHA0 | Kd | 5.4e-08 | A | B | 25.0 | 7.2 |
| 1pvh | BM5.5 PPI4DOC | PF00041 PF09240 PF01291 | 14527405 | signaling protein / cytokine | Homo sapiens | No | x-ray diffraction | 2.5 | P40189 P15018 | Kd | 8e-08 | A | B | 25.0 | 7.2 |
| 1pxv | BM5.5 PPI4DOC | PF05543 PF09023 | 12874290 | hydrolase | Staphylococcus aureus | Yes | x-ray diffraction | 1.8 | P0C1S6 Q9EYW6 | Ki | 3.1e-10 | A | C | 25.0 | 7.2 |
| 1r6q | BM5.5 | PF02861 PF02617 | 15037248 | chaperone / protein binding | Escherichia coli | No | x-ray diffraction | 2.35 | P0ABH9 P0A8Q6 | Kd | 3.3e-07 | A | C | 25.0 | 7.2 |
| 1rjc | PPI4DOC PDBind | PF00062 | 15659390 | immune system / hydrolase | Camelus dromedarius Gallus gallus | No | x-ray diffraction | 1.4 | P00698 | Kd | 7.7e-11 | A | B | 25.0 | 7.2 |
| 1rv6 | BM5.5 PPI4DOC | PF00341 PF21339 | 14684734 | hormone / growth factor / receptor | Homo sapiens | No | x-ray diffraction | 2.45 | P49763 P17948 | IC50 | 2.75e-07 | VW | X | 25.0 | 7.2 |
| 1sbb | BM5.5 | PF07654 PF01123 PF02876 | 9881971 | immune system | Mus musculus Staphylococcus aureus | Yes | x-ray diffraction | 2.4 | P01852 P01552 | Kd | 0.00014 | A | B | 25.0 | 7.2 |
| 1sv0 | PPI4DOC | PF02198 PF02198 | 15260987 | transcription | Drosophila melanogaster | Yes | x-ray diffraction | 2.07 | Q01842 Q7K119 | Kd | 1.11e-08 | A | B | 25.0 | 7.2 |
| 1t0p | PPI4DOC PDBind | PF00092 PF03921 | 15728350 | immune system | Homo sapiens | Yes | x-ray diffraction | 1.66 | P20701 P32942 | Kd | 2.5e-05 | A | B | 25.0 | 7.2 |
| 1t5z | PDBind | PF00104 PF12489 | 15563469 | hormone / growth factor | Homo sapiens | No | x-ray diffraction | 2.3 | P10275 Q13772 | Kd | 3.3e-05 | 0 | 1 | 25.0 | 7.2 |
| 1t6b | BM5.5 PPI4DOC 3DComplexV6 PDBind | PF07691 PF17475 PF03495 PF17476 PF20835 PF00092 | 15243628 | membrane protein / toxin | Bacillus anthracis Homo sapiens | No | x-ray diffraction | 2.5 | P13423 P58335 | Kd | 4e-10 | X | Y | 25.0 | 7.4 |

| PDB | Database origin | Protein family | PubMed | Classification | Organism | Mutation | Method | Res. (Å) | UniprotKB | Affinity type | Affinity (M) | Chain 1 | Chain 2 | Temp. (C) | pH |
|---|---|---|---|---|---|---|---|---|---|---|---|---|---|---|---|
| 1ta3 | PPI4DOC PDBind | PF00704 PF00331 | 15181003 | hydrolase inhibitor / hydrolase | Triticum aestivum Aspergillus nidulans | No | x-ray diffraction | 1.7 | Q8L5C6 Q00177 | Ki | 9e-09 | A | B | 25.0 | 7.2 |
| 1u0s | PPI4DOC PDBind | PF00072 PF07194 | 15289606 | signaling protein | Thermotoga maritima | No | x-ray diffraction | 1.9 | Q56312 Q56310 | Kd | 2.3e-07 | A | Y | 25.0 | 7.2 |
| 1uad | PPI4DOC | PF00071 PF01833 | 12839989 | endocytosis / exocytosis | Homo sapiens Rattus norvegicus | No | x-ray diffraction | 2.1 | P11233 O54921 | Kd | 1.37e-07 | A | B | 25.0 | 7.2 |
| 1v7p | PPI4DOC | PF00059 PF00059 PF00092 | 15276841 | toxin / cell adhesion | Echis multisquamatus Homo sapiens | Yes | x-ray diffraction | 1.9 | Q7T2Q1 Q7T2Q0 P17301 | IC50 | 6e-09 | A | C | 25.0 | 7.2 |
| 1veu | PDBind | PF08923 PF03259 | 15263099 | signaling protein / protein binding | Mus musculus | Yes | x-ray diffraction | 2.15 | O88653 Q9JHS3 | Kd | 1.28e-08 | 0 | 1 | 25.0 | 7.2 |
| 1vg0 | PPI4DOC PDBind | PF00996 PF00071 | 15186776 | protein binding / protein transport | Rattus norvegicus | Yes | x-ray diffraction | 2.2 | P37727 P09527 | Kd | 5e-09 | A | B | 25.0 | 7.2 |
| 1vrk | PDBind | | 10194305 | complex(calcium-binding protein / peptide) | synthetic construct Gallus gallus | No | x-ray diffraction | 1.9 | P11799 | Kd | 6.8e-09 | 0 | 1 | 25.0 | 7.2 |
| 1wej | BM5.5 PPI4DOC | PF07686 PF00034 | 9698550 | complex (antibody / electron transport) | Mus musculus Equus caballus | No | x-ray diffraction | 1.8 | P01635 P00004 | Kd | 6e-08 | HL | F | 25.0 | 7.2 |
| 1wlp | PDBind | PF05038 PF00018 | 16326715 | oxidoreductase / signaling protein | Homo sapiens | No | solution nmr | | P13498 P14598 | Kd | 6.4e-07 | 0 | 1 | 25.0 | 7.2 |
| 1xg2 | PPI4DOC 3DComplexV6 PDBind | PF01095 PF04043 | 15722470 | hydrolase / hydrolase inhibitor | Solanum lycopersicum Actinidia chinensis | No | x-ray diffraction | 1.9 | P14280 P83326 | Kd | 5e-09 | A | B | 25.0 | 7.2 |
| 1xt9 | PDBind | PF02902 PF00240 | 15567417 | hydrolase / hydrolase inhibitor | Homo sapiens | No | x-ray diffraction | 2.2 | Q96LD8 Q15843 | Kd | 2e-07 | 0 | 1 | 25.0 | 7.2 |
| 1z0k | BM5.5 | PF00071 PF11464 | 16034420 | protein transport | Homo sapiens | Yes | x-ray diffraction | 1.92 | P20338 Q9H1K0 | Kd | 7.7e-06 | A | B | 25.0 | 7.5 |
| 1z7x | PPI4DOC | PF00074 PF13516 PF18779 | 17350650 | hydrolase / hydrolase inhibitor | Homo sapiens | No | x-ray diffraction | 1.95 | P07998 P13489 | Kd | 2.9e-16 | Y | Z | 25.0 | 7.2 |
| 1ze3 | PPI4DOC | PF02753 PF00345 PF00419 PF13954 | 15920478 | chaperone / structural / membrane protein | Escherichia coli | No | x-ray diffraction | 1.84 | P31697 P08191 P30130 | Kd | 1.2e-06 | C | H | 25.0 | 7.2 |
| 1zli | BM5.5 PDBind | PF00246 PF10468 | 15961103 | hydrolase / hydrolase inhibitor | Homo sapiens Rhipicephalus bursa | No | x-ray diffraction | 2.09 | P15086 Q5EPH2 | Ki | 1.3e-09 | A | B | 25.0 | 7.2 |
| 2a7u | PDBind | PF00213 | 16128580 | hydrolase | Escherichia coli O157:H7 | No | solution nmr | | P0ABB2 P0ABA5 | Kd | 1.2e-07 | 0 | 1 | 25.0 | 7.2 |
| 2arp | PDBind | PF00019 PF09289 PF07648 | 16482217 | hormone / growth factor | Homo sapiens Rattus norvegicus | No | x-ray diffraction | 2.0 | P08476 P21674 | Kd | 4.3e-07 | 0 | 1 | 25.0 | 7.2 |
| 2ast | PPI4DOC | PF03931 PF01466 PF12937 PF01111 | 16209941 | ligase / ligase inhibitor | Homo sapiens | Yes | x-ray diffraction | 2.3 | P63208 Q13309 P61024 P46527 | Kd | 7e-06 | C | B | 25.0 | 7.2 |

| PDB | Database origin | Protein family | PubMed | Classification | Organism | Mutation | Method | Res. (Å) | UniprotKB | Affinity type | Affinity (M) | Chain 1 | Chain 2 | Temp. (C) | pH |
|---|---|---|---|---|---|---|---|---|---|---|---|---|---|---|---|
| 2b42 | BM5.5 PDBind | PF14541 PF14543 PF00457 | 19769747 | hydrolase inhibitor / hydrolase | Triticum aestivum Bacillus subtilis | No | x-ray diffraction | 2.5 | Q8H0K8 P18429 | Kd | 1.07e-09 | B | A | 22.0 | 5.0 |
| 2b7c | PDBind | PF00009 PF03143 PF03144 PF00736 | 16675455 | translation | Saccharomyces cerevisiae | Yes | x-ray diffraction | 1.8 | P02994 P32471 | Kd | 4e-07 | 0 | 1 | 25.0 | 7.2 |
| 2c0l | BM5.5 PPI4DOC PDBind | PF13181 PF02036 | 17157249 | transport protein / receptor | Homo sapiens | No | x-ray diffraction | 2.3 | P50542 P22307 | Kd | 1.09e-07 | A | B | 35.0 | 7.4 |
| 2c1m | PDBind | PF00514 PF01749 PF16186 PF08911 | 16222336 | protein transport / membrane protein | Mus musculus | No | x-ray diffraction | 2.2 | P52293 Q9JIH2 | Kd | 1.1e-09 | 0 | 1 | 25.0 | 7.2 |
| 2cjs | PPI4DOC | PF00168 PF02318 | 16732694 | exocytosis | Rattus norvegicus | Yes | x-ray diffraction | 1.78 | Q62768 Q9JIS1 | Kd | 1e-07 | B | C | 25.0 | 7.2 |
| 2dd8 | BM5.5 PPI4DOC | PF07686 PF07654 PF16451 PF09408 | 16597622 | immune system / viral protein | Homo sapiens Severe acute respiratory syndrome-related coronavirus | No | x-ray diffraction | 2.3 | Q8N355 P59594 | Kd | 2e-08 | HL | S | 25.0 | 7.2 |
| 2dsp | PDBind | PF00219 PF00049 | 16924115 | protein binding / hormone / growth factor | Homo sapiens | No | x-ray diffraction | 2.5 | P22692 Q9NP10 | Kd | 3e-06 | 0 | 1 | 25.0 | 7.2 |
| 2es4 | 3DComplexV6 | PF00561 PF03280 | 16518399 | hydrolase | Burkholderia glumae | Yes | x-ray diffraction | 1.85 | P0DUB8 Q05490 | Kd | 5e-09 | A | B | 25.0 | 7.2 |
| 2f31 | PDBind | PF06371 PF06367 | 16472745 | structural protein | Mus musculus | No | x-ray diffraction | 2.1 | Q3US76 O08808 | Kd | 1.02e-07 | 0 | 1 | 25.0 | 7.2 |
| 2f4m | PDBind | PF01841 PF09280 | 16500903 | hydrolase | Mus musculus | No | x-ray diffraction | 1.85 | Q9JI78 P54728 | Kd | 6.5e-08 | 0 | 1 | 25.0 | 7.2 |
| 2f5z | PPI4DOC | PF07992 PF02852 PF02817 | 16442803 | oxidoreductase / protein binding | Homo sapiens | Yes | x-ray diffraction | 2.18 | P09622 O00330 | Kd | 7.8e-10 | B | C | 25.0 | 7.2 |
| 2f9z | 3DComplexV6 | PF04509 PF03975 | 16469702 | signaling protein | Thermotoga maritima MSB8 | No | x-ray diffraction | 2.4 | Q9X006 Q9X005 | Kd | 9e-07 | A | C | 25.0 | 7.2 |
| 2fdb | PPI4DOC | PF00167 PF13927 PF07679 | 16384934 | hormone / growth factor / transferase | Homo sapiens | Yes | x-ray diffraction | 2.28 | P55075 P21802 | Kd | 1.55e-07 | M | P | 25.0 | 7.2 |
| 2few | PDBind | PF00359 PF02302 | 16443929 | transferase | Escherichia coli Escherichia coli O157:H7 | Yes | solution nmr | | P00550 P00550 | Kd | 0.0037 | 0 | 1 | 25.0 | 7.2 |
| 2fu5 | PPI4DOC | PF04421 PF00071 | 16541104 | signaling protein | Homo sapiens Mus musculus | No | x-ray diffraction | 2.0 | P47224 P55258 | Kd | 7e-10 | A | B | 25.0 | 7.2 |
| 2fyl | PDBind | PF06400 PF00057 | 16938309 | surface active protein | Homo sapiens | No | solution nmr | | P30533 Q07954 | Kd | 2.8e-06 | 0 | 1 | 25.0 | 7.2 |
| 2gng | PDBind | PF00069 PF02827 | 16699172 | transferase / transferase inhibitor | Bos taurus Mus musculus | Yes | x-ray diffraction | 1.87 | P00517 P63248 | IC50 | 6e-06 | 0 | 1 | 25.0 | 7.2 |

| PDB | Database origin | Protein family | PubMed | Classification | Organism | Mutation | Method | Res. (Å) | UniprotKB | Affinity type | Affinity (M) | Chain 1 | Chain 2 | Temp. (C) | pH |
|---|---|---|---|---|---|---|---|---|---|---|---|---|---|---|---|
| 2hev | PPI4DOC PDBind | PF00020 | 16905106 | cytokine | Homo sapiens | Yes | x-ray diffraction | 2.41 | P23510 P43489 | IC50 | 6.2e-08 | R | F | 25.0 | 7.2 |
| 2hqs | BM5.5 PPI4DOC | PF04052 PF07676 PF00691 | 17375930 | transport protein / lipoprotein | Escherichia coli | No | x-ray diffraction | 1.5 | P0A855 P0A912 | Kd | 2.7e-08 | A | H | 25.0 | 7.2 |
| 2hqw | PDBind | PF13499 | 18073110 | metal binding protein | Rattus norvegicus | No | x-ray diffraction | 1.9 | P0DP29 P35439 | Kd | 2e-09 | 0 | 1 | 25.0 | 7.2 |
| 2hrk | BM5.5 PPI4DOC PDBind | | 16914447 | ligase / rna binding protein | Saccharomyces cerevisiae | No | x-ray diffraction | 2.05 | P46655 P46672 | Kd | 9e-09 | A | B | 25.0 | 7.2 |
| 2ij0 | PPI4DOC | PF07686 | 17268555 | protein binding | Staphylococcus aureus Homo sapiens | No | x-ray diffraction | 2.25 | P06886 A0A5B4 | Kd | 1.8e-10 | A | B | 25.0 | 7.2 |
| 2j1k | PPI4DOC | PF07686 | 16923808 | virus / receptor | Homo sapiens Canine adenovirus 2 | No | x-ray diffraction | 2.3 | P78310 Q65914 | Kd | 1.1e-09 | F | C | 25.0 | 7.2 |
| 2j4w | PPI4DOC | PF02430 PF07686 PF07654 PF07686 PF07654 | 17229439 | immune system | Plasmodium vivax Mus musculus | Yes | x-ray diffraction | 2.5 | Q9TY14 Q569B4 Q5XFY8 | Kd | 6e-10 | HL | D | 25.0 | 7.2 |
| 2j59 | PPI4DOC | PF00025 PF00169 | 17347647 | hydrolase | Mus musculus Homo sapiens | Yes | x-ray diffraction | 2.1 | P84078 Q5T5U3 | Kd | 5.5e-08 | A | B | 25.0 | 7.2 |
| 2j7p | BM5.5 | PF00448 PF02881 PF00448 | 17184999 | signal recognition | Thermus aquaticus | No | x-ray diffraction | 1.97 | O07347 P83749 | Kd | 1e-08 | A | D | 25.0 | 7.2 |
| 2j8x | PPI4DOC | PF03167 PF18880 | 17157317 | hydrolase / inhibitor | human gammaherpesvirus 4 Bacillus phage PBS2 | No | x-ray diffraction | 2.3 | P12888 P14739 | Ki | 8e-09 | A | B | 25.0 | 7.2 |
| 2jby | PDBind | PF11099 PF00452 | 17386268 | apoptosis | Myxoma virus Homo sapiens | No | x-ray diffraction | 2.41 | Q85295 Q16611 | IC50 | 5e-08 | 0 | 1 | 25.0 | 7.2 |
| 2jod | PDBind | PF02793 PF00123 | 17470806 | signaling protein | Homo sapiens | Yes | solution nmr | | P41586 P18509 | Ki | 3.5e-07 | 0 | 1 | 25.0 | 7.2 |
| 2k2s | PDBind | PF11476 PF12661 | 18818666 | cell adhesion | Toxoplasma gondii | No | solution nmr | | O00834 Q9XYH7 | Kd | 5.3e-08 | 0 | 1 | 25.0 | 7.2 |
| 2k79 | PDBind | PF00018 PF00017 | 19361414 | transferase | Mus musculus | No | solution nmr | | Q03526 Q03526 | Kd | 0.00067 | 0 | 1 | 25.0 | 7.2 |
| 2mp0 | PDBind | PF00391 PF05524 PF00358 | 25131700 | transferase | Escherichia coli K-12 | No | solution nmr | | P08839 P69783 | Kd | 0.025 | 0 | 1 | 25.0 | 7.2 |
| 2o3b | BM5.5 PDBind | PF01223 PF07924 | 17138564 | hydrolase / hydrolase inhibitor | Nostoc sp. PCC 7120 = FACHB-418 | Yes | x-ray diffraction | 2.3 | P38446 Q7A260 | Ki | 3.2e-12 | A | B | 25.0 | 7.2 |
| 2omu | PDBind | PF00028 | 17715295 | cell invasion / cell adhesion | Listeria monocytogenes EGD-e Homo sapiens | Yes | x-ray diffraction | 1.8 | P12830 | Kd | 6e-10 | 0 | 1 | 25.0 | 7.2 |
| 2oob | BM5.5 PPI4DOC PDBind | PF00240 | 17679095 | ligase | Homo sapiens Bos taurus | No | x-ray diffraction | 1.9 | Q13191 P0CH28 | Kd | 5.7e-05 | A | B | 25.0 | 7.0 |

| PDB | Database origin | Protein family | PubMed | Classification | Organism | Mutation | Method | Res. (Å) | UniprotKB | Affinity type | Affinity (M) | Chain 1 | Chain 2 | Temp. (C) | pH |
|---|---|---|---|---|---|---|---|---|---|---|---|---|---|---|---|
| 2oor | BM5.5 | PF05222 PF01262 PF02233 | 17323922 | oxidoreductase | Rhodospirillum rubrum | No | x-ray diffraction | 2.32 | Q2RSB2 P0C188 | Kd | 1.55e-08 | AB | C | 25.0 | 7.2 |
| 2ot3 | BM5.5 PPI4DOC PDBind | PF18151 PF02204 PF00071 | 17450153 | protein transport | Homo sapiens | No | x-ray diffraction | 2.1 | Q9UJ41 Q9UL25 | Kd | 1.8e-06 | B | A | 25.0 | 7.2 |
| 2p8q | PDBind | PF00514 PF03810 PF11538 | 18187419 | protein transport | Homo sapiens | No | x-ray diffraction | 2.35 | Q14974 O95149 | Kd | 1.56e-08 | 0 | 1 | 25.0 | 7.2 |
| 2pcc | BM5.5 | PF00141 PF00034 | 1334573 | oxidoreductase / electron transport | Saccharomyces cerevisiae | No | x-ray diffraction | 2.3 | P00431 P00044 | Kd | 1.6e-06 | A | B | 25.0 | 7.2 |
| 2ptt | PPI4DOC PDBind | PF07686 PF11465 | 17950006 | immune system | Mus musculus | Yes | x-ray diffraction | 1.63 | P18181 Q07763 | Kd | 4e-06 | A | B | 25.0 | 7.2 |
| 2q0o | PPI4DOC | PF03472 PF00196 PF09228 | 17921255 | transcription | Sinorhizobium fredii NGR234 | No | x-ray diffraction | 2.0 | P55407 P55408 | Kd | 1.49e-08 | A | C | 25.0 | 7.2 |
| 2qc1 | PPI4DOC | PF02931 | 17643119 | protein binding | Bungarus multicinctus Mus musculus | Yes | x-ray diffraction | 1.94 | P60616 P04756 | Kd | 1e-11 | A | B | 25.0 | 7.2 |
| 2qxv | PDBind | PF00400 PF11616 | 17937919 | gene regulation | Mus musculus | No | x-ray diffraction | 1.82 | Q921E6 Q61188 | Kd | 3.8e-07 | 0 | 1 | 25.0 | 7.2 |
| 2uyz | PPI4DOC PDBind | PF00179 PF11976 | 17491593 | ligase | Mus musculus Homo sapiens | Yes | x-ray diffraction | 1.4 | P63280 P63165 | Kd | 8.2e-08 | A | B | 25.0 | 7.2 |
| 2v3b | PPI4DOC PDBind | PF07992 PF18113 PF00301 | 17636129 | oxidoreductase | Pseudomonas aeruginosa PAO1 | No | x-ray diffraction | 2.45 | Q9HTK9 Q9HTK8 | Kd | 5e-06 | A | B | 25.0 | 7.2 |
| 2v5q | PPI4DOC | PF00069 | 18391401 | transferase | Homo sapiens synthetic construct | No | x-ray diffraction | 2.3 | P53350 | Kd | 7.1e-08 | A | B | 25.0 | 7.2 |
| 2v6x | PDBind | PF04212 PF03357 | 17928861 | protein transport | Saccharomyces cerevisiae | No | x-ray diffraction | 1.98 | P52917 P36108 | Kd | 2.8e-05 | 0 | 1 | 25.0 | 7.2 |
| 2v8s | PPI4DOC PDBind | PF01417 PF05008 | 18033301 | protein transport | Homo sapiens | No | x-ray diffraction | 2.22 | Q14677 Q9UEU0 | Kd | 2.2e-05 | E | V | 25.0 | 7.2 |
| 2ver | PDBind | PF04619 PF07686 | 18086185 | cell adhesion | Escherichia coli Homo sapiens | No | solution nmr | | Q57254 P06731 | Kd | 1.31e-05 | 0 | 1 | 25.0 | 7.2 |
| 2vln | PDBind | PF01320 PF21431 | 18471830 | protein binding | Escherichia coli | Yes | x-ray diffraction | 1.6 | P13479 P09883 | Kd | 1.68e-12 | 0 | 1 | 25.0 | 7.2 |
| 2vog | PDBind | PF00452 PF15185 | 18462686 | apoptosis | Mus musculus | Yes | x-ray diffraction | 1.9 | Q07440 Q91ZE9 | Kd | 2.1e-07 | 0 | 1 | 25.0 | 7.2 |
| 2w2x | PPI4DOC | PF00071 PF00018 PF00018 | 19394299 | signaling protein / hydrolase | Homo sapiens | Yes | x-ray diffraction | 2.3 | P15153 P16885 P16885 | Kd | 2.19e-05 | A | B | 25.0 | 7.2 |
| 2wel | PDBind | PF00069 PF13499 | 20668654 | transferase | Homo sapiens | No | x-ray diffraction | 1.9 | Q13557 P0DP23 | Kd | 3e-07 | 0 | 1 | 25.0 | 7.2 |
| 2wwk | PDBind | PF07679 PF07679 | 20133654 | transferase / structural protein | Homo sapiens | Yes | x-ray diffraction | 1.7 | O75147 Q8WZ42 | Kd | 1.28e-06 | 0 | 1 | 25.0 | 7.2 |

| PDB | Database origin | Protein family | PubMed | Classification | Organism | Mutation | Method | Res. (Å) | UniprotKB | Affinity type | Affinity (M) | Chain 1 | Chain 2 | Temp. (C) | pH |
|---|---|---|---|---|---|---|---|---|---|---|---|---|---|---|---|
| 2wy8 | PDBind | PF07678 | 21055811 | immune system | Homo sapiens Staphylococcus aureus | Yes | x-ray diffraction | 1.7 | P01024 | Kd | 3.6e-07 | 0 | 1 | 25.0 | 7.2 |
| 2x9a | BM5.5 PPI4DOC 3DComplexV6 | PF05357 PF06519 | 21110981 | viral protein | Escherichia phage If1 Escherichia coli | No | x-ray diffraction | 2.47 | O80297 Q8X965 | Kd | 4.4e-06 | D | C | 25.0 | 7.0 |
| 2xgy | PPI4DOC PDBind | PF00160 | 20833376 | viral protein / isomerase | Oryctolagus cuniculus Homo sapiens | No | x-ray diffraction | 1.8 | P62937 | Kd | 3e-05 | A | B | 25.0 | 7.2 |
| 2yvj | BM5.5 PPI4DOC | PF07992 PF14759 PF00355 | 17850818 | oxidoreductase / electron transport | Pseudomonas sp. | No | x-ray diffraction | 1.9 | Q52437 Q52440 | Kd | 0.000294 | A | B | 25.0 | 7.2 |
| 2z3q | PPI4DOC | PF02372 | 17643103 | cytokine / cytokine receptor | Homo sapiens | No | x-ray diffraction | 1.85 | P40933 Q13261 | Kd | 3.8e-11 | A | B | 25.0 | 7.2 |
| 2z58 | PDBind | PF00082 | 17988685 | hydrolase | Thermococcus kodakarensis KOD1 | Yes | x-ray diffraction | 1.88 | P58502 P58502 | Kd | 1.6e-08 | 0 | 1 | 25.0 | 7.2 |
| 2z7f | PPI4DOC | PF00089 PF00095 | 18421166 | hydrolase / hydrolase inhibitor | Homo sapiens | No | x-ray diffraction | 1.7 | P08246 P03973 | IC50 | 1.3e-08 | E | I | 25.0 | 7.2 |
| 3aaa | BM5.5 | PF01267 PF01115 PF12796 | 20625546 | protein binding | Gallus gallus Homo sapiens | No | x-ray diffraction | 2.2 | P13127 P14315 P58546 | Kd | 2.1e-08 | AB | C | 20.0 | 7.0 |
| 3ajb | PDBind | PF04882 | 21102411 | protein transport | Homo sapiens | Yes | x-ray diffraction | 2.5 | P56589 P40855 | Kd | 4.08e-08 | 0 | 1 | 25.0 | 7.2 |
| 3aon | PDBind | PF01813 PF01990 | 22114184 | hydrolase | Enterococcus hirae | No | x-ray diffraction | 2.0 | P43435 P43455 | Kd | 3.2e-09 | 0 | 1 | 25.0 | 7.2 |
| 3au4 | PDBind | PF00788 PF00373 PF00784 PF06583 | 21642953 | motor protein / apoptosis | Homo sapiens | No | x-ray diffraction | 1.9 | Q9HD67 P43146 | Kd | 5.3e-07 | 0 | 1 | 25.0 | 7.2 |
| 3blh | PDBind | PF00069 PF00134 PF21797 | 18566585 | transcription | Homo sapiens | Yes | x-ray diffraction | 2.48 | P50750 O60563 | Kd | 3e-07 | 0 | 1 | 25.0 | 7.2 |
| 3bx1 | PPI4DOC | PF00082 PF00197 | 18556023 | hydrolase / hydrolase inhibitor | Lederbergia lenta Hordeum vulgare | No | x-ray diffraction | 1.85 | P29600 P07596 | Kd | 1.3e-06 | A | B | 25.0 | 7.2 |
| 3c4p | PDBind | PF13354 PF00144 PF07467 | 18775544 | hydrolase / hydrolase inhibitor | Klebsiella pneumoniae Streptomyces clavuligerus | Yes | x-ray diffraction | 1.75 | P0AD64 P35804 | Kd | 4.4e-09 | 0 | 1 | 25.0 | 7.2 |
| 3c5t | PDBind | PF02793 PF00123 | 18287102 | signaling protein / signaling protein | Homo sapiens Heloderma suspectum | No | x-ray diffraction | 2.1 | P43220 P26349 | IC50 | 6e-10 | 0 | 1 | 25.0 | 7.2 |
| 3c9a | PPI4DOC | PF11581 | 18500331 | hormone / signaling protein | Drosophila melanogaster | No | x-ray diffraction | 1.6 | Q00805 Q01083 | Kd | 7.7e-09 | A | B | 25.0 | 7.2 |
| 3cqx | PPI4DOC | PF00012 | 19029896 | chaperone | Mus musculus | No | x-ray diffraction | 2.3 | P63017 Q91YN9 | Kd | 4.5e-06 | B | C | 25.0 | 7.2 |
| 3cx7 | PDBind | PF00503 PF09128 | 18940608 | signaling protein | Mus musculus Rattus norvegicus | No | x-ray diffraction | 2.25 | P27601 Q9ES67 | Kd | 5.2e-07 | 0 | 1 | 25.0 | 7.2 |

| PDB | Database origin | Protein family | PubMed | Classification | Organism | Mutation | Method | Res. (Å) | UniprotKB | Affinity type | Affinity (M) | Chain 1 | Chain 2 | Temp. (C) | pH |
|---|---|---|---|---|---|---|---|---|---|---|---|---|---|---|---|
| 3d2u | PPI4DOC | PF00129 PF07654 PF07654 PF13895 PF00022 | 18632577 | immune system | Human betaherpesvirus 5 Homo sapiens | Yes | x-ray diffraction | 2.21 | P08560 P61769 Q8NHL6 P60709 | Kd | 4e-09 | C | B | 25.0 | 7.2 |
| 3ddc | PDBind | PF00071 | 18596699 | hydrolase / apoptosis | Homo sapiens Mus musculus | Yes | x-ray diffraction | 1.8 | P01112 Q5EBH1 | Kd | 7.7e-07 | 0 | 1 | 25.0 | 7.2 |
| 3doe | PDBind | PF00025 PF11527 | 19368893 | signaling protein / hydrolase | Homo sapiens | No | x-ray diffraction | 2.25 | P36404 Q9Y2Y0 | Kd | 2e-08 | 0 | 1 | 25.0 | 7.2 |
| 3e1z | PPI4DOC PDBind | PF09394 PF00112 | 19143838 | hydrolase inhibitor / hydrolase | Trypanosoma cruzi Carica papaya | No | x-ray diffraction | 1.86 | Q966X9 P00784 | Ki | 3.6e-11 | A | B | 25.0 | 7.2 |
| 3fju | PDBind | PF00246 PF15270 | 19179285 | hydrolase / hydrolase inhibitor | Homo sapiens Ascaris suum | No | x-ray diffraction | 1.6 | P15085 P19399 | Ki | 1.6e-09 | 0 | 1 | 25.0 | 7.2 |
| 3fpu | PDBind | PF19429 PF00048 | 20041127 | immune system | Rhipicephalus sanguineus Homo sapiens | Yes | x-ray diffraction | 1.76 | P0C8E7 P10147 | Kd | 1.2e-10 | 0 | 1 | 25.0 | 7.2 |
| 3gc3 | 3DComplexV6 PDBind | PF00339 PF02752 PF09268 PF13838 PF01394 | 19710023 | endocytosis | Bos taurus | No | x-ray diffraction | 2.2 | P17870 P49951 | Kd | 2.1e-06 | 0 | 1 | 25.0 | 7.2 |
| 3gmw | PPI4DOC | PF13354 PF00144 PF07467 | 19332077 | protein binding | Escherichia sp. Sflu5 Streptomyces exfoliatus | No | x-ray diffraction | 2.1 | A5PHA6 Q9KJ90 | Kd | 5e-11 | A | B | 25.0 | 7.2 |
| 3gni | PPI4DOC PDBind | PF08569 PF00069 | 19513107 | signaling protein / signaling protein | Homo sapiens | No | x-ray diffraction | 2.35 | Q9Y376 Q7RTN6 | Kd | 1.2e-08 | A | B | 25.0 | 7.2 |
| 3gqi | PPI4DOC PDBind | PF07714 PF00017 | 19665973 | transferase / transferase inhibitor | Homo sapiens Rattus norvegicus | Yes | x-ray diffraction | 2.5 | P11362 P10686 | Kd | 3.3e-08 | A | B | 25.0 | 7.2 |
| 3gxu | PDBind | PF01404 PF00812 | 19875447 | transferase | Homo sapiens | No | x-ray diffraction | 2.5 | P54764 P52799 | Kd | 2.03e-07 | 0 | 1 | 25.0 | 7.2 |
| 3h42 | SabDab | PF05922 PF18463 PF18464 PF18459 | 19443683 | hydrolase / immune system | Homo sapiens | Yes | x-ray diffraction | 2.3 | Q8NBP7 Q8NBP7 | Kd | 4e-12 | HL | BA | 25.0 | 7.2 |
| 3h6s | PPI4DOC | PF00112 PF10467 | 19846555 | hydrolase / hydrolase inhibitor | Homo sapiens Clitocybe nebularis | Yes | x-ray diffraction | 2.22 | O60911 Q3Y9I6 | Ki | 8e-11 | A | B | 25.0 | 7.2 |
| 3hh2 | PPI4DOC | PF00019 PF09289 PF21333 PF07648 | 19644449 | signaling protein / cytokine | Mus musculus Homo sapiens | No | x-ray diffraction | 2.15 | O08689 P19883 | Kd | 1.23e-08 | A | B | 25.0 | 7.2 |
| 3ixe | PDBind | PF12796 PF00412 | 19963065 | signaling protein / signaling protein | Homo sapiens | No | x-ray diffraction | 1.9 | Q13418 Q7Z4I7 | IC50 | 2.3e-06 | 0 | 1 | 25.0 | 7.2 |
| 3k1r | PDBind | PF21219 PF00595 PF00536 | 20142502 | structural protein | Homo sapiens | Yes | x-ray diffraction | 2.3 | Q9Y6N9 Q495M9 | Kd | 1e-09 | 0 | 1 | 25.0 | 7.2 |
| 3k2m | PPI4DOC | PF00017 | 20357770 | signaling protein / protein binding | Homo sapiens | No | x-ray diffraction | 1.75 | P00519 | Kd | 7e-09 | A | B | 25.0 | 7.2 |

| PDB | Database origin | Protein family | PubMed | Classification | Organism | Mutation | Method | Res. (Å) | UniprotKB | Affinity type | Affinity (M) | Chain 1 | Chain 2 | Temp. (C) | pH |
|---|---|---|---|---|---|---|---|---|---|---|---|---|---|---|---|
| 3kr3 | PPI4DOC | PF00049 | 20515953 | immune system | Homo sapiens | No | x-ray diffraction | 2.2 | P01344 | Kd | 4.9e-11 | HL | D | 25.0 | 7.2 |
| 3kuc | PDBind | PF00071 PF02196 | 20361980 | gtp binding protein / transferase | Homo sapiens | Yes | x-ray diffraction | 1.92 | P62834 P04049 | Kd | 4.42e-07 | 0 | 1 | 25.0 | 7.2 |
| 3l5w | BM5.5 | PF03487 | 20226193 | immune system | Mus musculus Homo sapiens | No | x-ray diffraction | 2.0 | P35225 | Kd | 5.2e-11 | LH | I | 25.0 | 7.4 |
| 3l95 | PPI4DOC | PF06816 PF07684 PF00066 | 20393564 | immune system | Homo sapiens | No | x-ray diffraction | 2.19 | P46531 | Kd | 2.5e-09 | HL | X | 25.0 | 7.2 |
| 3l9j | PDBind | PF00059 PF00229 | 20179326 | immune system | Homo sapiens | No | x-ray diffraction | 2.1 | P05452 P01375 | Kd | 3.4e-10 | 0 | 1 | 25.0 | 7.2 |
| 3lev | PPI4DOC | PF00140 PF04542 | 20876137 | immune system | Thermus aquaticus Homo sapiens | Yes | x-ray diffraction | 2.5 | Q9EZJ8 | Kd | 6e-10 | HL | A | 25.0 | 7.2 |
| 3m63 | PDBind | PF10408 PF04564 PF00240 | 20427284 | ligase / protein binding | Saccharomyces cerevisiae | Yes | x-ray diffraction | 2.4 | P54860 P48510 | Kd | 1.75e-07 | 0 | 1 | 25.0 | 7.2 |
| 3mc0 | PPI4DOC | PF07686 PF01123 PF02876 | 21059660 | immune system | Mus musculus Staphylococcus aureus | Yes | x-ray diffraction | 2.0 | A2NTY6 D0EMB6 | Kd | 1.25e-07 | A | B | 25.0 | 7.2 |
| 3mxw | BM5.5 | PF01085 PF01079 | 20504762 | signaling protein | Homo sapiens | No | x-ray diffraction | 1.83 | Q15465 | Kd | 7e-10 | LH | A | 30.0 | 7.2 |
| 3mzg | PDBind | PF00103 PF09067 | 20889499 | hormone / hormone receptor | Homo sapiens | Yes | x-ray diffraction | 2.1 | P01236 P16471 | Kd | 0.00497 | 0 | 1 | 25.0 | 7.2 |
| 3nvn | PPI4DOC PDBind | PF01403 PF01437 | 20727575 | viral protein / signaling protein | Ectromelia virus Homo sapiens | No | x-ray diffraction | 2.26 | Q8JL80 O60486 | Kd | 9.4e-09 | A | B | 25.0 | 7.2 |
| 3o34 | PDBind | PF00439 PF00628 PF00125 | 21164480 | transcription / protein binding | Homo sapiens | No | x-ray diffraction | 1.9 | O15164 P68431 | Kd | 8.8e-06 | 0 | 1 | 25.0 | 7.2 |
| 3o5t | PDBind | PF03747 PF00543 | 22074780 | hydrolase / transcription | Azospirillum brasilense | No | x-ray diffraction | 2.09 | A7XNI2 P70731 | Kd | 1.5e-06 | 0 | 1 | 25.0 | 7.2 |
| 3oiq | PDBind | PF16853 | 20877309 | protein binding | Saccharomyces cerevisiae | No | x-ray diffraction | 2.4 | P32797 P13382 | Kd | 3.8e-06 | 0 | 1 | 25.0 | 7.2 |
| 3oky | PPI4DOC | PF01437 PF01403 PF17960 PF01437 PF01403 | 20877282 | signaling protein | Mus musculus | No | x-ray diffraction | 2.19 | P70207 O35464 | Kd | 1.3e-06 | A | B | 25.0 | 7.2 |
| 3orv | PPI4DOC | PF03150 PF02975 PF06433 | 20929212 | oxidoreductase / electron transport | Paracoccus denitrificans PD1222 | Yes | x-ray diffraction | 1.91 | Q51658 P22619 A1BB97 | Kd | 9.4e-06 | B | E | 25.0 | 7.2 |
| 3p9w | PPI4DOC | PF00341 | 23507309 | signaling protein / immune system | Homo sapiens | No | x-ray diffraction | 2.41 | P15692 | Kd | 1.6e-08 | A | B | 25.0 | 7.2 |
| 3qc8 | PDBind | PF02933 PF02359 PF00789 | 21739474 | protein binding | Homo sapiens | No | x-ray diffraction | 2.2 | P55072 Q9UNN5 | Kd | 1.12e-05 | 0 | 1 | 25.0 | 7.2 |

| PDB | Database origin | Protein family | PubMed | Classification | Organism | Mutation | Method | Res. (Å) | UniprotKB | Affinity type | Affinity (M) | Chain 1 | Chain 2 | Temp. (C) | pH |
|---|---|---|---|---|---|---|---|---|---|---|---|---|---|---|---|
| 3qht | PPI4DOC | PF00240 | 21518904 | de novo protein | Saccharomyces cerevisiae synthetic construct | No | x-ray diffraction | 2.4 | Q12306 | Kd | 8.2e-08 | A | B | 25.0 | 7.2 |
| 3rea | PPI4DOC | PF00469 PF00018 | 21625496 | protein binding | HIV-1 M:B ARV2 / SF2 Homo sapiens | Yes | x-ray diffraction | 2.0 | P03407 P08631 | Kd | 1.79e-07 | A | B | 25.0 | 7.2 |
| 3ro2 | PDBind | PF13181 PF13176 PF13374 PF13424 | 21816348 | protein binding | Mus musculus Homo sapiens | No | x-ray diffraction | 2.3 | Q8VDU0 Q14980 | Kd | 4.6e-08 | 0 | 1 | 25.0 | 7.2 |
| 3rt0 | PPI4DOC | PF00481 PF10604 | 21658606 | hydrolase / hydrolase inhibitor | Arabidopsis thaliana | Yes | x-ray diffraction | 2.11 | Q9CAJ0 Q8H1R0 | Kd | 1.2e-06 | A | B | 25.0 | 7.2 |
| 3sgb | PDBind | PF00089 PF00050 | 6414511 | complex(serine proteinase-inhibitor) | Streptomyces griseus Meleagris gallopavo | No | x-ray diffraction | 1.8 | P00777 P68390 | Kd | 1.79e-11 | 0 | 1 | 25.0 | 7.2 |
| 3tac | PDBind | PF00536 PF07647 PF00069 | 21855798 | transferase / protein binding | Homo sapiens | No | x-ray diffraction | 2.2 | O14936 O75334 | Kd | 5.5e-07 | 0 | 1 | 25.0 | 7.2 |
| 3tei | PDBind | PF00069 | 23047924 | transferase | Homo sapiens | Yes | x-ray diffraction | 2.4 | P28482 Q15418 | Kd | 5e-07 | 0 | 1 | 25.0 | 7.2 |
| 3tkl | PPI4DOC PDBind | PF00071 | 22416225 | protein transport / protein binding | Homo sapiens Legionella pneumophila str. Corby | No | x-ray diffraction | 2.18 | P62820 | Kd | 4.3e-07 | A | B | 25.0 | 7.2 |
| 3tnf | PDBind | PF00071 PF18641 | 22011575 | protein transport | Homo sapiens Legionella pneumophila subsp. pneumophila str. Philadelphia 1 | No | x-ray diffraction | 2.5 | P61006 Q5ZWZ3 | Kd | 1e-11 | 0 | 1 | 25.0 | 7.2 |
| 3tu3 | PDBind | PF01734 PF20848 PF20983 | 23166655 | toxin / toxin chaperone | Pseudomonas aeruginosa | No | x-ray diffraction | 1.92 | O66100 O34208 | Kd | 5.7e-08 | 0 | 1 | 25.0 | 7.2 |
| 3tz1 | PDBind | PF13499 PF00992 | 23096565 | contractile protein | Chlamys nipponensis akazara | No | x-ray diffraction | 1.8 | Q27428 Q7M3Y3 | Kd | 1.41e-07 | 0 | 1 | 25.0 | 7.2 |
| 3u43 | PDBind | PF01320 PF21431 | 22306467 | protein binding | Escherichia coli | No | x-ray diffraction | 1.72 | P04482 P04419 | Kd | 1e-15 | 0 | 1 | 25.0 | 7.2 |
| 3ul4 | PDBind | PF00963 PF00404 | 23118225 | cell adhesion / protein binding | Acetivibrio thermocellus ATCC 27405 | No | x-ray diffraction | 1.95 | Q06848 L7MTK2 | Kd | 1.14e-08 | 0 | 1 | 25.0 | 7.2 |
| 3uyp | PPI4DOC PDBind | PF02832 | 22285214 | immune system | Mus musculus dengue virus type 4 | No | x-ray diffraction | 2.0 | Q2YHF0 | Kd | 4.1e-06 | A | B | 25.0 | 7.2 |
| 3uzq | PPI4DOC PDBind | PF02832 | 22285214 | immune system | Mus musculus dengue virus type 1 | No | x-ray diffraction | 1.6 | P27909 | Kd | 8.2e-11 | A | B | 25.0 | 7.2 |
| 3uzv | PPI4DOC PDBind | PF00869 PF02832 | 22285214 | immune system | Dengue virus 2 Jamaica / 1409 / 1983 Mus musculus | No | x-ray diffraction | 2.1 | P07564 | Kd | 4.3e-10 | A | B | 25.0 | 7.2 |
| 3w8i | PDBind | PF20929 PF06840 PF20929 | 23665169 | protein binding / transferase | Homo sapiens | No | x-ray diffraction | 2.4 | Q9BUL8 Q9P289 | Kd | 2.15e-09 | 0 | 1 | 25.0 | 7.2 |

| PDB | Database origin | Protein family | PubMed | Classification | Organism | Mutation | Method | Res. (Å) | UniprotKB | Affinity type | Affinity (M) | Chain 1 | Chain 2 | Temp. (C) | pH |
|---|---|---|---|---|---|---|---|---|---|---|---|---|---|---|---|
| 3w9e | PPI4DOC | PF01537 | 24100313 | viral protein / immune system | Human herpesvirus 2 strain HG52 Homo sapiens | No | x-ray diffraction | 2.3 | Q69467 | IC50 | 3.6e-09 | A | B | 25.0 | 7.2 |
| 3wa5 | PDBind | | 24100309 | hydrolase | Pseudomonas aeruginosa PAO1 | No | x-ray diffraction | 1.9 | Q9HYC5 Q9HYC4 | Kd | 2.81e-08 | 0 | 1 | 25.0 | 7.2 |
| 3wdg | PDBind | PF03167 PF06106 | 24150946 | hydrolase / hydrolase inhibitor | Staphylococcus aureus subsp. aureus MRSA252 Staphylococcus aureus | No | x-ray diffraction | 2.2 | Q6GJ88 Q936H5 | Kd | 1.198e-09 | 0 | 1 | 25.0 | 7.2 |
| 3wqb | PDBind | PF00082 PF01483 PF18492 | 25784551 | hydrolase / chaperone | Aeromonas sobria | Yes | x-ray diffraction | 1.41 | Q9L5A4 W5JXD7 | Kd | 1.8e-09 | 0 | 1 | 25.0 | 7.2 |
| 3wwn | PPI4DOC PDBind | PF00696 PF21344 | 25392000 | metal binding protein / transferase | Thermus thermophilus HB27 | No | x-ray diffraction | 1.85 | O50147 Q9ZND7 | Kd | 6.62e-06 | A | B | 25.0 | 7.2 |
| 3zet | PDBind | PF00814 PF00814 | 23471679 | hydrolase | Salmonella enterica subsp. enterica serovar Typhimurium str. ST4 / 74 | No | x-ray diffraction | 2.31 | E8X8J1 E8XBD7 | Kd | 2.9e-07 | 0 | 1 | 25.0 | 7.2 |
| 3zkq | PPI4DOC PDBind | PF00026 | 23695257 | hydrolase / immune system | Homo sapiens Lama glama | No | x-ray diffraction | 1.51 | Q9Y5Z0 | Kd | 1.2e-09 | A | D | 25.0 | 7.2 |
| 3zo0 | PPI4DOC | PF07654 PF00622 PF13765 | 18420815 | immune system / ligase | Mus musculus | No | x-ray diffraction | 1.99 | P01863 Q62191 | Kd | 4.37e-07 | A | B | 25.0 | 7.2 |
| 3zu7 | PPI4DOC PDBind | PF00069 | 22843676 | transferase / de novo protein | Rattus norvegicus synthetic construct | No | x-ray diffraction | 1.97 | P63086 | Kd | 6.6e-09 | A | B | 25.0 | 7.2 |
| 3zwz | PDBind | PF02430 | 22737069 | immune system | Plasmodium falciparum 3D7 | No | x-ray diffraction | 2.1 | Q7KQK5 Q8IKV6 | Kd | 2.03e-08 | 0 | 1 | 25.0 | 7.2 |
| 4a1u | PDBind | PF02180 PF00452 PF08945 | 22040025 | apoptosis | Homo sapiens synthetic construct | No | x-ray diffraction | 1.54 | Q07817 O43521 | Ki | 5.4e-07 | 0 | 1 | 25.0 | 7.2 |
| 4a49 | PPI4DOC PDBind | PF00097 PF00179 | 22266821 | ligase | Homo sapiens | Yes | x-ray diffraction | 2.21 | P22681 P62837 | Kd | 4.2e-05 | A | B | 25.0 | 7.2 |
| 4an7 | PPI4DOC PDBind | PF00089 PF00197 | 23094997 | hydrolase / hydrolase inhibitor | Sus scrofa Tamarindus indica | No | x-ray diffraction | 2.23 | P00761 F4ZZG4 | Ki | 3.2e-09 | A | B | 25.0 | 7.2 |
| 4apx | PPI4DOC PDBind | PF00028 PF00028 PF18432 | 23135401 | cell adhesion | Mus musculus | No | x-ray diffraction | 1.65 | Q99PF4 Q99PJ1 | Kd | 2.9e-06 | A | B | 25.0 | 7.2 |
| 4awx | PDBind | PF09012 | 23024345 | metal transport | Klebsiella pneumoniae | No | x-ray diffraction | 2.3 | B5XTS6 | Kd | 5e-07 | 0 | 1 | 25.0 | 7.2 |
| 4ayi | PPI4DOC | PF00084 PF08794 PF20937 | 23133374 | immune system | Homo sapiens Neisseria meningitidis MC58 | Yes | x-ray diffraction | 2.31 | P08603 Q19KF7 | Kd | 2e-09 | A | D | 25.0 | 7.2 |

| PDB | Database origin | Protein family | PubMed | Classification | Organism | Mutation | Method | Res. (Å) | UniprotKB | Affinity type | Affinity (M) | Chain 1 | Chain 2 | Temp. (C) | pH |
|---|---|---|---|---|---|---|---|---|---|---|---|---|---|---|---|
| 4b1x | PDBind | PF02755 PF00022 | 23041370 | structural protein | Oryctolagus cuniculus Mus musculus | No | x-ray diffraction | 1.8 | P68135 Q2M3X8 | Kd | 4.34e-06 | 0 | 1 | 25.0 | 7.2 |
| 4b93 | PDBind | PF13774 PF12796 | 23104059 | exocytosis | Mus musculus Homo sapiens | No | x-ray diffraction | 2.0 | P70280 Q96NW4 | Kd | 2.3e-06 | 0 | 1 | 25.0 | 7.2 |
| 4bd9 | PDBind | PF00246 PF00014 | 23746805 | hydrolase / hydrolase inhibitor | Homo sapiens Sabellastarte magnifica | Yes | x-ray diffraction | 2.2 | Q9UI42 P84875 | Ki | 3.1e-08 | 0 | 1 | 25.0 | 7.2 |
| 4c2a | PDBind | PF00092 PF16164 PF13855 PF01462 | 24391089 | blood clotting | Homo sapiens | Yes | x-ray diffraction | 2.08 | P04275 P07359 | Kd | 9.7e-09 | 0 | 1 | 25.0 | 7.2 |
| 4c4p | PDBind | PF00071 PF09457 | 24056041 | protein transport | Homo sapiens | No | x-ray diffraction | 2.0 | P62491 Q7L804 | Kd | 2.5e-07 | 0 | 1 | 25.0 | 7.2 |
| 4c5g | PDBind | PF02820 | 24186981 | transcription | Drosophila melanogaster | No | x-ray diffraction | 2.1 | Q9VK33 Q8ST83 | Kd | 1.03e-06 | 0 | 1 | 25.0 | 7.2 |
| 4c7n | PDBind | PF11851 PF00010 | 24631970 | transcription | Homo sapiens synthetic construct | No | x-ray diffraction | 2.1 | O75030 | Kd | 1e-09 | 0 | 1 | 25.0 | 7.2 |
| 4c9b | PDBind | PF00270 PF00271 PF02854 | 24218557 | splicing | Homo sapiens | No | x-ray diffraction | 2.0 | P38919 Q9HCG8 | Kd | 2.84e-08 | 0 | 1 | 25.0 | 7.2 |
| 4cj0 | PPI4DOC PDBind | PF00759 PF02927 | 24823716 | hydrolase / de novo protein | Acetivibrio thermocellus synthetic construct | No | x-ray diffraction | 1.1 | P0C2S4 | Kd | 9.8e-08 | A | B | 25.0 | 7.2 |
| 4cmm | PPI4DOC PDBind | PF07686 PF08204 | 24550402 | signaling protein | Homo sapiens | Yes | x-ray diffraction | 1.92 | P78324 Q08722 | Kd | 8e-07 | A | B | 25.0 | 7.2 |
| 4ct0 | PDBind | PF00875 PF03441 PF12114 | 24855952 | circadian clock protein | Mus musculus | No | x-ray diffraction | 2.45 | P97784 O54943 | Kd | 2.8e-08 | 0 | 1 | 25.0 | 7.2 |
| 4d0g | PDBind | PF00071 PF09457 | 26032412 | hydrolase | Homo sapiens | Yes | x-ray diffraction | 2.5 | P61106 Q6WKZ4 | Kd | 1.8e-06 | 0 | 1 | 25.0 | 7.2 |
| 4d0n | PPI4DOC PDBind | PF00071 PF00169 PF00621 | 25186459 | cell cycle | Homo sapiens | No | x-ray diffraction | 2.1 | P61586 Q12802 | Kd | 1.81e-05 | A | B | 25.0 | 7.2 |
| 4drx | 3DComplexV[illegible] | PF00091 PF03953 PF00091 PF03953 | 22778434 | cell cycle | Ovis aries synthetic construct | No | x-ray diffraction | 2.22 | D0VWZ0 D0VWY9 | Kd | 1.55e-07 | AB | F | 25.0 | 7.2 |
| 4dt1 | PDBind | PF16834 PF16836 | 22465956 | dna binding protein | Saccharomyces cerevisiae S288C | No | x-ray diffraction | 1.9 | P40465 Q12318 | Kd | 5.6e-07 | 0 | 1 | 25.0 | 7.2 |
| 4dtg | PPI4DOC | PF00014 | 22563084 | blood clotting inhibitor / immune system | Homo sapiens | No | x-ray diffraction | 1.8 | P10646 | Kd | 7.3e-11 | HL | K | 25.0 | 7.2 |
| 4dxa | PDBind | PF00071 PF00373 | 22577140 | protein binding | Homo sapiens | Yes | x-ray diffraction | 1.95 | P61224 O00522 | Kd | 1.8e-06 | 0 | 1 | 25.0 | 7.2 |
| 4eig | PDBind | PF00186 | 23911607 | oxidoreductase / immune system | Escherichia coli K-12 Lama glama | No | x-ray diffraction | 2.5 | P0ABQ4 | Kd | 1e-09 | 0 | 1 | 25.0 | 7.2 |

| PDB | Database origin | Protein family | PubMed | Classification | Organism | Mutation | Method | Res. (Å) | UniprotKB | Affinity type | Affinity (M) | Chain 1 | Chain 2 | Temp. (C) | pH |
|---|---|---|---|---|---|---|---|---|---|---|---|---|---|---|---|
| 4eoz | PPI4DOC | PF00651 PF00888 | 22632832 | protein binding | Homo sapiens | Yes | x-ray diffraction | 2.4 | O43791 Q13618 | Kd | 1e-06 | A | D | 25.0 | 7.2 |
| 4etp | PDBind | PF00225 PF16796 | 22734002 | motor protein | Saccharomyces cerevisiae S288C | Yes | x-ray diffraction | 2.3 | P17119 Q12045 | Kd | 3.7e-07 | 0 | 1 | 25.0 | 7.2 |
| 4etw | PPI4DOC | PF00561 PF00550 | 23045647 | hydrolase | Shigella flexneri Shigella flexneri 5 str. 8401 | Yes | x-ray diffraction | 2.05 | Q83PW0 Q0T5U2 | Kd | 3.1e-06 | A | B | 25.0 | 7.2 |
| 4euk | PDBind | PF00072 PF01627 | 23132142 | signaling protein | Arabidopsis thaliana | No | x-ray diffraction | 1.95 | Q3S4A7 Q9ZNV9 | Kd | 4.1e-06 | 0 | 1 | 25.0 | 7.2 |
| 4fqi | BM5.5 | PF00509 PF00509 | 22878502 | viral protein / immune system | Influenza A virus (A / Viet Nam / 1203 / 2004(H5N1)) Homo sapiens | No | x-ray diffraction | 1.71 | Q6DQ33 Q6DQ33 | Kd | 9e-10 | HL | AB | 30.0 | 7.4 |
| 4fqj | PPI4DOC | PF00509 | 22878502 | viral protein / immune system | Influenza B virus (B / reassortant / NYMC BX-21A(Lee / 1940 x Florida / 04 / 2006)) Homo sapiens | No | x-ray diffraction | 2.5 | I0B7N4 | Kd | 9e-10 | HL | A | 25.0 | 7.2 |
| 4fzv | PDBind | PF01189 PF02536 | 23022348 | transferase | Homo sapiens | No | x-ray diffraction | 2.0 | Q96CB9 Q7Z6M4 | Kd | 1.33e-08 | 0 | 1 | 25.0 | 7.2 |
| 4g01 | PDBind | PF18151 PF02204 PF00071 | 23519409 | transport protein | Arabidopsis thaliana | No | x-ray diffraction | 2.2 | Q9LT31 Q9SN68 | Kd | 3.4e-06 | 0 | 1 | 25.0 | 7.2 |
| 4g35 | PDBind | PF00452 | 22920569 | apoptosis / inhibitor | Mus musculus | No | x-ray diffraction | 2.0 | P97287 | Ki | 5.4e-08 | 0 | 1 | 25.0 | 7.2 |
| 4g59 | PPI4DOC | PF14586 PF11624 | 23169621 | immune system | Mus musculus Murid betaherpesvirus 1 | No | x-ray diffraction | 2.44 | O08604 Q83156 | Kd | 4.2e-07 | A | B | 25.0 | 7.2 |
| 4g6u | PDBind | PF21111 PF21483 PF07262 | 23236156 | toxin | Escherichia coli O157:H7 | Yes | x-ray diffraction | 2.35 | F2WK69 F2WK70 | Kd | 1.78e-08 | 0 | 1 | 25.0 | 7.2 |
| 4gaf | PDBind | PF00340 PF18452 PF13895 | 23431173 | signaling protein | Homo sapiens | No | x-ray diffraction | 2.15 | P01584 P14778 | Kd | 1.8e-12 | 0 | 1 | 25.0 | 7.2 |
| 4gi3 | PPI4DOC PDBind | PF00082 | 23075397 | hydrolase / hydrolase inhibitor | Bacillus licheniformis Schistocerca gregaria | No | x-ray diffraction | 1.75 | Q9FDF2 P85064 | Ki | 6.8e-10 | A | C | 25.0 | 7.2 |
| 4h2w | PPI4DOC | PF00587 PF00550 | 23541895 | ligase | Bradyrhizobium diazoefficiens USDA 110 Agrobacterium fabrum str. C58 | No | x-ray diffraction | 1.95 | Q89VT8 A9CHM9 | Kd | 1.73e-06 | A | D | 25.0 | 7.2 |
| 4h5s | PPI4DOC PDBind | PF07686 PF07686 | 23871486 | cell adhesion | Homo sapiens | No | x-ray diffraction | 1.7 | O95727 Q9BY67 | Kd | 1.25e-05 | A | B | 25.0 | 7.2 |

| PDB | Database origin | Protein family | PubMed | Classification | Organism | Mutation | Method | Res. (Å) | UniprotKB | Affinity type | Affinity (M) | Chain 1 | Chain 2 | Temp. (C) | pH |
|---|---|---|---|---|---|---|---|---|---|---|---|---|---|---|---|
| 4h6j | PDBind | PF08447 PF14598 | 23033253 | transcription | Homo sapiens | Yes | x-ray diffraction | 1.52 | Q16665 P27540 | Kd | 1.9e-07 | 0 | 1 | 25.0 | 7.2 |
| 4hep | PDBind | PF18338 PF08931 | 23530214 | viral protein | Lactococcus phage TP901-1 Lama glama | No | x-ray diffraction | 1.75 | Q9G096 | Kd | 3e-07 | 0 | 1 | 25.0 | 7.2 |
| 4hff | PDBind | PF14113 PF16695 | 23288853 | hydrolase | Salmonella enterica subsp. enterica serovar Typhimurium str. LT2 | No | x-ray diffraction | 2.4 | Q93IS4 Q8ZRL5 | Kd | 2.69e-10 | 0 | 1 | 25.0 | 7.2 |
| 4hfk | PPI4DOC | PF14113 PF16695 | 23288853 | hydrolase | Enterobacter cloacae subsp. cloacae ATCC 13047 | No | x-ray diffraction | 2.1 | A0A0H3CIJ2 A0A0H3CIX8 | Kd | 2.69e-10 | C | D | 25.0 | 7.2 |
| 4hgm | PPI4DOC PDBind | PF00273 | 23632026 | immune system | Squalus acanthias Homo sapiens | No | x-ray diffraction | 2.34 | P02768 | Kd | 1.6e-08 | B | A | 25.0 | 7.2 |
| 4hjj | PPI4DOC | PF00340 | 23549062 | immune system | Homo sapiens | Yes | x-ray diffraction | 2.1 | Q14116 | Kd | 1.6e-10 | HL | A | 25.0 | 7.2 |
| 4i2x | PPI4DOC | PF07686 PF07654 | 23826770 | immune system | Homo sapiens | No | x-ray diffraction | 2.48 | Q9P1W8 | Kd | 1.2e-06 | C | B | 25.0 | 7.2 |
| 4i9x | PPI4DOC | PF16758 PF00020 | 23555243 | apoptosis | Human herpesvirus 5 strain Merlin Homo sapiens | No | x-ray diffraction | 2.1 | Q6RJQ3 O14763 | Kd | 5.96e-09 | B | D | 25.0 | 7.2 |
| 4ilw | PPI4DOC | PF00965 PF00413 | 24073280 | hydrolase / hydrolase inhibitor | Homo sapiens | No | x-ray diffraction | 2.1 | P16035 P09238 | Ki | 5.8e-09 | A | B | 25.0 | 7.2 |
| 4iu3 | PDBind | PF18244 | 23580648 | structural protein | Ruminococcus flavefaciens | No | x-ray diffraction | 1.97 | A0AEF6 A0AEF5 | Kd | 2.083e-08 | 0 | 1 | 25.0 | 7.2 |
| 4jeg | PPI4DOC PDBind | PF00017 PF00041 | 23980151 | signaling protein / protein binding | Homo sapiens | No | x-ray diffraction | 2.3 | Q06124 P02751 | Kd | 1.2e-08 | A | B | 25.0 | 7.2 |
| 4jeh | PPI4DOC PDBind | PF00995 PF05739 PF00804 | 23858467 | endocytosis / exocytosis | Rattus norvegicus | No | x-ray diffraction | 2.5 | P61765 P32851 | Kd | 8.1e-09 | A | B | 25.0 | 7.2 |
| 4jpk | PPI4DOC | PF00516 | 23539181 | immune system / viral protein | Homo sapiens Human immunodeficiency virus 1 | No | x-ray diffraction | 2.4 | P04578 | Kd | 4.4e-08 | HL | A | 25.0 | 7.2 |
| 4k2u | PPI4DOC | PF05424 | 23717209 | immune system | Plasmodium falciparum 3D7 Mus musculus | No | x-ray diffraction | 2.45 | Q8IBE8 | Kd | 1.53e-11 | A | B | 25.0 | 7.2 |
| 4k5a | PPI4DOC PDBind | PF02180 PF00452 | 24747052 | apoptosis | Bos taurus Escherichia coli | Yes | x-ray diffraction | 1.5 | Q1RMX3 | Kd | 1.03e-08 | A | B | 25.0 | 7.2 |
| 4k94 | PPI4DOC | | 24127596 | immune system | Homo sapiens | No | x-ray diffraction | 2.4 | P10721 | Kd | 6.3e-10 | HL | C | 25.0 | 7.2 |
| 4ka2 | PDBind | PF00516 | 23710622 | viral protein / inhibitor | Human immunodeficiency virus 1 | No | x-ray diffraction | 1.79 | P35961 | Kd | 8.4e-12 | 0 | 1 | 25.0 | 7.2 |

| PDB | Database origin | Protein family | PubMed | Classification | Organism | Mutation | Method | Res. (Å) | UniprotKB | Affinity type | Affinity (M) | Chain 1 | Chain 2 | Temp. (C) | pH |
|---|---|---|---|---|---|---|---|---|---|---|---|---|---|---|---|
| 4kgq | PPI4DOC | PF00020 PF00229 | 25087510 | immune system | Homo sapiens | Yes | x-ray diffraction | 2.27 | O95407 O43557 | Kd | 3.6e-08 | C | B | 25.0 | 7.2 |
| 4kt1 | PPI4DOC | PF13855 PF01462 PF15913 | 23756652 | hormone receptor / cell adhesion | Homo sapiens | No | x-ray diffraction | 2.5 | Q9BXB1 Q2MKA7 | Kd | 5.65e-08 | A | E | 25.0 | 7.2 |
| 4kt3 | PDBind | PF01832 PF21576 | 23878199 | hydrolase | Pseudomonas protegens Pf-5 | No | x-ray diffraction | 1.44 | Q4KC90 Q4KC91 | Kd | 2.6e-10 | 0 | 1 | 25.0 | 7.2 |
| 4l0p | PPI4DOC | PF01404 PF00812 | 25993310 | transferase / transferase receptor | Homo sapiens | No | x-ray diffraction | 2.26 | P29320 P52803 | Kd | 9e-09 | A | B | 25.0 | 7.2 |
| 4lad | PDBind | PF00179 PF13639 | 23942235 | ligase / ligase | Homo sapiens | No | x-ray diffraction | 2.3 | P60604 Q9UKV5 | Kd | 3.8e-08 | 0 | 1 | 25.0 | 7.2 |
| 4lgr | PPI4DOC PDBind | PF00161 | 24907552 | hydrolase / immune system | Ricinus communis Vicugna pacos | No | x-ray diffraction | 1.65 | P02879 | Kd | 1.1e-10 | A | B | 25.0 | 7.2 |
| 4llo | 3DComplexV6 | PF00027 PF13426 | 23975098 | transport protein | Mus musculus | No | x-ray diffraction | 2.0 | Q60603 Q60603 | Kd | 1.32e-05 | A | B | 25.0 | 7.2 |
| 4lxr | PPI4DOC | PF13855 PF01462 PF00560 PF01463 PF16077 | 24733933 | immune system / cytokine | Drosophila melanogaster | No | x-ray diffraction | 2.2 | P08953 P48607 | Kd | 5.9e-09 | A | JK | 25.0 | 7.2 |
| 4m0w | PPI4DOC PDBind | PF08715 PF00240 | 24531491 | hydrolase / protein binding | Severe acute respiratory syndrome-related coronavirus Bos taurus | Yes | x-ray diffraction | 1.4 | P0C6U8 P62992 | Kd | 6.7e-06 | A | B | 25.0 | 7.2 |
| 4m1g | PPI4DOC | PF05966 | 26325270 | immune system | Mus musculus Vaccinia virus | Yes | x-ray diffraction | 1.6 | Q71TT1 | Kd | 1.44e-08 | HL | AB | 25.0 | 7.2 |
| 4m5z | BM5.5 | PF00509 | 24027321 | viral protein / immune system | Influenza A virus Homo sapiens | No | x-ray diffraction | 2.25 | G8XMJ2 | Kd | 5e-09 | HL | A | 25.0 | 7.2 |
| 4m62 | PPI4DOC | PF07686 | 25254371 | immune system | Homo sapiens synthetic construct | No | x-ray diffraction | 1.8 | P01619 | Kd | 2.57e-09 | HL | S | 25.0 | 7.2 |
| 4mjs | PPI4DOC | PF00564 PF00564 | 24369353 | transferase / protein binding | Rattus norvegicus Homo sapiens | Yes | x-ray diffraction | 2.5 | P09217 Q13501 | Kd | 2.16e-08 | A | B | 25.0 | 7.2 |
| 4mrt | 3DComplexV6 PDBind | PF00550 PF01648 | 24704508 | transport protein / transferase | Brevibacillus parabrevis Bacillus subtilis subsp. subtilis str. 168 | Yes | x-ray diffraction | 2.0 | O30409 P39135 | Kd | 9.2e-07 | 0 | 1 | 25.0 | 7.2 |
| 4nl9 | PPI4DOC | PF00536 PF00536 | 24998259 | structural protein | Homo sapiens | Yes | x-ray diffraction | 1.5 | Q6ZW76 Q68DC2 | Kd | 2.49e-07 | A | B | 25.0 | 7.2 |
| 4nqw | PDBind | PF04545 PF10099 | 24699647 | dna binding protein / protein binding | Mycobacterium tuberculosis | No | x-ray diffraction | 2.4 | P9WGH7 P9WGX5 | Kd | 4.2e-07 | 0 | 1 | 25.0 | 7.2 |

| PDB | Database origin | Protein family | PubMed | Classification | Organism | Mutation | Method | Res. (Å) | UniprotKB | Affinity type | Affinity (M) | Chain 1 | Chain 2 | Temp. (C) | pH |
|---|---|---|---|---|---|---|---|---|---|---|---|---|---|---|---|
| 4nso | PDBind | PF21277 PF20889 | 24751834 | protein binding | Vibrio cholerae O1 biovar El Tor str. N16961 | No | x-ray diffraction | 2.4 | Q9KN42 Q9KN41 | Kd | 9.8e-10 | 0 | 1 | 25.0 | 7.2 |
| 4nu1 | PDBind | PF00069 | 24642411 | transferase / peptide | Mus musculus Homo sapiens | No | x-ray diffraction | 2.5 | Q9WV60 O15169 | Ki | 6e-05 | 0 | 1 | 25.0 | 7.2 |
| 4p3y | PPI4DOC PDBind | PF00009 PF03143 PF03144 PF01323 | 24860094 | translation / oxidoreductase | Escherichia coli BL21(DE3) Acinetobacter baumannii AYE | No | x-ray diffraction | 2.15 | P0CE47 | Kd | 1.46e-07 | A | B | 25.0 | 7.2 |
| 4pas | PDBind | PF18455 | 24778228 | signaling protein | Homo sapiens | No | x-ray diffraction | 1.62 | Q9UBS5 O75899 | Kd | 1.266e-07 | 0 | 1 | 25.0 | 7.2 |
| 4pbv | PPI4DOC | PF13855 PF16920 PF00047 PF13927 PF07679 | 25385546 | signaling protein | Gallus gallus | Yes | x-ray diffraction | 2.5 | Q91044 F1NWE3 | Kd | 2.4e-06 | A | B | 25.0 | 7.2 |
| 4pbz | PDBind | PF12265 PF00400 | 24920672 | cell cycle | Homo sapiens | No | x-ray diffraction | 2.15 | Q09028 Q13330 | Kd | 5e-08 | 0 | 1 | 25.0 | 7.2 |
| 4per | PPI4DOC PDBind | PF13516 PF18779 PF00074 | 24941155 | hydrolase / hydrolase inhibitor | Gallus gallus | No | x-ray diffraction | 1.92 | Q5ZIY8 P27043 | Kd | 1e-15 | A | B | 25.0 | 7.2 |
| 4pj2 | PPI4DOC | PF16743 PF05497 | 25664745 | hydrolase / hydrolase inhibitor | Aeromonas hydrophila subsp. hydrophila ATCC 7966 Meretrix lusoria | No | x-ray diffraction | 1.24 | A0KHJ5 P86383 | Kd | 4.7e-11 | A | B | 25.0 | 7.2 |
| 4pp8 | PPI4DOC | PF00059 PF14586 | 11825567 | immune system | Mus musculus | No | x-ray diffraction | 1.95 | O54709 O08603 | Kd | 4.86e-07 | A | C | 25.0 | 7.2 |
| 4pqt | PDBind | PF01398 | 24787148 | hydrolase / transcription | Schizosaccharomyces pombe 972h- Homo sapiens | Yes | x-ray diffraction | 2.05 | Q9P371 | Kd | 1.1e-06 | 0 | 1 | 25.0 | 7.2 |
| 4pw9 | PDBind | PF00174 PF03404 PF13442 | 26687009 | oxidoreductase / electron transport | Sinorhizobium meliloti 1021 | No | x-ray diffraction | 2.49 | Q92M24 Q92M25 | Kd | 1.35e-05 | 0 | 1 | 25.0 | 7.2 |
| 4qci | PPI4DOC | PF04692 PF00341 | 25707433 | cytokine / cytokine receptor | Homo sapiens | No | x-ray diffraction | 2.3 | P01127 | Kd | 2.8e-11 | A | C | 25.0 | 7.2 |
| 4qd2 | PPI4DOC 3DComplexV6 | PF14200 PF05588 PF03505 PF17993 PF00028 | 24948737 | cell adhesion | Clostridium botulinum A str. Hall Mus musculus | No | x-ray diffraction | 2.4 | A5HZZ6 A5HZZ5 A5HZZ4 P09803 | Kd | 2.7e-06 | D | E | 25.0 | 7.2 |
| 4qlp | PDBind | PF15598 PF14021 | 26237511 | hydrolase / protein binding | Mycobacterium tuberculosis H37Rv | No | x-ray diffraction | 1.1 | O05443 O05442 | Kd | 2.3e-10 | 0 | 1 | 25.0 | 7.2 |
| 4qxa | PDBind | PF00071 | 25220469 | protein transport / protein binding | Mus musculus | Yes | x-ray diffraction | 2.3 | Q9R0M6 Q8BPQ7 | Kd | 3.2e-07 | 0 | 1 | 25.0 | 7.2 |
| 4rey | PDBind | PF04495 PF19046 | 26363069 | membrane protein | Homo sapiens | No | x-ray diffraction | 1.96 | Q9BQQ3 Q08379 | Kd | 1.08e-07 | 0 | 1 | 25.0 | 7.2 |

| PDB | Database origin | Protein family | PubMed | Classification | Organism | Mutation | Method | Res. (Å) | UniprotKB | Affinity type | Affinity (M) | Chain 1 | Chain 2 | Temp. (C) | pH |
|---|---|---|---|---|---|---|---|---|---|---|---|---|---|---|---|
| 4tq1 | PDBind | PF20637 PF20638 PF04106 | 25484072 | protein binding | Homo sapiens | No | x-ray diffraction | 1.8 | Q9H1Y0 Q7Z6L1 | Kd | 0.00035 | 0 | 1 | 25.0 | 7.2 |
| 4u32 | PPI4DOC PDBind | PF00014 PF00089 | 25301953 | hydrolase / hydrolase inhibitor | Homo sapiens | Yes | x-ray diffraction | 1.65 | O43291 P35030 | Ki | 3.79e-07 | A | X | 25.0 | 7.2 |
| 4u4c | PDBind | PF08148 PF00270 PF00271 PF21408 PF13234 | 25175027 | hydrolase | Saccharomyces cerevisiae S288C | No | x-ray diffraction | 2.4 | P47047 P53632 Q12476 | Kd | 3.1e-07 | 0 | 1 | 25.0 | 7.2 |
| 4w4l | PPI4DOC | PF00934 PF00823 PF14011 | 25275011 | protein transport | Mycobacterium tuberculosis str. Erdman = ATCC 35801 | No | x-ray diffraction | 2.45 | A0A0H3LBR3 A0A0H3LBN6 A0A0H3LAM1 | Kd | 1.3e-09 | C | B | 25.0 | 7.2 |
| 4w6x | PDBind | | 25502211 | cell adhesion | Escherichia coli Lama glama | No | x-ray diffraction | 1.88 | Q47212 | Kd | 5.25e-09 | 0 | 1 | 25.0 | 7.2 |
| 4wem | PPI4DOC PDBind | PF07686 | 25828907 | structural protein | Escherichia coli Lama glama | No | x-ray diffraction | 1.55 | P14190 R9W2R6 | Kd | 6.8e-06 | A | B | 25.0 | 7.2 |
| 4wnd | PDBind | PF13181 PF13374 PF13176 | 25664792 | signaling protein / protein binding | Homo sapiens | No | x-ray diffraction | 1.5 | P81274 Q14CM0 | Kd | 3.29e-08 | 0 | 1 | 25.0 | 7.2 |
| 4x33 | PDBind | PF05207 PF13540 | 25604895 | electron transport | Saccharomyces cerevisiae S288C | No | x-ray diffraction | 1.45 | Q3E840 P31386 | Kd | 2.5e-07 | 0 | 1 | 25.0 | 7.2 |
| 4x7f | PPI4DOC | PF00915 PF08435 | 25520510 | viral protein | Norwalk virus Vicugna pacos | No | x-ray diffraction | 1.7 | Q5F4T5 | Kd | 2.8e-08 | A | C | 25.0 | 7.2 |
| 4x7s | PDBind | | 25849503 | immune system | Homo sapiens | No | x-ray diffraction | 1.9 | | Kd | 1e-08 | 0 | 1 | 25.0 | 7.2 |
| 4xl1 | PPI4DOC | PF00008 PF12661 PF07645 PF01414 PF07657 | 25700513 | protein binding | Rattus norvegicus | Yes | x-ray diffraction | 2.3 | Q07008 D3ZHH1 | Kd | 3.04e-07 | A | B | 25.0 | 7.2 |
| 4xwj | PDBind | PF04353 PF00381 | 26457424 | transcription / transferase | Escherichia coli K-12 | Yes | x-ray diffraction | 2.1 | P0AFX4 P0AA04 | Kd | 4.45e-09 | 0 | 1 | 25.0 | 7.2 |
| 4xxb | PDBind | PF00281 PF00673 PF00641 | 26220995 | rna binding protein / metal binding protein | Homo sapiens | No | x-ray diffraction | 2.4 | P62913 Q00987 | Kd | 1.48e-06 | 0 | 1 | 25.0 | 7.2 |
| 4y5o | PDBind | PF16545 PF00564 | 26235885 | transferase | Homo sapiens | No | x-ray diffraction | 2.35 | Q9BSQ5 Q99759 | Kd | 1.4e-06 | 0 | 1 | 25.0 | 7.2 |
| 4yl8 | PDBind | PF09379 PF09380 PF00373 | 25792740 | protein binding | Mus musculus Drosophila melanogaster | No | x-ray diffraction | 1.5 | P26041 P10040 | Kd | 5.4e-06 | 0 | 1 | 25.0 | 7.2 |
| 4yn0 | PPI4DOC PDBind | PF00020 PF12925 | 25838500 | apoptosis / cell adhesion | Mus musculus | No | x-ray diffraction | 2.2 | Q9EPU5 P12023 | Kd | 8.5e-08 | A | B | 25.0 | 7.2 |
| 4yvq | PDBind | PF00745 PF13424 | 26037924 | oxidoreductase / fluorescent protein | Arabidopsis thaliana | No | x-ray diffraction | 2.4 | P42804 Q940U6 | Kd | 2.03e-06 | 0 | 1 | 25.0 | 7.2 |

| PDB | Database origin | Protein family | PubMed | Classification | Organism | Mutation | Method | Res. (Å) | UniprotKB | Affinity type | Affinity (M) | Chain 1 | Chain 2 | Temp. (C) | pH |
|---|---|---|---|---|---|---|---|---|---|---|---|---|---|---|---|
| 4zii | PDBind | PF00452 PF06393 | 26158515 | apoptosis | Homo sapiens | Yes | x-ray diffraction | 2.19 | Q07812 P55957 | IC50 | 6.6e-08 | 0 | 1 | 25.0 | 7.2 |
| 4zqu | PDBind | PF21111 PF07262 | 26449640 | toxin | Yersinia pseudotuberculosis YPIII Salmonella enterica subsp. enterica serovar Rubislaw str. ATCC 10717 | No | x-ray diffraction | 2.09 | A0A0H3B0B8 A0A0R4I987 | Kd | 1.6e-08 | 0 | 1 | 25.0 | 7.2 |
| 4zw2 | PDBind | PF00625 PF12052 | 28351836 | metal transport | Mus musculus | Yes | x-ray diffraction | 1.86 | Q8R3Z5 Q02789 | Kd | 4.9e-09 | 0 | 1 | 25.0 | 7.2 |
| 5b76 | PDBind | PF00628 PF00125 | 27775714 | transferase | Homo sapiens | Yes | x-ray diffraction | 1.65 | Q92794 K7EMV3 | Kd | 5.8e-06 | 0 | 1 | 25.0 | 7.2 |
| 5cxb | PDBind | PF08154 PF00400 PF08145 PF00400 | 26476442 | protein binding | Thermochaetoides thermophila | No | x-ray diffraction | 2.1 | G0SFB5 G0SCK6 | Kd | 9.19e-09 | 0 | 1 | 25.0 | 7.2 |
| 5dfw | PDBind | PF00335 | 26637054 | cell adhesion | Homo sapiens Mus musculus | No | x-ray diffraction | 2.33 | P60033 | Kd | 5e-10 | 0 | 1 | 25.0 | 7.2 |
| 5djt | PDBind | PF13426 | 27427858 | signaling protein | Avena sativa Staphylococcus aureus | No | x-ray diffraction | 1.4 | O49004 | Kd | 1.7e-08 | 0 | 1 | 25.0 | 7.2 |
| 5dob | PDBind | PF02718 PF04541 | 26511021 | dna binding protein | Human herpesvirus 5 strain AD169 | Yes | x-ray diffraction | 2.47 | P16794 P16791 | Kd | 1e-06 | 0 | 1 | 25.0 | 7.2 |
| 5ee5 | PDBind | PF16213 PF00025 | 27373159 | transcription | Homo sapiens | No | x-ray diffraction | 2.28 | Q9Y6D6 P40616 | Kd | 2.6e-05 | 0 | 1 | 25.0 | 7.2 |
| 5elu | PDBind | PF11976 | 29138295 | signaling protein | synthetic construct Homo sapiens | No | x-ray diffraction | 2.35 | P61956 | Kd | 4.14e-07 | 0 | 1 | 25.0 | 7.2 |
| 5eo9 | PDBind | PF07686 PF13927 PF13927 PF00047 | 26687361 | cell adhesion | Drosophila melanogaster | No | x-ray diffraction | 2.3 | M9PC40 Q9W4R3 | Kd | 3.7e-07 | 0 | 1 | 25.0 | 7.2 |
| 5f4e | PDBind | PF16706 PF15005 PF03024 | 27309818 | cell adhesion | Homo sapiens | No | x-ray diffraction | 2.4 | Q8IYV9 A6ND01 | Kd | 4.8e-08 | 0 | 1 | 25.0 | 7.2 |
| 5f5s | PDBind | PF03371 PF06991 | 27773687 | splicing | Homo sapiens | No | x-ray diffraction | 2.4 | Q8NAV1 P55081 | Kd | 2.2e-08 | 0 | 1 | 25.0 | 7.2 |
| 5g1x | PDBind | PF00069 PF01056 | 27837025 | transferase | Homo sapiens | Yes | x-ray diffraction | 1.72 | O14965 P04198 | Kd | 1.21e-05 | 0 | 1 | 25.0 | 7.2 |
| 5gjk | PDBind | PF04433 PF04855 | 28438634 | transcription | Homo sapiens | No | x-ray diffraction | 2.05 | Q92922 Q12824 | Kd | 1.2e-07 | 0 | 1 | 25.0 | 7.2 |
| 5gpg | PDBind | PF00254 PF08771 | 27610411 | isomerase / transferase | Homo sapiens | No | x-ray diffraction | 1.67 | Q00688 P42345 | IC50 | 2.61e-09 | 0 | 1 | 25.0 | 7.2 |

| PDB | Database origin | Protein family | PubMed | Classification | Organism | Mutation | Method | Res. (Å) | UniprotKB | Affinity type | Affinity (M) | Chain 1 | Chain 2 | Temp. (C) | pH |
|---|---|---|---|---|---|---|---|---|---|---|---|---|---|---|---|
| 5h3j | PDBind | PF04495 | 28049725 | protein transport | Mus musculus | No | x-ray diffraction | 1.33 | Q99JX3 Q8R2X8 | Kd | 2.7e-07 | 0 | 1 | 25.0 | 7.2 |
| 5h7y | PDBind | PF18443 PF18426 | 28979890 | hydrolase inhibitor / peptide | Pseudomonas aeruginosa PAO1 | No | x-ray diffraction | 2.19 | Q9I3K3 Q9I3K2 | Kd | 1.25e-07 | 0 | 1 | 25.0 | 7.2 |
| 5hgg | BM5.5 | PF00089 | 27226628 | hydrolase / inhibitor | Homo sapiens Vicugna pacos | Yes | x-ray diffraction | 1.97 | P00749 | Kd | 5.4e-11 | T | A | 25.0 | 7.2 |
| 5hpk | PDBind | PF00632 PF00240 | 26949039 | ligase | Homo sapiens | No | x-ray diffraction | 2.43 | Q96PU5 P62987 | Kd | 9.7e-09 | 0 | 1 | 25.0 | 7.2 |
| 5hu3 | PDBind | PF00069 | 30381148 | transferase | Drosophila melanogaster | Yes | x-ray diffraction | 1.89 | Q00168 Q02280 | Kd | 1e-06 | 0 | 1 | 25.0 | 7.2 |
| 5hys | BM5.5 | PF07654 | 27194387 | immune system | Homo sapiens | Yes | x-ray diffraction | 2.5 | P01854 | Kd | 3.89e-11 | CD | JK | 25.0 | 7.2 |
| 5imk | PDBind | PF07686 | 27889311 | immune system | Camelidae Homo sapiens | No | x-ray diffraction | 1.23 | Q9Y279 | Kd | 8.5e-07 | 0 | 1 | 25.0 | 7.2 |
| 5imm | PDBind | PF07686 | 27889311 | immune system | Camelidae Mus musculus | No | x-ray diffraction | 1.2 | F6TUL9 | Kd | 3.5e-09 | 0 | 1 | 25.0 | 7.2 |
| 5inb | PDBind | PF00149 PF16891 PF15276 | 27572260 | hydrolase / protein binding | Homo sapiens | No | x-ray diffraction | 1.3 | P36873 Q69YH5 | Kd | 1.23e-07 | 0 | 1 | 25.0 | 7.2 |
| 5jds | PDBind | PF07686 | 28280600 | immune system | Camelidae Homo sapiens | No | x-ray diffraction | 1.7 | Q9NZQ7 | Kd | 3e-09 | 0 | 1 | 25.0 | 7.2 |
| 5jjd | PDBind | PF00169 PF01852 | 28652409 | lipid transport | Homo sapiens | No | x-ray diffraction | 2.4 | Q9Y5P4 Q9Y5P4 | Kd | 9.2e-06 | 0 | 1 | 25.0 | 7.2 |
| 5jw9 | PDBind | PF05110 PF07303 | 28134250 | protein binding | Homo sapiens | Yes | x-ray diffraction | 2.0 | Q9UHB7 O00472 | Kd | 8.6e-08 | 0 | 1 | 25.0 | 7.2 |
| 5kve | PDBind | PF02832 | 27475895 | viral protein / immune system | Zika virus Mus musculus | No | x-ray diffraction | 1.7 | A0A024B7W | Kd | 3.5e-08 | 0 | 1 | 25.0 | 7.2 |
| 5kxh | PDBind | PF10250 PF00008 | 28530709 | transferase | Mus musculus | No | x-ray diffraction | 1.33 | Q91ZW2 P70375 | Kd | 4.9e-07 | 0 | 1 | 25.0 | 7.2 |
| 5l21 | PDBind | PF07951 PF07953 | 28785006 | toxin | Clostridium botulinum A str. Hall Vicugna pacos | Yes | x-ray diffraction | 1.68 | P0DPI1 | Kd | 3.7e-09 | 0 | 1 | 25.0 | 7.2 |
| 5li1 | PDBind | PF00069 PF00433 | 27554858 | transferase | Homo sapiens Xenopus tropicalis | No | x-ray diffraction | 2.0 | P41743 Q28E03 | Kd | 4.7e-07 | 0 | 1 | 25.0 | 7.2 |
| 5ma6 | PDBind | PF01353 | 29176615 | fluorescent protein | Aequorea victoria synthetic construct | Yes | x-ray diffraction | 2.3 | P42212 | Kd | 3.03e-10 | 0 | 1 | 25.0 | 7.2 |
| 5me5 | PDBind | PF01652 | 28522457 | translation | Cucumis melo | No | x-ray diffraction | 1.9 | Q00LS8 A0A1S3C4H6 | Kd | 2.15e-06 | 0 | 1 | 25.0 | 7.2 |

| PDB | Database origin | Protein family | PubMed | Classification | Organism | Mutation | Method | Res. (Å) | UniprotKB | Affinity type | Affinity (M) | Chain 1 | Chain 2 | Temp. (C) | pH |
|---|---|---|---|---|---|---|---|---|---|---|---|---|---|---|---|
| 5ml9 | PDBind | PF13895 | 29247053 | immune system | Homo sapiens synthetic construct | Yes | x-ray diffraction | 2.35 | P08637 | Kd | 2.17e-07 | 0 | 1 | 25.0 | 7.2 |
| 5mtj | PDBind | PF00017 | 28347651 | signaling protein | Mus musculus Homo sapiens | No | x-ray diffraction | 1.95 | Q04736 | Kd | 3.38e-07 | 0 | 1 | 25.0 | 7.2 |
| 5mtm | PDBind | PF00017 | 28347651 | transferase | Homo sapiens Mus musculus | No | x-ray diffraction | 2.4 | P06239 | Kd | 7e-09 | 0 | 1 | 25.0 | 7.2 |
| 5nqg | PDBind | PF02430 | 28817634 | cell invasion | Plasmodium vivax Sal-1 Plasmodium vivax | Yes | x-ray diffraction | 2.15 | A5K4Z2 A5K3N8 | Kd | 5e-08 | 0 | 1 | 25.0 | 7.2 |
| 5nus | PDBind | PF03850 | 28977422 | transcription | Thermochaetoides thermophila DSM 1495 | No | x-ray diffraction | 2.2 | G0RXV8 G0RZE6 | Kd | 1.1e-08 | 0 | 1 | 25.0 | 7.2 |
| 5o90 | PDBind | PF00069 | 29229647 | transferase | Mus musculus Homo sapiens | Yes | x-ray diffraction | 2.49 | P47811 Q15750 | Kd | 1.1e-06 | 0 | 1 | 25.0 | 7.2 |
| 5oaq | PDBind | PF17725 | 28960584 | transcription | Homo sapiens | No | x-ray diffraction | 1.95 | Q15561 P46937 | Kd | 5.8e-08 | 0 | 1 | 25.0 | 7.2 |
| 5oyl | PDBind | PF00974 PF00057 | 29531262 | viral protein | Recombinant vesicular stomatitis Indiana virus rVSV-G / GFP Homo sapiens | No | x-ray diffraction | 2.25 | B7UCZ5 P01130 | Kd | 7.5e-06 | 0 | 1 | 25.0 | 7.2 |
| 5szh | PDBind | PF12130 PF00071 | 27552051 | endocytosis | Homo sapiens | No | x-ray diffraction | 2.3 | O94851 Q9H0U4 | Kd | 5.2e-06 | 0 | 1 | 25.0 | 7.2 |
| 5t0f | PDBind | PF14215 | 28137867 | transcription | Arabidopsis thaliana | No | x-ray diffraction | 2.4 | Q9FIP9 Q93ZM9 | IC50 | 5.8e-08 | 0 | 1 | 25.0 | 7.2 |
| 5tar | PDBind | PF00071 PF05351 | 27791178 | oncoprotein | Homo sapiens | No | x-ray diffraction | 1.9 | P01116 O43924 | Kd | 2.3e-06 | 0 | 1 | 25.0 | 7.2 |
| 5tvq | PDBind | PF03372 PF11976 | 28912134 | hydrolase | Mus musculus | No | x-ray diffraction | 2.35 | Q9JJX7 P61957 | Kd | 8.8e-07 | 0 | 1 | 25.0 | 7.2 |
| 5tzp | PDBind | PF00452 PF12201 | 28411240 | apoptosis | Fowlpox virus strain NVSL Gallus gallus | No | x-ray diffraction | 1.35 | Q9J5G4 | Kd | 3e-08 | 0 | 1 | 25.0 | 7.2 |
| 5un7 | PDBind | PF21375 | 28393830 | protein binding | Homo sapiens | No | x-ray diffraction | 2.1 | Q9NUX5 Q96AP0 | Kd | 1.2e-07 | 0 | 1 | 25.0 | 7.2 |
| 5uul | PDBind | PF00452 PF15826 | 28594323 | apoptosis | Homo sapiens | Yes | x-ray diffraction | 1.33 | Q16548 Q9BXH1 | Ki | 4.8e-09 | 0 | 1 | 25.0 | 7.2 |
| 5vko | PDBind | PF00017 | 28826505 | transcription | Saccharomyces cerevisiae S288C | No | x-ray diffraction | 1.8 | P23615 P04050 | Kd | 4e-09 | 0 | 1 | 25.0 | 7.2 |
| 5vmo | PDBind | PF00452 PF08945 | 29483196 | viral protein / apoptosis | Grouper iridovirus Danio rerio | No | x-ray diffraction | 1.7 | Q5GAF0 B8JK68 | Kd | 8.87e-07 | 0 | 1 | 25.0 | 7.2 |
| 5vz4 | PDBind | PF00019 PF02351 | 28953886 | signaling protein / protein binding | Homo sapiens | No | x-ray diffraction | 2.2 | Q99988 Q6UXV0 | Kd | 8e-09 | 0 | 1 | 25.0 | 7.2 |

| PDB | Database origin | Protein family | PubMed | Classification | Organism | Mutation | Method | Res. (Å) | UniprotKB | Affinity type | Affinity (M) | Chain 1 | Chain 2 | Temp. (C) | pH |
|---|---|---|---|---|---|---|---|---|---|---|---|---|---|---|---|
| 5w89 | PDBind | PF00452 | 29339518 | peptide binding protein | Homo sapiens | No | x-ray diffraction | 1.42 | Q07820 | IC50 | 2.5e-08 | 0 | 1 | 25.0 | 7.2 |
| 5wgg | PDBind | PF04055 PF13186 PF13353 PF13165 | 28704043 | peptide binding protein | Acetivibrio thermocellus ATCC 27405 | No | x-ray diffraction | 2.04 | A3DDW1 A3DDW2 | Kd | 7e-07 | 0 | 1 | 25.0 | 7.2 |
| 5wpa | PDBind | PF08075 PF00076 PF08075 PF00076 | 29530979 | nuclear protein | Homo sapiens | No | x-ray diffraction | 2.29 | P23246 Q8WXF1 | Kd | 5.5e-07 | 0 | 1 | 25.0 | 7.2 |
| 5wrv | PDBind | PF16969 PF17004 | 28369529 | protein transport | Homo sapiens | Yes | x-ray diffraction | 1.7 | Q9UHB9 O76094 | Kd | 7.3e-07 | 0 | 1 | 25.0 | 7.2 |
| 5wuj | PDBind | PF14842 | 29229777 | motor protein | Helicobacter pylori 26695 | No | x-ray diffraction | 2.3 | O25118 O25119 | Kd | 4.22e-07 | 0 | 1 | 25.0 | 7.2 |
| 5xiu | PDBind | PF00240 | 29330428 | transferase / ribosomal protein | Homo sapiens Mus musculus | No | x-ray diffraction | 1.8 | Q8IYW5 P62983 | Kd | 2.5e-05 | 0 | 1 | 25.0 | 7.2 |
| 5xln | PDBind | PF01652 | 30902983 | rna binding protein / ligase | Homo sapiens | No | x-ray diffraction | 1.9 | O60573 P26639 | Kd | 2.48e-06 | 0 | 1 | 25.0 | 7.2 |
| 5xoc | PDBind | PF03166 PF00085 | 29588413 | transcription | Homo sapiens Escherichia coli K-12 | No | x-ray diffraction | 2.4 | P84022 P0AA25 O75593 | Kd | 8e-06 | 0 | 1 | 25.0 | 7.2 |
| 5xod | PDBind | PF03166 | 29588413 | transcription | Homo sapiens | No | x-ray diffraction | 1.85 | Q15796 P12755 | Kd | 2.81e-08 | 0 | 1 | 25.0 | 7.2 |
| 5y9j | BM5.5 | PF00229 | 29572471 | protein binding | Homo sapiens | No | x-ray diffraction | 2.05 | Q9Y275 | Kd | 9.95e-10 | HL | A | 25.0 | 7.2 |
| 5yi8 | PDBind | PF00640 | 29467404 | cell cycle | Drosophila melanogaster | No | x-ray diffraction | 2.0 | P16554 Q9W4I7 | Kd | 1.8e-06 | 0 | 1 | 25.0 | 7.2 |
| 5yip | PDBind | PF02991 | 29867141 | signaling protein | Mus musculus Rattus norvegicus | No | x-ray diffraction | 1.85 | Q8R3R8 O70511 | Kd | 3.7e-09 | 0 | 1 | 25.0 | 7.2 |
| 5yr0 | PDBind | PF17675 PF10186 | 29866835 | endocytosis | Mus musculus | Yes | x-ray diffraction | 1.9 | O88597 Q8K245 | Kd | 7.6e-06 | 0 | 1 | 25.0 | 7.2 |
| 5ywr | PDBind | PF00179 PF13639 | 29626159 | signaling protein | Homo sapiens | No | x-ray diffraction | 1.47 | P61088 Q8ND25 | Kd | 3.9e-08 | 0 | 1 | 25.0 | 7.2 |
| 6a0z | BM5.5 | PF00509 | 29925655 | viral protein / immune system | Influenza A virus (A / chicken / Chachoengsao / Thailand / CU-11 / 04(H5N1)) Human immunodeficiency virus 1 Mus musculus | No | x-ray diffraction | 2.33 | Q6DQ34 M1E1E4 | Kd | 5.3e-08 | HL | A | 25.0 | 7.2 |
| 6aaf | PDBind | PF02991 | 30451685 | membrane protein | Schizosaccharomyces pombe 972h- | No | x-ray diffraction | 2.2 | O94272 Q09906 | Kd | 1.61e-07 | 0 | 1 | 25.0 | 7.2 |

| PDB | Database origin | Protein family | PubMed | Classification | Organism | Mutation | Method | Res. (Å) | UniprotKB | Affinity type | Affinity (M) | Chain 1 | Chain 2 | Temp. (C) | pH |
|---|---|---|---|---|---|---|---|---|---|---|---|---|---|---|---|
| 6akm | PDBind | PF05769 | 30622739 | protein binding | Homo sapiens | No | x-ray diffraction | 2.3 | Q9BRV8 Q14BN4 | Kd | 6.9e-06 | 0 | 1 | 25.0 | 7.2 |
| 6b0s | BM5.5 | PF00090 | 29167197 | immune system | Homo sapiens Plasmodium falciparum | No | x-ray diffraction | 1.95 | P19597 | Kd | 1.78e-07 | HL | C | 25.0 | 7.2 |
| 6b6u | PDBind | PF00224 PF02887 | 29182273 | transferase | Homo sapiens | Yes | x-ray diffraction | 1.35 | P14618 | Kd | 2.01e-05 | 0 | 1 | 25.0 | 7.2 |
| 6bmt | PDBind | PF03098 | 29524428 | oxidoreductase / inhibitor | Homo sapiens Staphylococcus delphini | No | x-ray diffraction | 2.4 | P05164 A0A2A4GXB5 | Kd | 3.1e-07 | 0 | 1 | 25.0 | 7.2 |
| 6bw9 | PDBind | PF00514 PF01749 PF16186 | 30209309 | transport protein | Homo sapiens Hendra virus horse / Australia / Hendra / 1994 | No | x-ray diffraction | 1.6 | O00629 P0C1C6 | Kd | 4.39e-09 | 0 | 1 | 25.0 | 7.2 |
| 6d4p | PDBind | PF00179 PF00240 | 31634471 | transferase | Homo sapiens | Yes | x-ray diffraction | 2.11 | P51668 Q59EM9 | IC50 | 6.5e-08 | 0 | 1 | 25.0 | 7.2 |
| 6dgf | PDBind | PF00443 PF00240 | 30763569 | protein binding | Homo sapiens | Yes | x-ray diffraction | 2.34 | O75604 P0CG47 | IC50 | 9.5e-09 | 0 | 1 | 25.0 | 7.2 |
| 6f0f | PDBind | PF04729 | 31543461 | chaperone | Homo sapiens | No | x-ray diffraction | 2.0 | Q9Y294 | Kd | 1.8e-07 | 0 | 1 | 25.0 | 7.2 |
| 6fbx | PDBind | PF00452 PF10514 | 30237469 | apoptosis | Danio rerio | No | x-ray diffraction | 1.64 | Q8UWD5 Q4V925 | Kd | 3.43e-07 | 0 | 1 | 25.0 | 7.2 |
| 6fc3 | PDBind | PF01652 PF17052 | 30053226 | translation | Saccharomyces cerevisiae S288C | Yes | x-ray diffraction | 1.75 | P07260 P12962 | Kd | 2e-08 | 0 | 1 | 25.0 | 7.2 |
| 6fp7 | PDBind | PF01353 | 30237292 | de novo protein | Clavularia sp. synthetic construct | Yes | x-ray diffraction | 1.58 | Q9U6Y3 | Kd | 3e-09 | 0 | 1 | 25.0 | 7.2 |
| 6fub | PDBind | | 29988155 | antifungal protein | Pyricularia oryzae Oryza sativa Japonica Group | No | x-ray diffraction | 1.3 | C4B8C2 B5UBC1 | Kd | 2.9e-08 | 0 | 1 | 25.0 | 7.2 |
| 6fv0 | PDBind | PF21376 PF00515 PF13374 PF13424 | 30320553 | motor protein | Mus musculus Lama glama | No | x-ray diffraction | 2.29 | O88447 Q9ER39 | Kd | 1.72e-06 | 0 | 1 | 25.0 | 7.2 |
| 6gho | PDBind | PF03960 PF13743 | 30982633 | protein binding | Bacillus subtilis subsp. subtilis str. 168 Geobacillus kaustophilus HTA426 | No | x-ray diffraction | 1.79 | O31602 Q5L1S1 | Kd | 8e-07 | 0 | 1 | 25.0 | 7.2 |
| 6gum | PDBind | PF00179 | 31292170 | transferase | Arabidopsis thaliana | No | x-ray diffraction | 1.79 | Q42551 Q9SJT1 | Kd | 1.2e-06 | 0 | 1 | 25.0 | 7.2 |
| 6har | PDBind | PF00089 PF00014 | 30700553 | protein fibril | Homo sapiens | Yes | x-ray diffraction | 1.5 | Q8N2U3 H7C0V9 | Ki | 6.1e-11 | 0 | 1 | 25.0 | 7.2 |

| PDB | Database origin | Protein family | PubMed | Classification | Organism | Mutation | Method | Res. (Å) | UniprotKB | Affinity type | Affinity (M) | Chain 1 | Chain 2 | Temp. (C) | pH |
|---|---|---|---|---|---|---|---|---|---|---|---|---|---|---|---|
| 6her | PDBind | PF00377 | 31815959 | protein binding | Mus musculus Camelus dromedarius | No | x-ray diffraction | 1.2 | P04925 | Kd | 4e-08 | 0 | 1 | 25.0 | 7.2 |
| 6idx | PDBind | PF11841 PF04727 | 30604775 | cell adhesion | Homo sapiens Mus musculus | No | x-ray diffraction | 1.7 | Q96JJ3 Q3UHD1 | Kd | 3.36e-06 | 0 | 1 | 25.0 | 7.2 |
| 6imf | PDBind | PF08562 PF00188 PF05825 | 30504218 | toxin / antitoxin | Protobothrops flavoviridis | No | x-ray diffraction | 2.3 | Q8JI39 A7VN14 | Kd | 2.4e-08 | 0 | 1 | 25.0 | 7.2 |
| 6isc | PDBind | PF07686 PF07686 | 30591568 | immune system | Mus musculus Homo sapiens | No | x-ray diffraction | 2.2 | Q8K4F0 P15151 | Kd | 2.4e-06 | 0 | 1 | 25.0 | 7.2 |
| 6iwd | PDBind | PF00102 PF00527 | 31323018 | oncoprotein | Homo sapiens human papillomavirus 18 | No | x-ray diffraction | 1.8 | Q15678 P06788 | Kd | 1.82e-08 | 0 | 1 | 25.0 | 7.2 |
| 6j4o | PDBind | PF14822 PF15674 | 31235911 | hydrolase | Homo sapiens | No | x-ray diffraction | 2.3 | Q86V25 Q8N300 | Kd | 1.8e-08 | 0 | 1 | 25.0 | 7.2 |
| 6jwj | PDBind | PF05020 PF05021 | 31836717 | protein binding | Saccharomyces cerevisiae S288C | Yes | x-ray diffraction | 1.58 | P33755 P53044 | Kd | 8.57e-08 | 0 | 1 | 25.0 | 7.2 |
| 6kbr | PDBind | PF00089 PF00050 | 31391482 | hydrolase / hydrolase inhibitor | Homo sapiens | No | x-ray diffraction | 2.0 | Q9Y5K2 P20155 | Kd | 8.91e-12 | 0 | 1 | 25.0 | 7.2 |
| 6mav | PDBind | PF00413 PF00965 | 31040180 | hydrolase / hydrolase inhibitor | Homo sapiens | Yes | x-ray diffraction | 2.37 | P08254 P01033 | Ki | 3.35e-11 | 0 | 1 | 25.0 | 7.2 |
| 6ne4 | PDBind | PF01392 | 31086346 | biosynthetic protein / signaling protein | Escherichia coli Homo sapiens | No | x-ray diffraction | 1.65 | O75084 | Kd | 1.6e-09 | 0 | 1 | 25.0 | 7.2 |
| 6pnp | PDBind | PF02210 PF06312 | 31566781 | cell adhesion | Mus musculus Rattus norvegicus | Yes | x-ray diffraction | 1.94 | Q9CS84 Q63366 | Kd | 2.992e-07 | 0 | 1 | 25.0 | 7.2 |
| 6r2g | PDBind | PF00517 PF00517 | 31255705 | biosynthetic protein | Human immunodeficiency virus 1 Human immunodeficiency virus type 1 (BRU ISOLATE) | Yes | x-ray diffraction | 1.9 | P03377 P03377 | Kd | 7.58e-13 | 0 | 1 | 25.0 | 7.2 |
| 6umt | PDBind | PF07686 | 31727844 | immune system | Homo sapiens | Yes | x-ray diffraction | 1.99 | Q15116 Q9BQ51 | Kd | 2.6e-09 | 0 | 1 | 25.0 | 7.2 |
